# KDM6A knockout in human iPSCs alters the genome-wide histone methylation profile at active and poised enhancers, activating expression of ectoderm gene expression pathways

**DOI:** 10.1101/2021.03.09.434633

**Authors:** Shailendra S. Maurya, Wei Yang, Qiang Zhang, Petra Erdmann-Gilmore, Amelia Bystry, Reid Townsend, Todd E. Druley

## Abstract

KDM6A is a histone demethylase, known to remove methyl moieties at the lysine residues of histone 3-labeled (H3K27me3) poised enhancers and bivalent promoters, which regulates gene expression during the differentiation of embryonic stem cells and tissue-specific development. However, while tissue- and disease-specific analyses have been performed, little is known about the location and consequences on gene expression of these regulatory regions in human pluripotent cells. Poised enhancers and bivalent promoters function in a coordinated fashion during development, which requires timely and efficient histone modifications. Identification of KDM6A-specific gene-regulatory domains is important for understanding the developmental mechanisms controlled by these histone modifications in pluripotency. In this study, we compared genome-wide histone modification and gene expression differences in isogenic wild type and cas9-mediated *KDM6A* knockout human induced pluripotent stem cells (hiPSC) lines. Here, we report that the absence of KDM6A does not alter the pluripotent phenotype but does substantially alter the histone modification profile at poised and active enhancers, resulting in decreased expression of associated COMPASS complex genes KMT2C and KMT2D and subsequently increasing the expression of gene pathways involved in ectoderm differentiation.

## Background

The lysine-specific demethylase 6A (KDM6A, also known as UTX) acts as an essential component of the COMPASS complexes anchored by KMT2C and KMT2D, which function to regulate histone modifications that control gene expression [1,2]. KDM6A demethylates H3K27me2/3 at promoters and enhancers [3]. H3K27me3 is associated with transcriptional regulation of important developmentally-related lineage genes and maintains them at a poised state in stem cells, as demonstrated by genome-wide mapping studies [4–7]. Given its repressive nature, H3K27 must be marked by trimethylation (H3K27me3) to close chromatin at loci unrelated to a particular cell fate and then removed to open chromatin in order to facilitate gene transcription necessary for the desired developmental program. This close/open mechanism was observed during the conversion of mouse embryonic fibroblasts to induced pluripotent stem cells (iPSCs) [8,9], suggesting similar mechanisms may impact other mammalian cell fate decision processes.

KDM6A is important for the differentiation of embryonic stem cells (ESCs) and the development of various tissues [10], both dependent and independent of its demethylase activity. KDM6A is required for ectoderm and mesoderm differentiation *in vitro*, independent of its catalytic activity [11]. KDM6A contributes to a variety of tissue-specific and developmental processes, including cardiac development, hematopoiesis, myogenesis, osteogenic differentiation, wound healing, and aging [12–17]. Functional studies have shown that germline *KDM6A* homozygous mutations cause severe midgestational defects, developmental delays, neural tube closure, yolk sack, and heart defects [12,13,16,18–20]. *KDM6A*KO mouse embryonic stem cells fail to induce Brachyury, a transcription factor essential for mesoderm development, resulting in severe mesodermal defects [21]. During ESC differentiation, the reactivation of developmental genes is associated with loss of H3K27me3 [7], which is a critical function of KDM6A when resolving promoter bivalency. Loss of KDM6A does not influence ESC self-renewal and proliferation but seems to provoke an effect on the differentiation capacity of ESCs [11–13,22]. This differentiation defect is due to a loss of expression of specific developmental genes thought essential for all three germ layers, presumably due to loss of KDM6A binding to the promoter region of these genes [11,21]. Finally, deleterious *KDM6A* mutations are found in a variety of cancer types [23].

KDM6A is a unique part of the KMT2C and KMT2D COMPASS complexes [1,24,25]. We previously found that infants with leukemia have an abundance of non-synonymous, predicted deleterious germline variation in KMT2C, a methyltransferase [26] and KDM6A (unpublished data). Functionally, deletion of *KMT2C* in human pluripotent stem cells (hiPSCs) cells resulted in genome-wide epigenomic reprogramming leading to a failure to specify hemogenic endothelium *in vitro* (manuscript in review). Therefore, we hypothesized that deletion of *KDM6A* in hiPSCs would result in a similar genome-wide changes in histone modification, chromatin state and subsequent gene expression. We found that *KDM6A*KO hiPSCs, compared to isogenic controls, demonstrate genome-wide histone modification differences primarily at bivalent promoters and poised enhancers, consistent with KDM6A mediating H3K27me3 repressive marks. In contrast, active enhancers were mostly unaffected, suggesting that KDM6A loss has profound effects on chromatin status early in differentiation, when fate commitment is not decided. To our knowledge, no comparative study exists describing the role of KDM6A and its demethylase activity in hiPSCs.

## Methods

### Wild type and isogenic *KDM6A*KO hiPSCs

Reprogrammed hiPSCs were generated from white blood cells collected from a healthy human male by the Washington University Genome Engineering and iPSC Core (GEiC; http://geic.wustl.edu/). From this control line, the GEiC generated an isogenic, bi-allelic *KDM6A*KO line via CRISPR-guided non-homologous end-joining via guide RNA (GGCATCCTGAGGCTGGTTGCNGG) targeting the gene locus on the X-chromosome resulting in a truncation of exons 4-29 (of 29 total exons) and validated by karyotyping and sequencing. hiPSCs were cultured in a feeder-free cell culture medium with established protocols provided by the manufacturer (STEMCELL #85850).

### Immunofluorescence staining

For staining, hiPSCs cells were grown on Matrigel in a chamber slide (Millipore sigma Z734535-96EA). Next, hiPSCs were fixed with 4% PFA (5 minutes at room temperature) then washed three times with PBS containing 0.1% BSA. Fixed hiPSCs were permeabilized (0.1% Triton X-100 in PBS) for 45 minutes followed by incubation in blocking solution (20% FCS, 0.1% Triton X-100 in PBS) for 30 minutes. Subsequently, samples were incubated overnight at 4°C with primary antibodies diluted in blocking solution. Samples were washed three times with PBS + 0.1% BSA before incubation (60 minutes, room temperature) with secondary antibodies diluted in blocking solution. After incubation, cells were washed three times with PBS. DAPI was added to the last washing step followed by mounting onto slides. Microscopy-based inspection and imaging of samples was performed on an inverted confocal laser scanning microscope.

### Verification of WT and KDM6A hiPSCs pluripotency

WT and *KDM6A*KO iPSCs cells were differentiated to ectoderm, mesoderm and endoderm using the media supplement provided by the R&D system kit (#SC027B), according to manufactures protocol. To further evaluate lineage commitment, cells were stained with goat anti-human OTX2 antibody, goat anti-human Brachyury, and goat anti-human SOX17 antibody provided with kit (#SC027B) as described by the manufacturer’s protocol.

### ChIPmentation

ChIPmentation was carried out as previously described [27] with minor modifications. Briefly, cells were washed once with PBS followed by fixation using 1% formaldehyde in up to 1 ml PBS for 10 min at room temperature. Glycine was used to stop the reaction. Cells were collected at 500g for 10 min at 4°C (subsequent work was performed in a 4°C cold room using ice-cold buffers unless otherwise specified) and washed once with 150 ul ice-cold PBS supplemented with protease inhibitors (Thermo Scientific #A32955). After that, fixed cells were left at −80°C for future experiments. Next, fixed frozen cells were lysed in sonication buffer supplemented with a protease inhibitor, as described, and then sonicated in a Covaris microtube (AFA fiber crimp-cap) with a Covaris E220 sonicator with the following settings: Peak incident power: 200, Duty factor: 10%, Cycles per burst: 200, Treatment time: 150 seconds, until the DNA fragments size, was in the range of 250-700bp. Following sonication, an equilibration buffer was added to the lysate. Lysates were centrifuged at 14000 RPM for 4°C and 10 minutes. The supernatant containing the sonicated chromatin was transferred into a new 1.5 ml DNA LoBind Eppendorf tube for immunoprecipitation. For each immunoprecipitation, 20ul magnetic DynabeadTM Protein A (Life Technologies) were washed twice and re-suspended in 2X PBS supplemented with 0.1% BSA. Then, 1μg of the appropriate antibody (described below) was added and bound to beads by rotating at least 6 hours at 4°C. Blocked antibody and conjugated beads were then placed on a magnetic separator (DYNAL Invitrogen bead separations from Invitrogen). The supernatant was aspirated, and the sonicated lysate was added to the beads followed by overnight incubation for 4°C on a rotator. Beads were washed as described in the original protocol subsequently at 4°C with various buffers as provided in the protocol. Beads were then re-suspended in the tagmentation mix (19 μl tagmentation buffer + 1μl Tagment DNA enzyme supplemented with 5μl nuclease-free water) from the Nextera DNA Sample Prep kit (Illumina) and incubated at 37°C for 10 minutes in a thermocycler. The beads were washed with an appropriate buffer (150 μl) and then transferred into a new 1.5 mL microfuge tube. The supernatant was immediately aspirated, leaving beads attached to the tube’s wall on a magnetic separator. Bead pellets were then resuspended with 45μl elution buffer supplemented with proteinase K (NEB) and incubated for 1 hour at 55°C and then 8-10 hours at 65°C to revert formaldehyde cross-linking. The resulting supernatant was transferred to a new tube, and the beads were then discarded. Finally, DNA was purified via MinElute kit by Qiagen. qPCR was performed from this purified DNA described in the protocol to estimate the optimum number of enrichment cycles. The libraries’ final enrichment was then performed according to protocol and subsequently purified using AMPure XP beads followed by a size selection to recover libraries with a fragment length of 250-400bp before sequencing.

### Antibodies used in ChiPmentation

ChIP antibodies were purchased from Diagenode: H3K4me3 (#C15410003), H3K4me1 (#C15410037), H3K27ac (#C15410174), H3K27me3 (#C15410069), Rabbit IgG (#C15410206), Embryonic Stem Cell Marker Panel (Human: OCT4, NANOG, SOX2, SSEA4) (ab109884).

### RNA sequencing

Cells were cultured to 70% confluency and then washed once with PBS, trypsinized, and pelleted by centrifugation at 500g for 10 min at 4°C. Cell pellets were transferred to the Genome Technology Access Center at Washington University for mRNA selection, Illumina sequencing library preparation, and sequencing on the NextSeq500 platform.

### ChIPmentation analysis

Biological replicates were prepared for each histone modification - H3K4me1, H3K4me3, H3K27ac, and H3K27me3 - in WT and *KDM6A*KO hiPSC along with two replicates of rabbit IgG as a negative control. Raw sequence reads were processed using the ENCODE Transcription factor and Histone Chip-Seq processing pipeline (2020a (http://github.com/ENCODE-DCC/chip-seq-pipeline2), assessed on Feb 27, 2019). The pipeline filtered and mapped the reads to hg19, validated the data’s quality, and generated fold-change signal tracks over the control samples using MACS2. Peaks were further analyzed using epic2 [28] using a false discovery rate (FDR) of 0.05, enabling both broad and narrow histone mark peaks to be efficiently identified. Transcription factor binding motif searches upstream and downstream of enhancer and bivalent promoter histone modification profiles (see below) was conducted using Homer v4.8.3 (http://homer.ucsd.edu/homer/index.html). Identifying an enrichment of differential peak-associated genes called Gene Ontology (http://geneontology.org) was performed using the Bioconductor R package cluster profile v3.12.0 [29].

### RNA-seq analysis

RNA-seq reads were aligned to the Ensembl release 72 primary assemblies with STAR version 2.5.1a. [30]. Gene counts were derived from the number of uniquely aligned unambiguous reads by Subread: feature count version 1.4.6-p5 [31]. All gene counts were then imported into the R/Bioconductor package EdgeR [32], and TMM normalization size factors were calculated to adjust for differences in library size. Ribosomal genes and genes not expressed in the smallest group size minus one sample greater than one count-per-million were excluded from further analysis. The TMM size factors and the matrix of counts were then imported into the R/Bioconductor package Limma [33]. Weighted likelihoods based on the observed mean-variance relationship of every gene and sample were then calculated for all samples with the voomWithQualityWeights [34]. All genes’ performance was assessed with plots of the residual standard deviation of every gene to their average log-count with a robustly fitted trend line of the residuals. Differential expression analysis was then performed to analyze for differences between conditions, and the results were filtered for only those genes with Benjamini-Hochberg FDR adjusted p-values £0.05. All data are available at GEO using accession number: GSE168252.

### Definition of enhancers and promoters

As listed in Table 1, promoters were defined as non-overlapping −1kb and +1kb intervals around transcription start sites (TSS). Enhancers were defined by H3K4me1 peaks and were assigned to their closest promoter, allowing for a maximum distance of 500 kb. Active enhancers were those overlapped with H3K27ac peaks. Poised and primed enhancers were assigned to promoters after excluding those associated with any active enhancers. Poised enhancers overlapped with H3K27me3, whereas primed enhancers did not. Promoters were defined by H3K4me3 peaks within 1kb of TSS. Bivalent promoters were determined by the overlapping peak of H3K4me3 and H3K27me3.

**Table 1:**
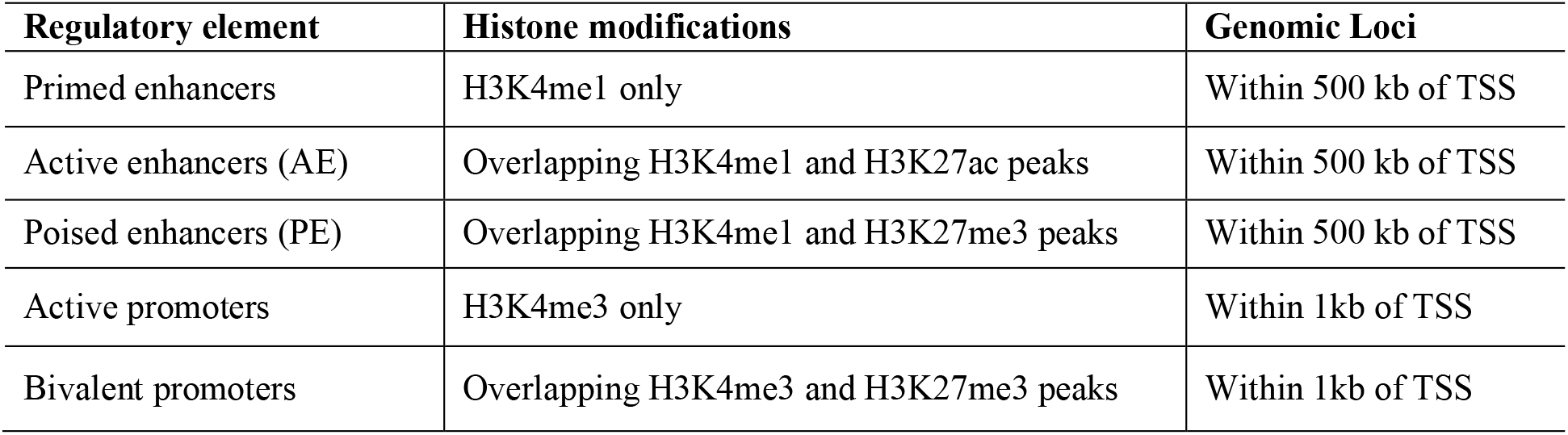
Histone modifications used to define gene regulatory elements

| Regulatory element | Histone modifications | Genomic Loci |
| --- | --- | --- |
| Primed enhancers | H3K4me1 only | Within 500 kb of TSS |
| Active enhancers (AE) | Overlapping H3K4me1 and H3K27ac peaks | Within 500 kb of TSS |
| Poised enhancers (PE) | Overlapping H3K4me1 and H3K27me3 peaks | Within 500 kb of TSS |
| Active promoters | H3K4me3 only | Within 1kb of TSS |
| Bivalent promoters | Overlapping H3K4me3 and H3K27me3 peaks | Within 1kb of TSS |

### Peptide preparation, isobaric labeling, and off-line fractionation for LC-MS

The frozen cell pellets (~10 million cells) were solubilized [35] in 0.5 mL of 8 M urea buffer (8 M urea, 75 mM NaCl, 50 mM Tris (pH 8.0), 1 mM EDTA, 2μg/mL aprotinin, 10 μg/mL leupeptin, 1 mM PMSF, 1:100 vol/vol Phosphatase Inhibitor Cocktail 2, 1:100 vol/vol Phosphatase Inhibitor Cocktail 3, 10 mM NaF) with ultrasonication using a Covaris S220X sonicator (Peak Incident Power: 150W, Duty Factor: 10%, cycles/burst: 500, time: 8min, temp: 4°C). The protein content was determined by the BCA method, as shown in Supp Table 1. For the reference pool, 60μg from each sample was combined, and 2 x 250 μg were processed with the samples. A protein aliquot (250 μg) was digested with trypsin after reduction and alkylation of disulfide bonds. Peptides were prepared and labeled with tandem mass tag reagents prior to off-line fractionation using high-pH reversed-phase chromatography [35]. Aliquots of the twenty-five fractions (~0.5 mg) were analyzed using LC-MS. The 25 fractions were further combined to 13 fractions for phosphopeptide enrichment as previously described [35] and analyzed by LC-MS.

### nano-LC-MS

The samples in 1% (vol/vol) aqueous FA were loaded (2.5 μL) onto a 75μm i.d. × 50cm AcclaimÒ PepMap 100 C18 RSLC column (Thermo-Fisher Scientific) on an EASY nano-LC (Thermo Fisher Scientific). The column was equilibrated using constant pressure (700 bar) with 11μL of solvent A (1% (vol/vol) aqueous FA). The peptides were eluted using the following gradient program with a flow rate of 300 L/min and using solvents A and B (1% (vol/vol) FA/MeCN): solvent A containing 5% B for 5 min, increased to 23% B over 105 min, to 35% B over 20 min, to 95% B over 1 min and constant 95% B for 19 min. The data were acquired in data-dependent acquisition (DDA) mode. The MS1 scans were acquired with the Orbitrap™ mass analyzer over m/z = 350 to 1500 and resolution set to 70,000. Twelve data-dependent high-energy collisional dissociation spectra (MS2) were acquired from each MS1 scan with a mass resolving power set to 35,000, a range of m/z = 100 – 2000, an isolation width of 1.2 m/z, and a normalized collision energy setting of 32%. The maximum injection time was 60 ms for parent-ion analysis and 120 ms for product-ion analysis. The ions that were selected for MS2 were dynamically excluded for 40 sec. The automatic gain control (AGC) was set at a target value of 3e6 ions for MS1 scans and 1e5 ions for MS2. Peptide ions with charge states of one or ≥ 7 were excluded for HCD acquisitions.

### Protein Identification

The unprocessed MS data from the mass spectrometer were converted to peak lists using Proteome Discoverer (version 2.1.0.81, Thermo-Fischer Scientific) with the integration of reporter-ion intensities of TMT 10-plex at a mass tolerance of ±3.15 mDa [36]. The MS2 spectra with charges +2, +3, and +4 were analyzed using Mascot software [37] (Matrix Science, London, UK; version 2.5.1). The mascot was set up to search against a SwissProt database of human (version June 2016, 20,237 entries) and common contaminant proteins (cRAP, version 1.0 Jan. 1st, 2012, 116 entries), assuming the digestion enzyme was trypsin/P with a maximum of 4 missed cleavages allowed. The searches were performed with a fragment ion mass tolerance of 0.02 Da and a parent ion tolerance of 20 ppm. Carbamidomethylation of cysteine was specified in Mascot as a fixed modification. Deamidation of asparagine, the formation of pyroglutamic acid from N-terminal glutamine, acetylation of protein N-terminus, oxidation of methionine, and pyro-carbamidomethylation of N-terminal cysteine were specified as variable modifications. Peptide spectrum matches (PSM) were filtered at a 1% FDR by searching against a reversed database, and the ascribed peptide identities were accepted. The uniqueness of peptide sequences among the database entries was determined using the principle of parsimony. Protein identities were inferred using a greedy set cover algorithm, and the identities containing ≥2 Occam’s razor peptides were accepted [38].

### Protein Relative Quantification

The processing, quality assurance, and analysis of TMT data were performed with proteoQ (version 1.0.0.0, https://github.com/qzhang503/proteoQ), a tool developed with the tidyverse approach (Wickham, 2017; Wickham, 2019) under the free software environment for statistical computing and graphics, R (R Core Team (2019). R: A language and environment for statistical computing. R Foundation for Statistical Computing, Vienna, Austria). URL [39] *f*s2and RStudio (RStudio Team (2016). RStudio: Integrated Development for R. RStudio, Inc., Boston, MA URL http://www.rstudio.com/). Briefly, reporter-ion intensities under 10-plex TMT channels were first obtained from Mascot, followed by the removals of PSM entries from shared peptides or with intensity values lower than 1E3. The intensity of PSMs was converted to logarithmic ratios at base two, relative to the average intensity of reference samples within a 10-plex TMT. Under each TMT channel, Dixon’s outlier removals were carried out recursively for peptides with greater than two identifying PSMs. The median of the ratios of PSM that can be assigned to the same peptide was first taken to represent the ratios of the incumbent peptide. The median of the ratios of peptides was then taken to represent the ratios of the incumbent protein.

To align protein ratios under different TMT channels, likelihood functions were first estimated for the log-ratios of proteins using finite mixture modelling, assuming two-component Gaussian mixtures (R package: mixtools: normalmixEM) [39]. The ratio distributions were then aligned in that the maximum likelihood of the log-ratios are centered at zero for each sample. Scaling normalization was performed to standardize the log-ratios of proteins across samples. To discount the influence of outliers from either log-ratios or reporter-ion intensities, the values between the 5th and 95th percentile of log-ratios and 5th and 95th percentile of intensity were used in the calculations of the standard deviations.

### Informatic and Statistical Analysis

Metric multi-dimensional scaling (MDS) and principal component analysis (PCA) of protein log2-ratios were performed using the base R function stats:cmdscale and stats:prcomp, respectively. Heat-map visualization of protein log2-ratios was performed with pheatmap. Linear modeling was performed using the contrast fit approach in Limma [33] to assess the statistical significance in protein abundance differences between indicated groups of contrasts. Adjustments of p-values for multiple comparisons were performed with Benjamini-Hochberg correction.

## Results

### Validation of *KDM6A* knockout in hiPSCs

To confirm *KDM6A* deletion in hiPSCs, we performed immunofluorescent staining for KDM6A. Compared with the wild type (WT) cells, we observed a complete loss of KDM6A protein in hiPSCs (Figure 1A,B). Consistent with this, the mRNA expression of KDM6A was almost undetectable in *KDM6A*KO hiPSCs in RNA-seq data (Supp Figure 1). The immunofluorescence results were further validated via nano-LC/MS data demonstrating that the KDM6A protein and phospho-protein levels were significantly reduced in *KDM6A*KO cells compared to isogenic WT type (Figure 1C,D).

**Figure 1:**
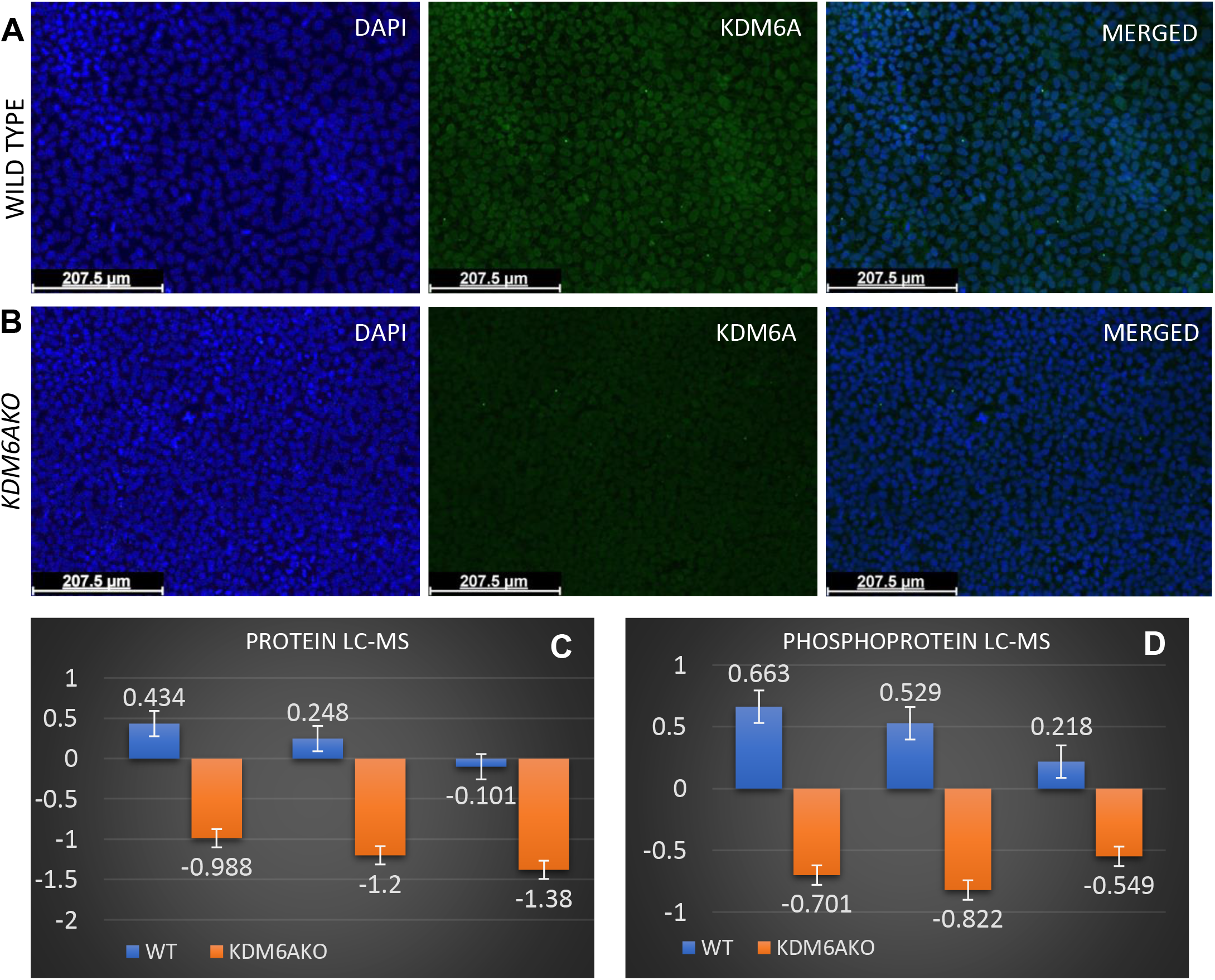
Validating lack of KDM6A protein expression in hiPSCs. (A) Representative immunostaining for DAPI, KDM6A and both merged in wild type hiPSCs, (B) the same staining in isogenic *KDM6A*KO cells demonstrates a lack of KDM6A protein staining. LC-MS protein (C) and phosphoprotein (D) quantification for KDM6A in triplicate wild type (WT) and *KDM6A*KO samples.

### Loss of KDM6A does not affect pluripotency and self-renewal of hiPSCs

Next, we examined if the loss of KDM6A altered pluripotency. We observed no morphological differences between WT and KDM6AKO cells (Supp Figure 2). Next, we successfully induced all three germ layers (ectoderm, mesoderm, and endoderm) in WT and *KDM6A*KO cells (Figure 2). In addition, alkaline phosphatase staining revealed that, like WT hiPSCs, *KDM6A*KO hiPSCs self-renew (Supp Figure 3). In summary, this observation supports the idea that *KDM6A* deletion does not fundamentally alter pluripotency, self-renewal capacity, and cell proliferation in hiPSCs.

**Figure 2:**
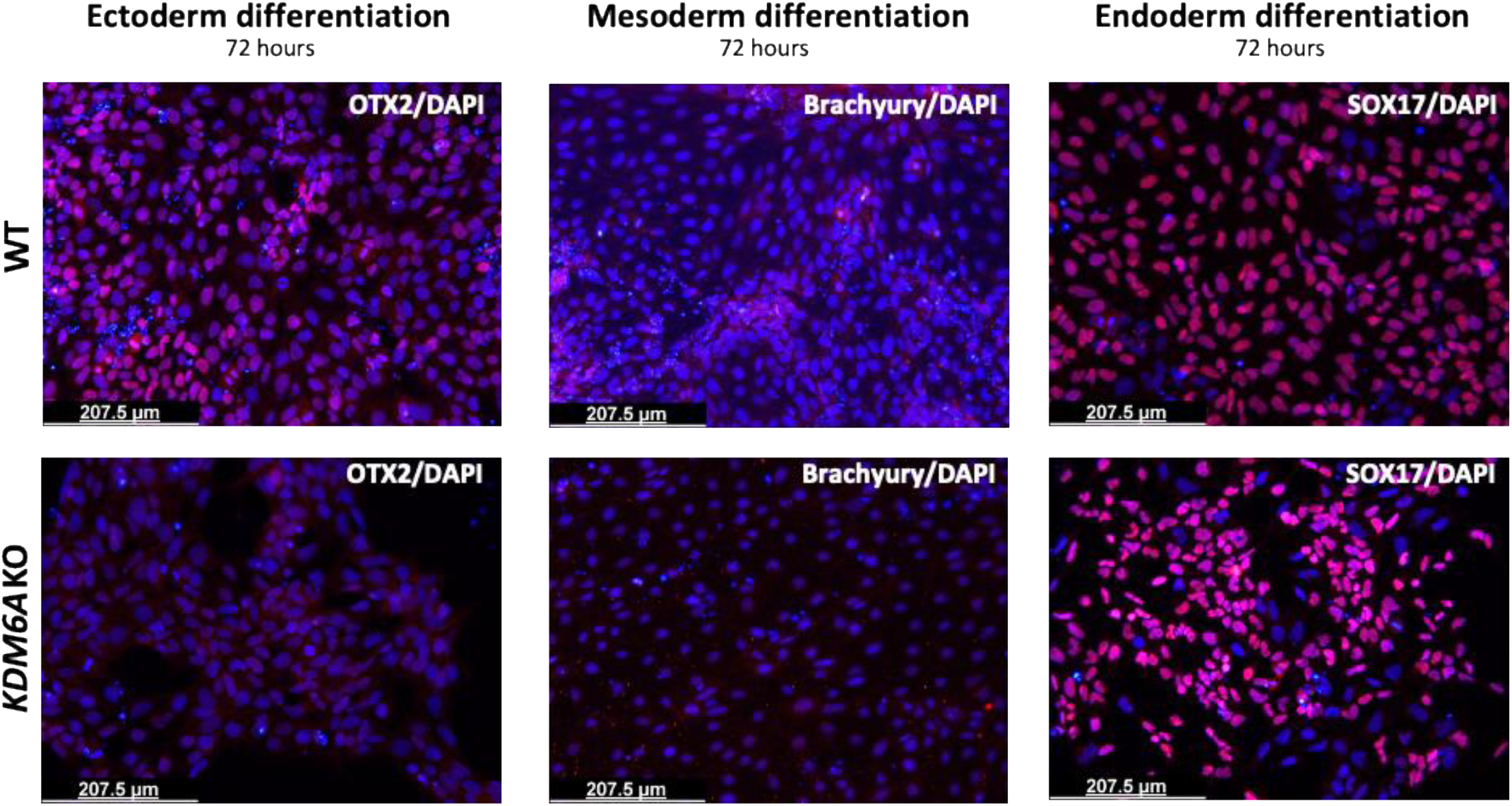
*KDM6A*KO does not alter the pluripotent phenotype of hiPSCs. WT and *KDM6A*KO hiPSCs cells were differentiated to ectoderm, mesoderm and endoderm using supplemental media. To further evaluate lineage commitment, cells were stained with goat anti-human OTX2 antibody, goat anti-human Brachyury antibody, and goat anti-human SOX17 antibody per manufacturer’s protocol.

### *KDM6A*KO alters histone modifications at promoters near transcription start sites (TSS)

KDM6A demethylates H3K27me3 histone marks during developmental transitions but knock-out murine models have shown some compensatory activation via KDM6C on the Y-chromosome in males (we are using an hiPSC line from an adult male) [11,18] To characterize the alterations of histone modifications due to KDM6A knockout in hiPSCs in an unbiased fashion, we performed ChIPmentation using specific antibodies against H3K4me1, H3K4me3, H3K27ac, and H3K27me3. Genome-wide differences in the modifications of these cis-regulatory elements in WT compared to *KDM6A*KO hiPSCs were noted (Supp Figure 4). The human genome was portioned into 10 distinct bins relative to RefSeq genes corresponding to the near promoter (≤1kb upstream of the TSS), mid-range promoter (1-2kb upstream of the TSS), distal promoter (2-3kb upstream of the TSS), 5’UTR, 3’UTR, 1^st^ exon, other intron, and distal intergenic regions. Genome-wide sites of histone modifications were assigned to these bins, and percentages were calculated (Table 2). As expected, the lack of KDM6A activity resulted in increases in H3K27me3 peaks at near promoters (2.18%) and 1^st^ exons (6.14%) in *KDM6A*KO hiPSCs compared to WT. Interestingly, near promoters also showed substantial increases in H3K4me3 (14.47%) marks, consistent with active promoters, as well as H3K27ac marks (17.04%). The increase in H3K4me3 and H3K27ac marks at near promoters appears to be compensatory to the 12.50% and 18.89% decreases, respectively, in marks at 1^st^ intron, other intron and distal intergenic regions, collectively. When overlapping the genomic locations of histone marks between WT and *KDM6A*KO cells, we observed considerable difference in genomic loci with all histone marks except H3K4me3 within 1kb of a transcription start site (Supp Figure 5). These data suggest that loss of KDM6A would be expected to have the broadest effect on enhancers, which are marked by H3K4me1, H3K27ac and H3K27me3.

**Table 2:** Genome-wide comparison of histone modification patterns between WT and *KDM6A*KO cells. The human genome was portioned into 10 bins relative to RefSeq genes. The table shows the percentage of the human genome with the respective histone modification mark in each bin. Red rectangles highlight data analyzed and discussed in the Results.

| Histone Marks | Cell line | Relative genomic location of peaks (%) |  |  |  |  |  |  |  |  |  |
| --- | --- | --- | --- | --- | --- | --- | --- | --- | --- | --- | --- |
|  |  | Near Promoter (<=1kb) | Mid-range Promoter (1-2kb) | Distal Promoter (2-3kb) | 5'UTR | 3'UTR | 1st Exon | Other Exon | 1st Intron | Other Intron | Distal intergenic |
| H3K4Me1 | WT | 15.53 | 3.09 | 1.37 | 1.61 | 3.12 | 0.58 | 6.82 | 11.43 | 19.6 | 36.36 |
|  | *KDM6A*KO | 12.8 | 3.85 | 1.43 | 1.56 | 2.67 | 0.5 | 6.54 | 10.78 | 20.45 | 38.97 |
| H3K4me3 | WT | 40.11 | 1.42 | 1.88 | 1.66 | 3.56 | 1.12 | 5.53 | 5.44 | 12.09 | 26.48 |
|  | *KDM6A*KO | 54.58 | 0.98 | 1.42 | 1.58 | 3.31 | 1.42 | 4.59 | 4.2 | 7.23 | 20.08 |
| H3K27me3 | WT | 8.49 | 2.49 | 1.4 | 0.89 | 1.68 | 0.59 | 3.84 | 6.66 | 15.73 | 57.59 |
|  | *KDM6A*KO | 10.67 | 1.71 | 1.25 | 0.95 | 1.68 | 6.73 | 3.33 | 6.73 | 15.82 | 56.51 |
| H3K27ac | WT | 8.39 | 1.47 | 1.67 | 3.21 | 4.01 | 0.49 | 12.87 | 14.49 | 28.03 | 24.87 |
|  | *KDM6A*KO | 25.43 | 2.3 | 2.62 | 3.2 | 5.87 | 0.55 | 10.95 | 9.72 | 16.32 | 22.46 |

### KDM6A loss primarily alters poised enhancers (PE) and active enhancers (AE) with binding motifs for critical developmentally-regulated TFs

We next examined the KDM6A-dependent differences in active and poised enhancers. “Primed” enhancers are marked by H3K4me1 only. “Active” enhancers (AE), which correlate with tissue-specific gene expression, are marked by H3K4me1 and H3K27ac [40]. “Poised” enhancers (PE), which correlate with potential gene expression at subsequent developmental stages, are simultaneously marked by H3K4me1 and H3K27me3 [40–42] within 500 kb of a TSS, driving gene expression later in the development [43]. The loss of H3K27me3, coupled with the acquisition of H3K27ac, endows these enhancers with gene regulatory functions and converts PE to AE when coupled with H3K4me1 marks.

As demethylation of H3K27 is a direct target of KDM6A, we expected that a loss of KDM6A would result in increased H3K27me3 marks overall and therefore increased numbers of PE. In WT compared to isogenic *KDM6A*KO hiPSCs, we identified 4,563 versus 2,892 unique overlapping H3K4me1 and H3K27me3 marks, respectively. Therefore, the absence of KDM6A resulted in decreased PE (Figure 3A) in hiPSCs with considerable differences in genomic loci (Supp Figure 4), indicating that the increases in H3K4me3 noted above near TSS and first exons were not associated with PE.

**Figure 3.**
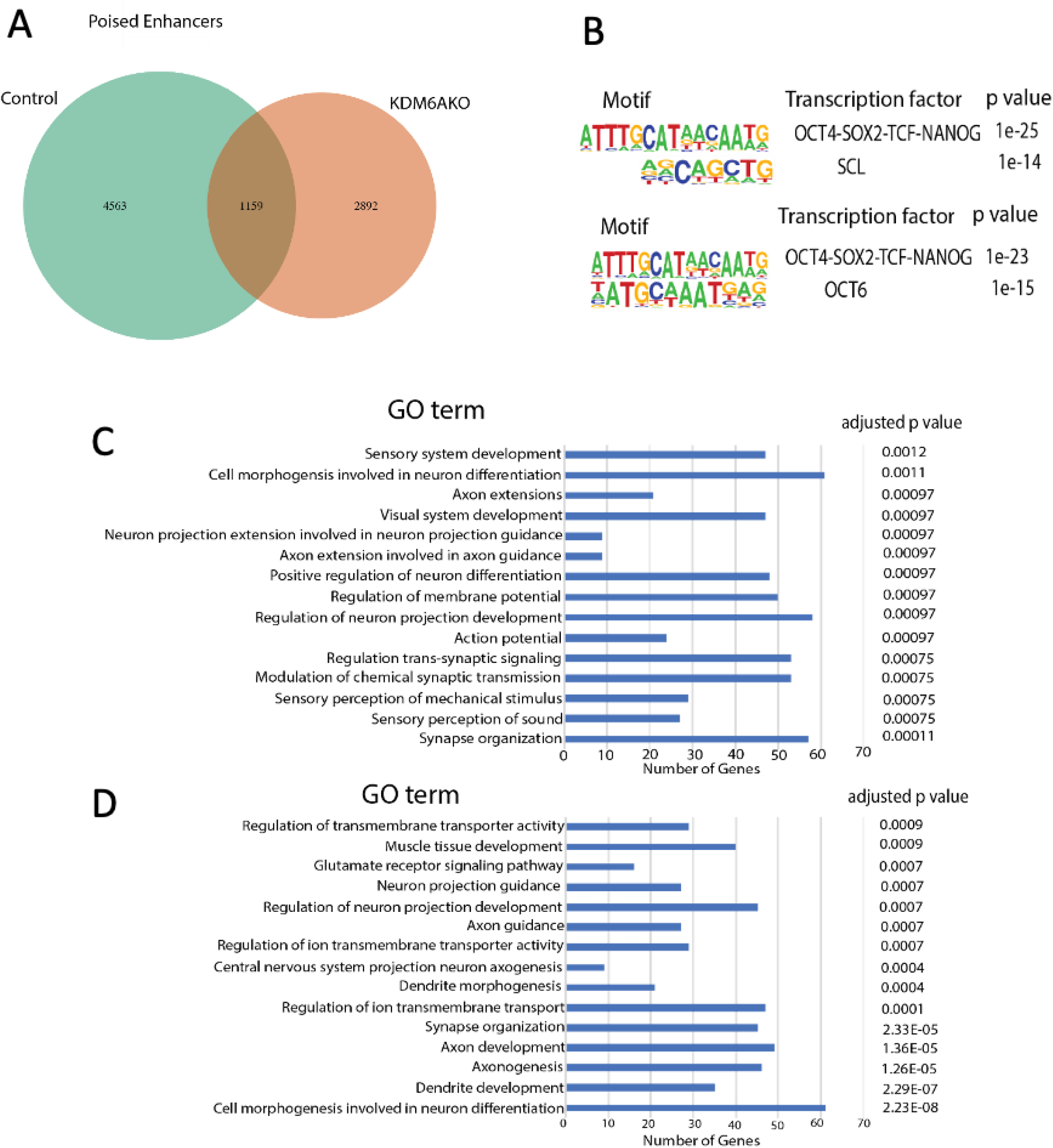
Annotation of the PE landscape. (A) Venn diagram of PE, (B) WT-specific TF binding site motifs associated with PE, (C) *KDM6A*KO-specific TF binding site motifs associated with PE. (D) WT-specific Gene Ontology (GO) terms associated with PE, (E) *KDM6A*KO-specific GO terms associated with PE.

Histone modification signatures act as binding sites for pioneer and lineage-determining transcription factors (TF) [44–47]. To identify TF binding sites within these differential regions, we applied motif analyses individually for WT- and *KDM6A*KO-specific PE signatures. In WT, the only significantly enriched PE motif was the developmentally-associated cooperative binding site for OCT4/POU5F1-SOX2-TCF-NANOG and SCL (Figure 3B). In comparison, the absence of KDM6A was also significantly different for the OCT4/POU5F1/-SOX2-TCF-NANOG binding site as well as the OCT6 binding motif (Figure 3B), suggesting that KDM6A mediates specific genomic locations for OCT4/POU5F1-SOX2-TCF-NANOG binding in pluripotency. With respect to these differences in genomic TF binding, the Gene Ontology (GO) terms associated with the PE marks differ entirely between WT and *KDM6A*KO hiPSCs (Figure 3C,D).

Next, we performed the same comparison for the AE landscape between WT and *KDM6A*KO. As shown in the Venn diagram of Figure 4A, there were a total of 14,653 active enhancer peaks called. Of these, only 1,866 (12.73%) were independent of *KDM6A*KO and shared between both lines. In contrast, 5,005 active enhancer peaks were specific to the *KDM6A*KO line, while 7,782 were specific to wild type. The AE landscape is mainly shaped by the cooperative binding of ubiquitous and cell-type-specific TFs [48]. The only significantly enriched AE motifs were again for the developmentally-regulated and cell-type specification cooperative binding sites for OCT4/POU5F1-SOX2-TCF-NANOG and OCT4/POU5F1 (Figure 4B). Similar to PE, the absence of KDM6A demonstrated a significant difference in the same OCT4/POU5F1-SOX2-TCF-NANOG (Figure 4B) motif, compared to WT. This also suggests that KDM6A is required for site-specific TF binding for AE, as well as PE. Figures 4C,D show the results of GO term analyses for the AE specific to WT versus *KDM6A*KO, respectively. In general, WT-specific AE-associated GO terms belong to positive regulation of neuron projection development, synapse organization, regulation of cell morphogenesis involved in differentiation and dendrite development. While *KDM6A*KO-specific AE GO terms falls under the categories of cell-cell signaling by WNT, positive regulation of GTPase activity, adherens junction organization, and Ras protein signal transduction. Overall, these data suggest that KDM6A mediates gene regulation for cell-type specific pathways.

**Figure 4.**
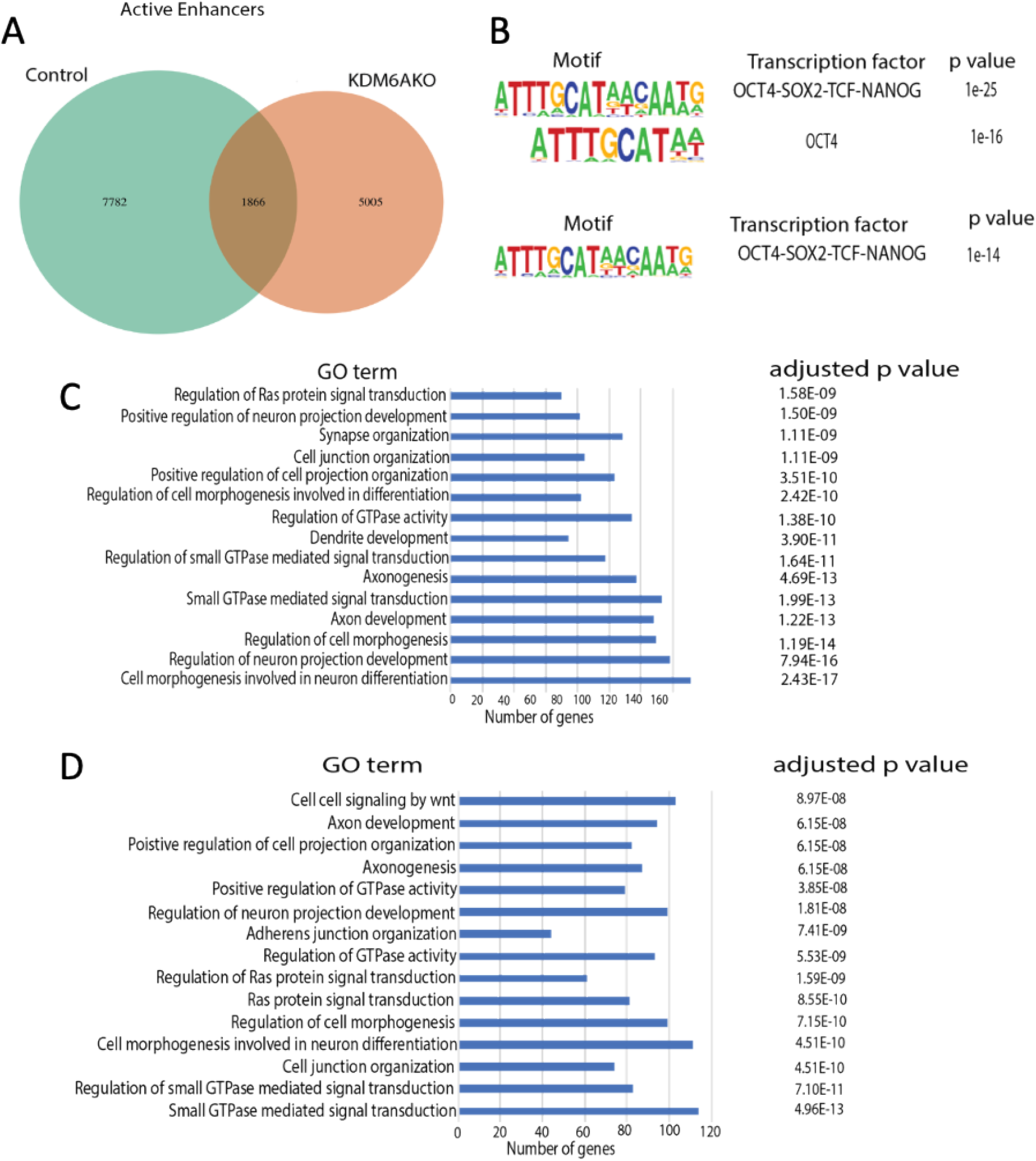
Annotation of the AE landscape. (A) Venn diagram of overlapping AE between WT and *KDM6A*KO hiPSCs. (B) WT-specific TF binding site motifs associated with AE, (C) *KDM6A*KO-specific TF binding site motifs associated with AE. (D) WT-specific Gene Ontology (GO) terms associated with AE, (E) *KDM6A*KO-specific GO terms associated with AE.

### Loss of KDM6A impact at bivalent promoters

Pluripotent cells are enriched for promoters harboring both the activating H3K4me3 as well as the repressive H3K27me3 marks at the same locus, a state called “bivalency”. While bivalent promoters are not unique to pluripotent cells, they are enriched in these cell types, mainly marking developmental and lineage-specific genes, which are generally static but can be rapidly activated or repressed and are more often associated with terminal differentiation [49].

KDM6A resolves bivalency at terminal differentiation steps by demethylating the repressive H3K27 mark converting the locus to an active promoter [50,51]. Thus, identifying the bivalent promoters still present after KDM6A loss in hiPSCs could help in predicting developmental mechanisms and defects. As shown in Supplementary Figure 6, a total of 4,536 bivalent promoters were observed in WT and *KDM6A*KO hiPSCs without significant differences in TF binding motifs, consistent with more of an impact at terminal differentiation steps. For the WT-specific bivalent promoters, there were no significantly enriched GO terms. Significantly enriched *KDM6A*KO-specific bivalent promoters and KDM6A-independent bivalent promoters shared in both cell lines are listed in Supplementary Figure 6B and C, respectively.

### Loss of KDM6A alters gene expression profiles in hiPSCs

We compared transcriptome sequencing identifying genes up- or downregulated >10-fold in *KDM6A*KO compared to WT (Table 3). In *KDM6A*KO, we observed the greatest upregulation in expression of *EIF2S3* (12.2-fold), and the greatest downregulation in *MAGEH1* (18.2-fold), both of which also reside on the X-chromosome with KDM6A. Like *KDM6A*, *MAGEH1* and *EIF2S3* escape X chromosome inactivation [52] indicative of gender-biased expression (2-fold in females compared to males). In addition to *EIF2S3*, two other X-chromosome genes were downregulated, *ACOT9* and *SAT1*. X-chromosome-related copy number variations involving these three genes have been associated with severe neurological defects[53,54] consistent with an important role for KDM6A-mediated regulation of ectoderm development as noted above in the loss of CNS-development associated GO terms specific to KDM6A loss for PE and AE (Figure 3D,4D).

**Table 3.** KDM6A-dependent changes in gene expression. Gene expression up or down-regulated by >10 fold in *KDM6A*KO compared to isogenic WT hiPSCs.

| Downregulated in *KDM6AKO* |  |  |  | Upregulated in *KDM6AKO* |  |  |  |
| --- | --- | --- | --- | --- | --- | --- | --- |
| Rank | Gene | Fold change (FC) | Log (FC) | Rank | Gene | Fold Change (FC) | Log(FC) |
| 1 | MAGEH1 | -18.1708 | 1.53E-15 | 1 | EIF2S3 | 12.15377 | 9.47E-12 |
| 2 | QPCT | -15.9647 | 2.72E-14 | 2 | ACOT9 | 10.69893 | 1.28E-10 |
| 3 | SPTSSB | -13.0566 | 2.11E-12 | 3 | MCM5 | 10.56954 | 1.63E-10 |
| 4 | ZNF208 | -11.4195 | 3.42E-11 | 4 | SAT1 | 10.40351 | 2.23E-10 |
| 5 | IRX2 | -10.8296 | 1.00E-10 |  |  |  |  |
| 6 | LITAF | -10.7509 | 1.16E-10 |  |  |  |  |
| 7 | ZNF257 | -10.6492 | 1.40E-10 |  |  |  |  |
| 8 | CXCL5 | -10.6056 | 1.52E-10 |  |  |  |  |
| 9 | DNAJC15 | -10.4078 | 2.22E-10 |  |  |  |  |
| 10 | CRYGD | -10.1294 | 3.78E-10 |  |  |  |  |

### Proteome and phospho-proteome mass spectroscopy reveal concomitant loss of EZH2, KMT2C and KMT2D in *KDM6A*KO hiPSCs

Since proteins are the ultimate functional effectors of activity in biological systems, we sought to correlate our epigenetic and expression results with an unbiased survey of global proteomic and phospho-proteomic expression. *KDM6A*KO displayed consistent changes in the basal proteomic and phosphorylation status of proteins (Figure 5A,B; Supp Figure 7). Among 1,601 differentially expressed proteins, 786 were upregulated, while 815 proteins were downregulated (Supp Table 2A,B). Only 1,368 phospho-proteins showed differential expression: 705 phosphoproteins were upregulated, and 663 proteins were downregulated (Supp Table 3A,B).

As KDM6A is an essential component of the KMT2C and KMT2D COMPASS complexes, we examined the expression of KMT2 proteins in our nanoLC/MS data. We observed a decrease in KMT2A/C/D protein expression, with the exception of KMT2B (Figure 5E). We further validated this finding with immunostaining for KMT2C (Figure 5F) and observed a similar decrease. As the literature indicates, the removal of KDM6A hampers the proper induction of ectoderm and mesoderm [11] and we also observed defective protein expression patterns of ectoderm and endoderm marker proteins, consistent with the loss of PE-specific enriched GO terms in the *KDM6A*KO hiPSCs (Figure 3D,E). Our RNA-seq data also shows the defective expression pattern for these marker genes (Supp Figure 8). However, mesodermal proteins were largely unaffected suggesting redundancy or regulation outside of the pluripotent state (Supp Table 4).

The only methyltransferase known for the H3K27me3 suppressive mark is the polycomb repressor complex protein, EZH2 [55–57]. As shown in Figure 6E, *KDM6A*KO hiPSCs demonstrated significantly decreased EZH2 expression (−1.14, 6.30e-03), consistent with the total decrease in H3K27me3 peaks after KDM6A loss (Supp Figure 5D).

**Figure 6.**
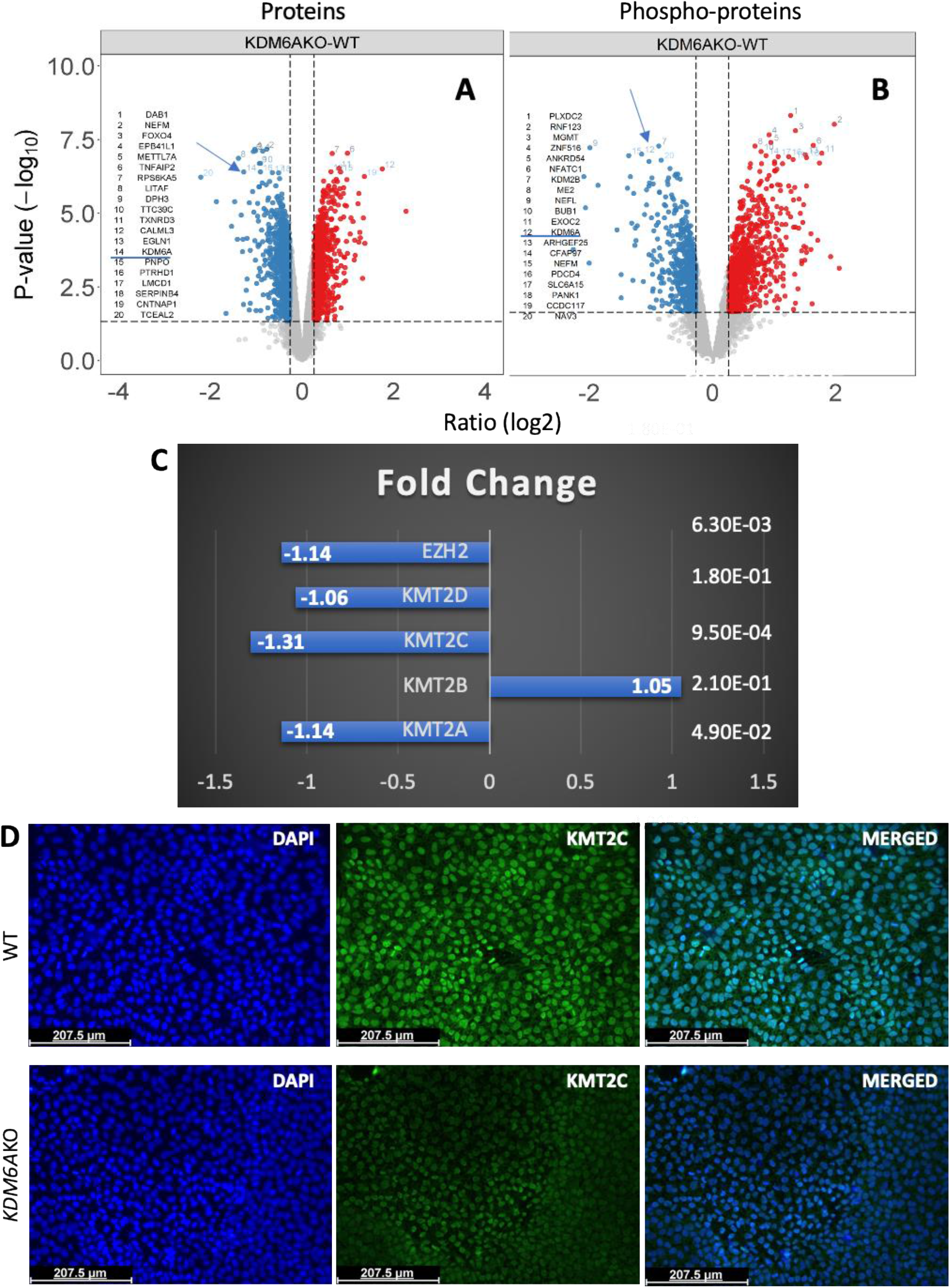
Proteomic and phospho-proteomic analyses. Mass spectrometry was performed to compare protein (A) and phospho-protein (B) expression between WT and *KDM6A*KO pluripotent cells. Both results show decreased protein expression of KDM6A (arrows). (C) Relative protein expression level of KMT2 and EZH2 proteins in nano LC/MS data. (D) Representative immuno-staining image of KMT2C in WT (top row) compared to *KDM6A*KO hiPSCs (bottom row).

## Discussion

KDM6A is known to regulate developmental gene expression via removal of trimethylation at H3K27, but its epigenomic targets in human pluripotent cells have not been characterized. Through a multi-omics approach, our data show that loss of KDM6A a) does not alter the pluripotent phenotype, consistent with prior work [58]; b) results in increases of H3K27me3, H3K4me3 and H3K27ac marks at near promoters with compensatory decreases in intronic loci; c) that these histone modifications primarily occur at PE and AE containing binding motifs for critical developmentally-regulated TFs; d) loss of KDM6A is accompanied by gene and protein expression decreases in the associated genes KMT2C, KMT2D and EZH2; e) loss of KDM6A demonstrates a consistent decrease in histone modification, gene and protein expression at loci associated with ectoderm development. While there is potential redundancy due to the presence of KDM6B and, in this male hiPSC line, KDM6C on the Y-chromosome, these results are consistent with prior work showing defects in ectoderm and mesoderm differentiation [11].

The KDM6A-dependent increase in H3K4me3, H3K27me3 and H3K27ac marks within one kb of TSS in concordance with a significant loss of a) GO terms associating these loci with ectoderm development, b) mRNA expression of genes (located on the X-chromosome like KDM6A) associated with nervous system and intellectual function, and c) protein expression of critical developmental regulators all suggest that KDM6A mediates epigenetic targets essential for ectoderm and, to a lesser degree, mesoderm specification. This is consistent with a report by Andricovich finding that loss of KDM6A deregulates COMPASS-like complexes and enhancer-chromatin profiles [59]. Dhar and colleagues defined KDM6A as a bivalency-resolving histone modifier necessary for stem cell differentiation [50], but more specifically, Tang reported that KDM6A regulates human neural differentiation and dendritic morphology by resolving bivalent promoters at terminal developmental steps [51].

Limitations to the current study include a single hiPSC line analysis without additional clones or a human ESC line for comparison. In addition, this line was kept in a pluripotent state in vitro without directed differentiation or xenotransplantation to characterize the impact of KDM6A-loss on terminal differentiation and cellular function. An expanded comparative study is needed to confirm these observations within dedicated lineages. Ongoing work within the lab will reveal more specific functional consequences of KDM6A loss in cell-type specific differentiation mechanisms. Overall, these results suggest that deletion of *KDM6A* significantly alters the cis-regulatory elements of the genome in a targeted and quantifiable manner that is expected to result in significant differences in tissue specification.

## Declarations

This work was funded by the Kellsie’s Hope Foundation to T.E.D.

The authors declare no conflicts of interest.

All data are available at GEO using accession number: GSE168252.

## Author Contributions

SSM and AB performed the experiments. WY performed the bioinformatic analysis on RNA-seq and ChIPmentation. QZ, PEG and RT performed the proteomics and subsequent analysis. TED oversaw the work and, with SSM, wrote and edited the manuscript and figures.

## Acknowledgements

We would like to acknowledge the Washington University Genome Engineering and iPSC Core for their assistance in establishing and validating the *KDM6A* knockout hiPSCs. We would also like to thank the Washington University Genome Technology Access Center.

