## Supplementary Figures for "KDM6A knockout in human iPSCs alters the genome-wide histone methylation profile at active and poised enhancers, activating expression of ectoderm gene expression pathways"

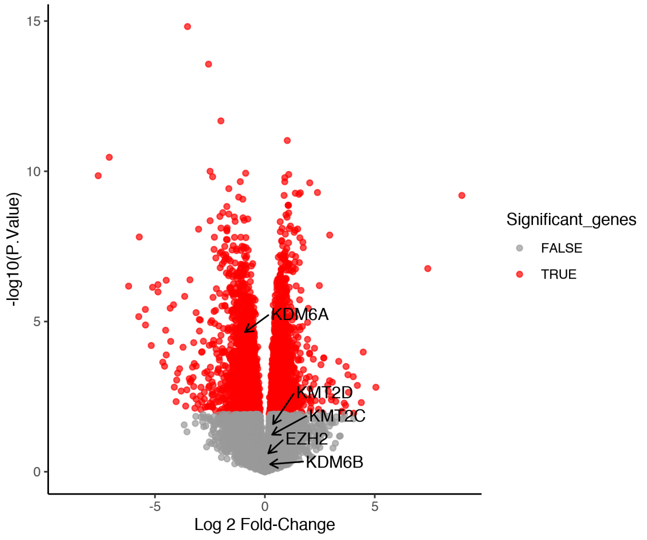


**Supplementary Figure 1:** RNA-seq shows reduced *KDM6A* gene expression

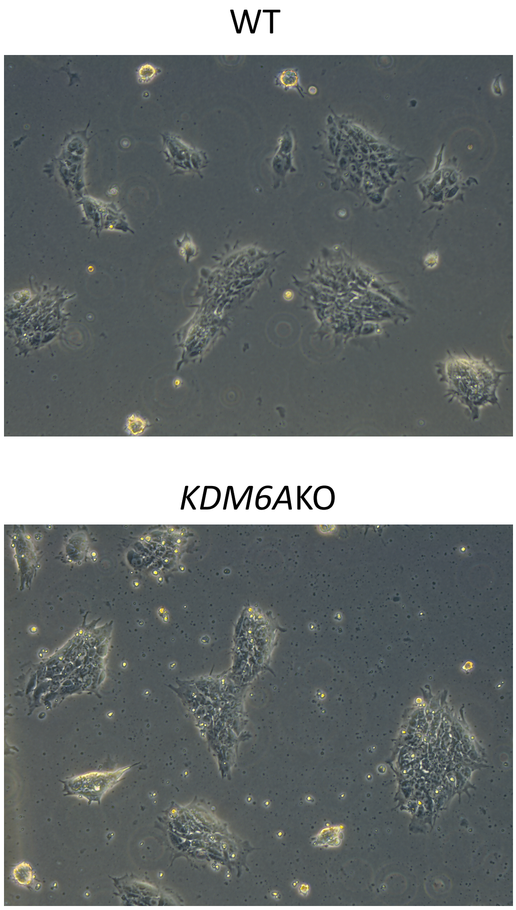


**Supplementary Figure 2:** *KDM6A*KO does not alter cellular morphology *in vitro* in hiPSCs. 20X magnification.

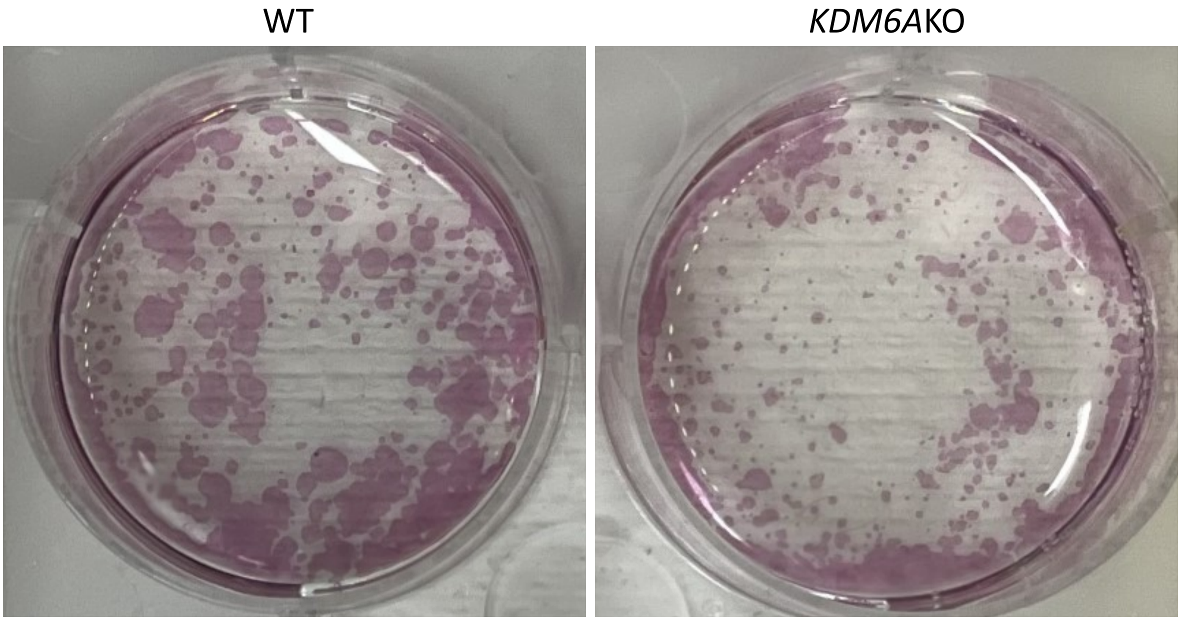


**Supplementary Figure 3:** Alkaline phosphatase staining shows comparable cell division

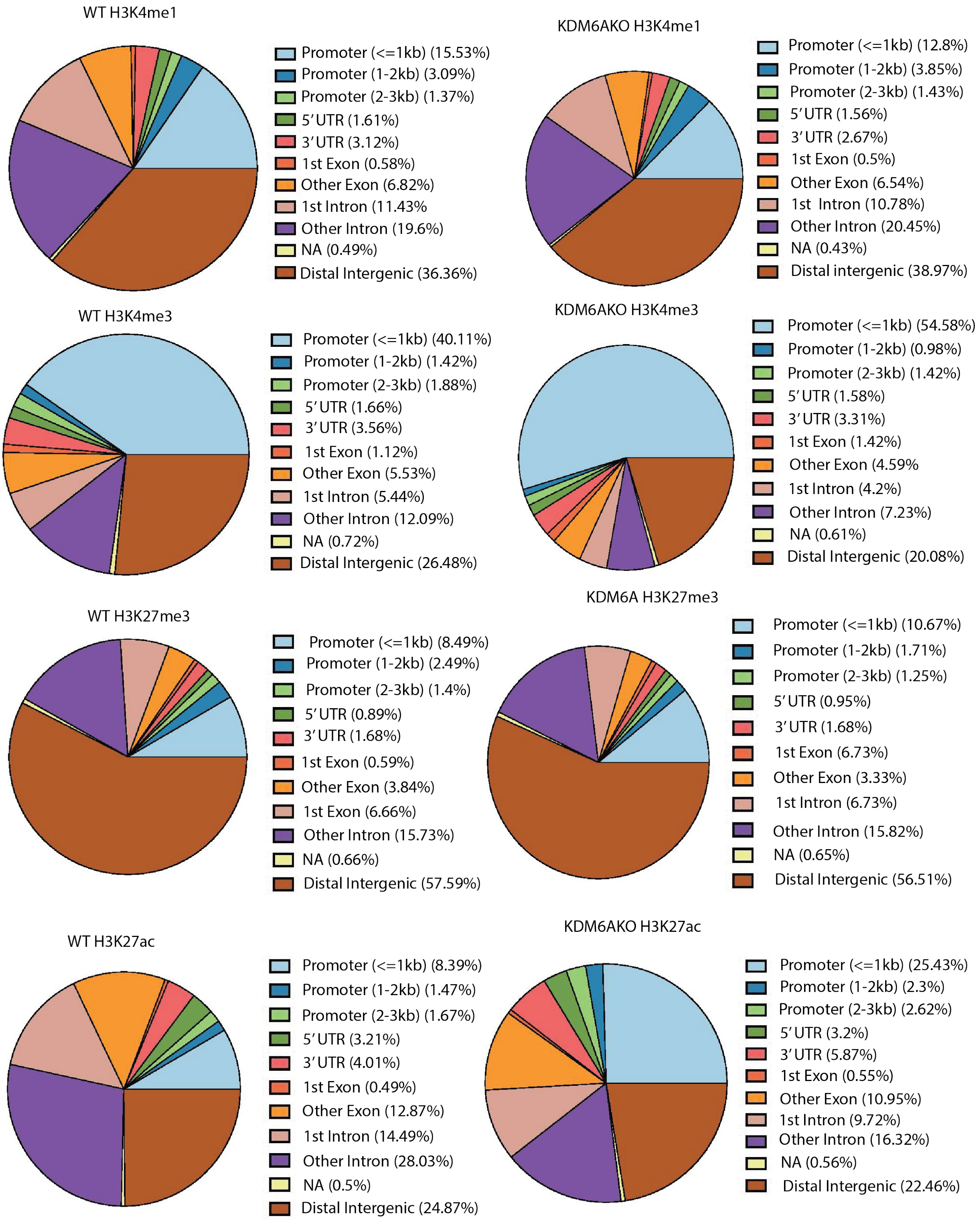


**Supplementary Figure 4:** Global distribution of histone marks in WT vs. *KDM6A*KO hiPSCs.

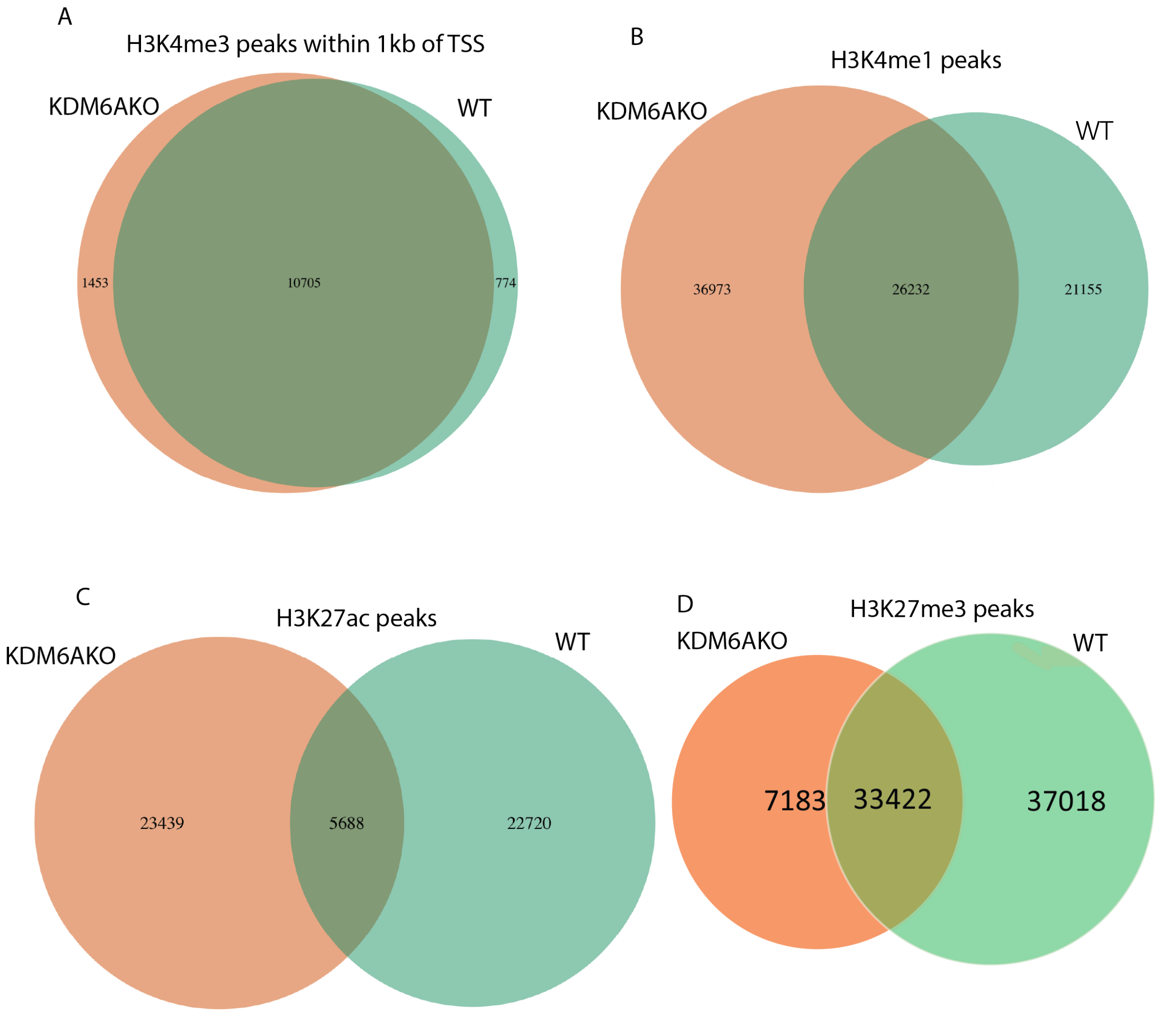


**Supplementary Figure 5: Overlapping peaks of histone marks in WT and *KDM6A*KO hiPSCs.** Venn diagrams showing the relative distribution of H3K4me3, H3K4me1, H3K27ac and H3K27me3 peaks between isogenic WT and *KDM6A*KO hiPSCs.

(A) Distribution of H3K4me3 peaks at near promoters within 1Kb of a transcription start site (TSS).
(B) Distribution of H3K4me1 peaks
(C) Distribution of H3K27ac peaks
(D) Distribution of H3K27me3 peaks

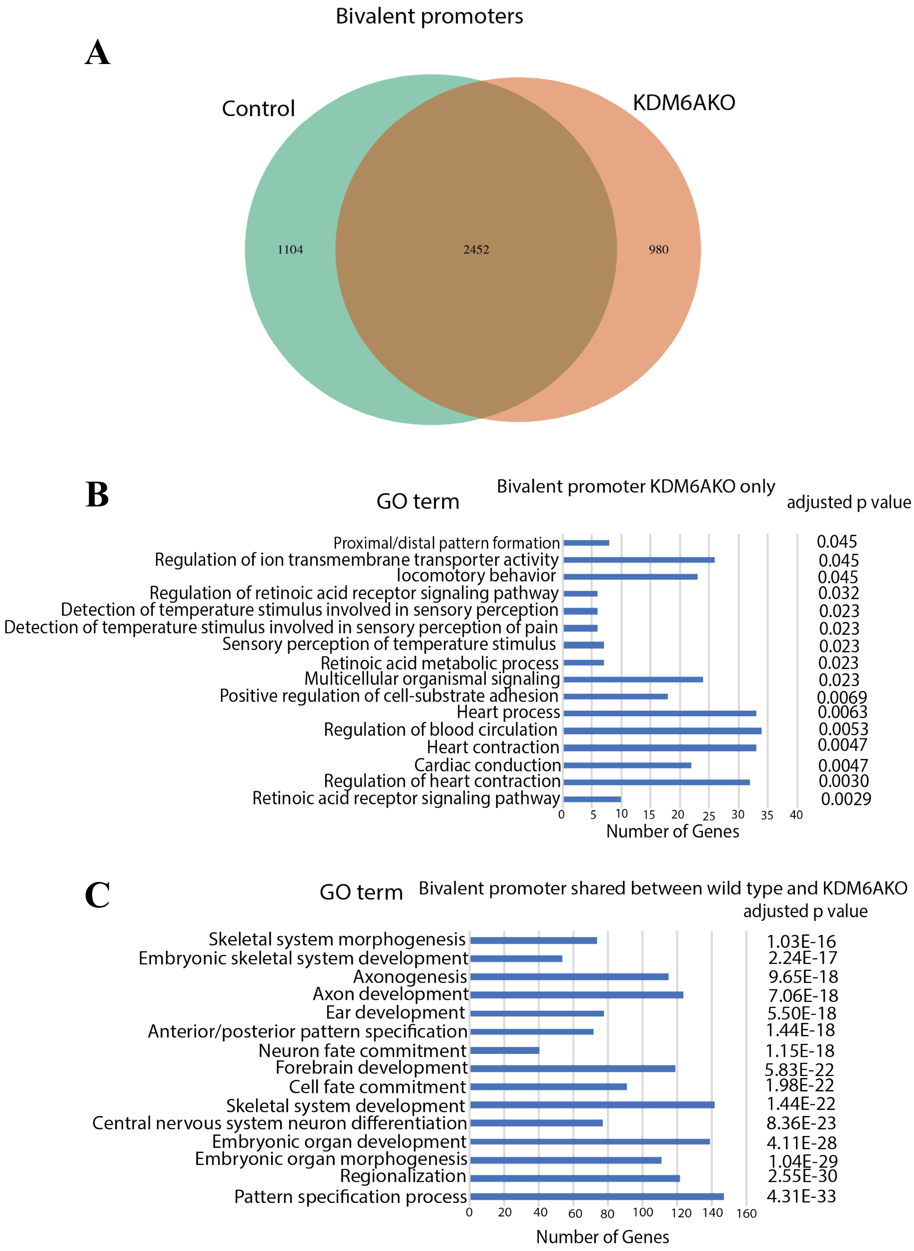


**Supplementary Figure 6. Annotation of the bivalent promoter landscape.** (A) Venn diagram of overlapping bivalent promoters between WT and *KDM6A*KO hiPSCs. (B) KDM6A-specific gene ontology (GO) terms associated with bivalent promoters, (C) GO terms associated with bivalent promoters shared between WT and *KDM6A*KO.

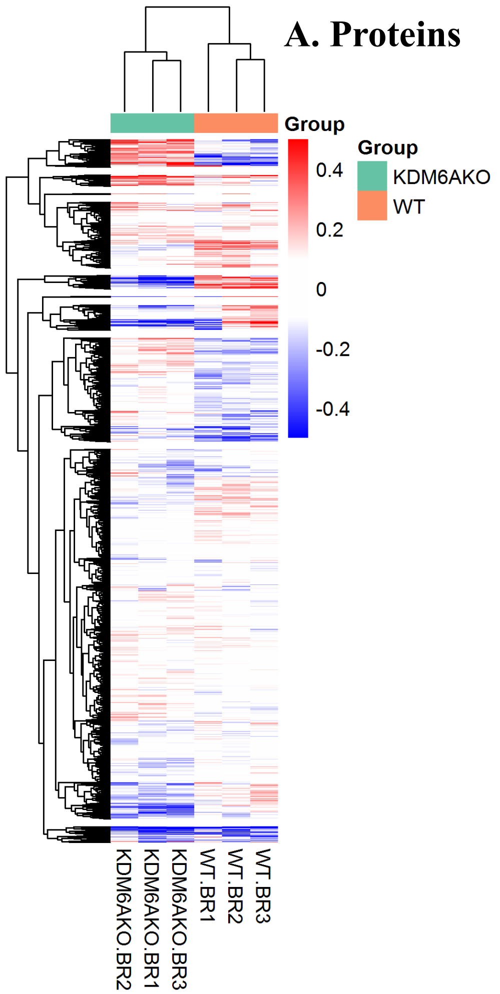


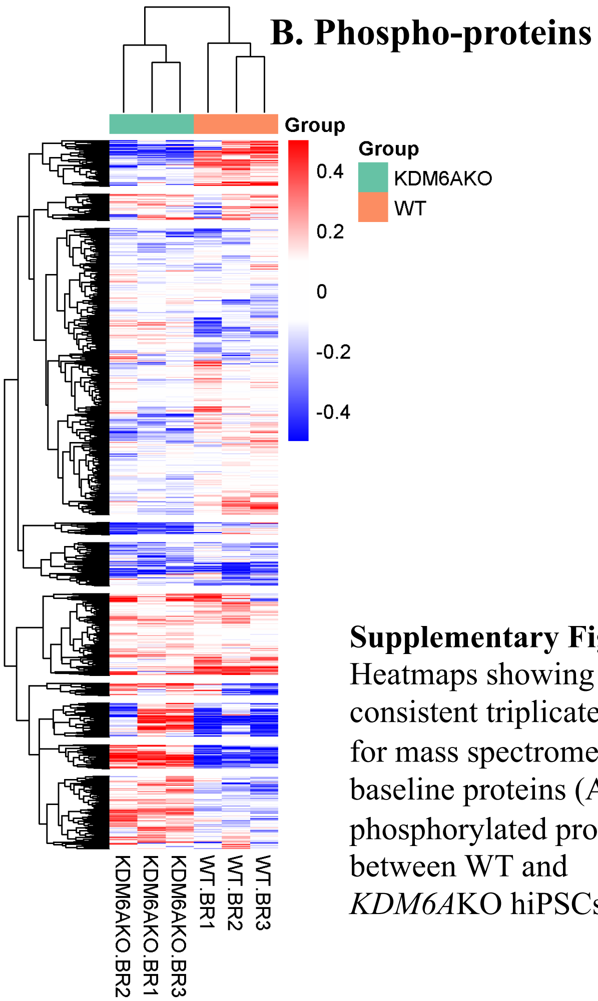


**Supplementary Figure 6**: Heatmaps showing consistent triplicate results for mass spectrometry for baseline proteins (A) and phosphorylated proteins (B) between WT and *KDM6A*KO hiPSCs.

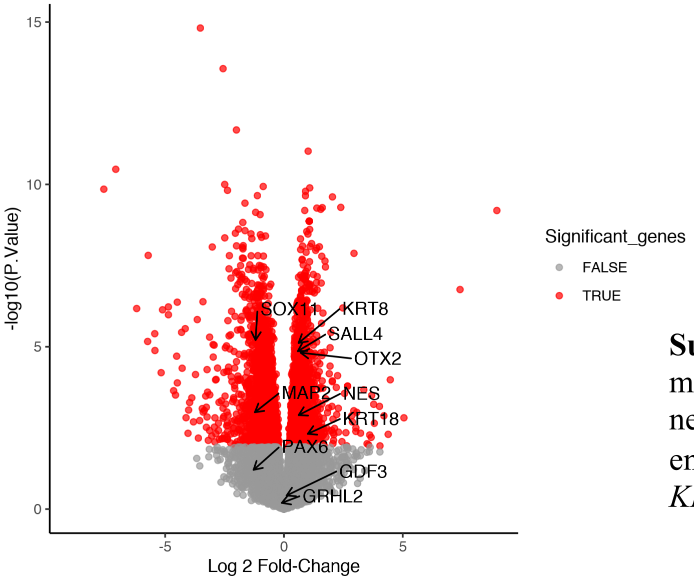


**Supplementary Figure 7**: Relative mRNA expression of select ectoderm, neuro-ectoderm and primitive endodermal genes between WT and *KDM6A*KO hiPSC.
