## Supplementary Tables for "KDM6A knockout in human iPSCs alters the genome-wide histone methylation profile at active and poised enhancers, activating expression of ectoderm gene expression pathways"

**Table of contents:**


| SUPPLEMENTARY TABLE | PAGE |
|---|---|
| 1) Protein concentration estimation by BCA method | 3 |
| 2A) *KDMA6A*KO upregulated protein expression by mass spectrometry | 4 |
| 2B) *KDM6A*KO downregulated protein expression by mass spectrometry | 24 |
| 3A) *KDM6A*KO upregulated phospho-protein expression by mass spec | 44 |
| 3B) *KDM6A*KO downregulated phospho-protein expression by mass spec | 62 |
| 4) Relative nano LC/MS protein expression of select germ layer markers | 79 |

**Supplementary Table 1: Protein concentration estimation by BCA method**

| Sample | Protein concentration µg/µl | Total Protein µg |
|---|---|---|
| WT, Replicate 1 | 5.17 | 1033 |
| WT, Replicate 2 | 4.37 | 1310 |
| WT, Replicate 3 | 2.38 | 475 |
| *KDM6A*KO, Repl 1 | 2.84 | 682 |
| *KDM6A*KO, Repl 2 | 4.84 | 1161 |
| *KDM6A*KO, Repl 3 | 5.93 | 1422 |

| **Supplementary Table 2A: *KDMA6A*KO upregulated protein expression by mass spec**<br>[pVal = p-value; logFC = log of the fold change; FC = fold change in expression] | | | | |
|---|---|---|---|---|
| **Protein** | **Accession** | **pVal** | **logFC** | **FC** |
| AARS | P49588 | 1.60E-04 | 0.4 | 1.32 |
| ABAT | P80404 | 1.70E-04 | 0.42 | 1.34 |
| ABCB6 | Q9NP58 | 3.20E-04 | 0.38 | 1.3 |
| ABCC2 | Q92887 | 0.012 | 0.29 | 1.22 |
| ACACB | O00763 | 4.40E-04 | 0.75 | 1.68 |
| ACAD10 | Q6JQN1 | 9.00E-05 | 0.37 | 1.3 |
| ACADS | P16219 | 4.10E-04 | 0.5 | 1.42 |
| ACADSB | P45954 | 9.20E-06 | 0.53 | 1.45 |
| ACAT1 | P24752 | 3.90E-04 | 0.35 | 1.27 |
| ACBD7 | Q8N6N7 | 4.50E-05 | 0.65 | 1.56 |
| ACCS | Q96QU6 | 0.0059 | 0.6 | 1.52 |
| ACO1 | P21399 | 0.0014 | 0.52 | 1.43 |
| ACOT9 | Q9Y305 | 8.00E-06 | 0.51 | 1.42 |
| ACSF2 | Q96CM8 | 1.30E-04 | 0.31 | 1.24 |
| ACSS3 | Q9H6R3 | 3.20E-05 | 0.44 | 1.36 |
| ACTRT1 | Q8TDG2 | 5.10E-05 | 1.05 | 2.07 |
| ACYP1 | P07311 | 0.004 | 0.42 | 1.34 |
| ACYP2 | P14621 | 0.0015 | 0.36 | 1.28 |
| ADA | P00813 | 3.20E-05 | 0.49 | 1.4 |
| ADAM23 | O75077 | 0.0018 | 0.39 | 1.31 |
| ADAM9 | Q13443 | 0.0064 | 0.3 | 1.24 |
| ADCY5 | O95622 | 0.0022 | 0.33 | 1.26 |
| ADGRA2 | Q96PE1 | 1.30E-04 | 0.61 | 1.53 |
| ADSSL1 | Q8N142 | 3.10E-04 | 0.45 | 1.36 |
| AGA | P20933 | 1.90E-04 | 0.52 | 1.43 |
| AGAP3 | Q96P47 | 6.40E-05 | 0.44 | 1.36 |
| AGO2 | Q9UKV8 | 9.90E-05 | 0.3 | 1.24 |
| AGO3 | Q9H9G7 | 4.90E-05 | 0.39 | 1.31 |
| AHNAK | Q09666 | 0.0096 | 0.37 | 1.29 |
| AHR | P35869 | 0.028 | 0.34 | 1.27 |
| ALAD | P13716 | 0.0051 | 0.38 | 1.31 |
| ALDH1L2 | Q3SY69 | 6.00E-05 | 0.66 | 1.58 |
| ALDH2 | P05091 | 1.60E-06 | 0.56 | 1.47 |
| ALDH5A1 | P51649 | 3.10E-06 | 0.46 | 1.37 |
| ALDH6A1 | Q02252 | 2.00E-04 | 0.37 | 1.29 |
| AMMECR1L | Q6DCA0 | 6.20E-04 | 0.37 | 1.29 |
| AMZ2 | Q86W34 | 0.008 | 0.3 | 1.23 |
| ANAPC15 | P60006 | 0.0076 | 0.37 | 1.29 |

| | | | | |
|---|---|---|---|---|
| ANGEL1 | Q9UNK9 | 0.026 | 0.27 | 1.21 |
| ANKDD1A | Q495B1 | 0.0048 | 0.32 | 1.25 |
| ANO1 | Q5XXA6 | 0.016 | 0.31 | 1.24 |
| AP1S1 | P61966 | 5.10E-04 | 0.32 | 1.25 |
| AP3B2 | Q13367 | 0.0036 | 0.39 | 1.31 |
| AP5B1 | Q2VPB7 | 0.0033 | 0.42 | 1.34 |
| APBB1 | O00213 | 3.30E-04 | 0.41 | 1.33 |
| APEH | P13798 | 1.70E-04 | 0.32 | 1.24 |
| APLF | Q8IW19 | 0.0034 | 0.34 | 1.27 |
| APOA1 | P02647 | 0.022 | 0.65 | 1.57 |
| APOBEC3B | Q9UH17 | 9.20E-07 | 0.77 | 1.7 |
| APOC1 | P02654 | 4.70E-04 | 0.47 | 1.39 |
| APOL2 | Q9BQE5 | 3.10E-06 | 0.83 | 1.78 |
| APOO | Q9BUR5 | 2.10E-06 | 0.51 | 1.42 |
| APPL2 | Q8NEU8 | 2.20E-05 | 0.45 | 1.37 |
| ARAF | P10398 | 7.90E-05 | 0.66 | 1.57 |
| ARHGEF10L | Q9HCE6 | 0.0026 | 0.3 | 1.23 |
| ARHGEF25 | Q86VW2 | 7.40E-04 | 0.78 | 1.71 |
| ARL1 | P40616 | 1.30E-04 | 0.27 | 1.2 |
| ARL2 | P36404 | 5.80E-05 | 0.4 | 1.32 |
| ARL2BP | Q9Y2Y0 | 4.00E-06 | 0.7 | 1.63 |
| ARPIN | Q7Z6K5 | 4.20E-05 | 0.67 | 1.59 |
| AS3MT | Q9HBK9 | 1.30E-04 | 0.43 | 1.35 |
| ASAH1 | Q13510 | 0.0044 | 0.33 | 1.26 |
| ASF1B | Q9NVP2 | 8.10E-06 | 0.39 | 1.31 |
| ASNS | P08243 | 6.30E-05 | 0.9 | 1.87 |
| ASS1 | P00966 | 2.70E-06 | 1.19 | 2.28 |
| ATHL1_HUMAN | NA | 0.001 | 0.3 | 1.23 |
| ATP1A2 | P50993 | 8.90E-06 | 0.7 | 1.62 |
| ATP1B4 | Q9UN42 | 0.002 | 0.95 | 1.93 |
| ATP6V0C | P27449 | 0.016 | 0.52 | 1.43 |
| B3GAT3 | O94766 | 0.0013 | 0.31 | 1.24 |
| BAG2 | O95816 | 6.20E-05 | 0.42 | 1.33 |
| BAP18 | Q8IXM2 | 4.30E-05 | 0.36 | 1.28 |
| BASP1 | P80723 | 0.0016 | 0.34 | 1.27 |
| BCAP29 | Q9UHQ4 | 2.60E-04 | 0.28 | 1.22 |
| BCAT1 | P54687 | 9.20E-04 | 0.46 | 1.38 |
| BCAT2 | O15382 | 3.90E-05 | 0.38 | 1.3 |
| BCDIN3D | Q7Z5W3 | 0.0028 | 0.53 | 1.44 |
| BCL9 | O00512 | 0.0038 | 0.32 | 1.25 |
| BDH2 | Q9BUT1 | 0.0033 | 0.26 | 1.2 |

| | | | | |
|---|---|---|---|---|
| BET1L | Q9NYM9 | 0.036 | 0.34 | 1.26 |
| BLVRA | P53004 | 3.00E-04 | 0.27 | 1.21 |
| BLVRB | P30043 | 8.10E-05 | 0.4 | 1.32 |
| BMT2 | Q1RMZ1 | 0.0065 | 0.66 | 1.58 |
| BRIP1 | Q9BX63 | 7.30E-04 | 0.26 | 1.2 |
| BST2 | Q10589 | 6.30E-04 | 0.62 | 1.54 |
| BTG3 | Q14201 | 0.0098 | 0.36 | 1.29 |
| C10orf76 | Q5T2E6 | 0.0019 | 0.34 | 1.27 |
| C10orf88 | Q9H8K7 | 1.20E-04 | 0.45 | 1.36 |
| C14orf1 | Q9UKR5 | 0.0029 | 0.43 | 1.35 |
| C16orf70 | Q9BSU1 | 0.0017 | 0.26 | 1.2 |
| C19orf12 | Q9NSK7 | 0.0086 | 0.4 | 1.32 |
| C1orf167 | Q5SNV9 | 1.40E-04 | 0.71 | 1.63 |
| C1orf174 | Q8IYL3 | 0.01 | 0.39 | 1.31 |
| C1orf226 | A1L170 | 7.70E-05 | 0.41 | 1.32 |
| C20orf194 | Q5TEA3 | 0.0015 | 0.26 | 1.2 |
| C3orf33 | Q6P1S2 | 0.011 | 0.37 | 1.29 |
| C6orf203 | Q9P0P8 | 0.0018 | 0.53 | 1.44 |
| CA8 | P35219 | 3.50E-05 | 0.33 | 1.26 |
| CACNA2D2 | Q9NY47 | 2.60E-04 | 0.35 | 1.27 |
| CACNB1 | Q02641 | 3.80E-04 | 0.45 | 1.37 |
| CALM_HUMAN | NA | 0.0017 | 0.85 | 1.81 |
| CALML3 | P27482 | 2.40E-07 | 1.74 | 3.34 |
| CAPN2 | P17655 | 3.50E-04 | 0.49 | 1.4 |
| CARS | P49589 | 7.10E-04 | 0.33 | 1.26 |
| CARS2 | Q9HA77 | 4.10E-04 | 0.31 | 1.24 |
| CASP7 | P55210 | 1.50E-05 | 0.4 | 1.32 |
| CASP8 | Q14790 | 4.00E-05 | 0.51 | 1.43 |
| CASZ1 | Q86V15 | 2.20E-04 | 0.29 | 1.22 |
| CBFB | Q13951 | 2.00E-04 | 0.8 | 1.75 |
| CBX5 | P45973 | 0.0039 | 0.28 | 1.22 |
| CCD53_HUMAN | NA | 0.004 | 0.36 | 1.29 |
| CCDC117 | Q8IWD4 | 1.80E-04 | 0.36 | 1.28 |
| CCDC127 | Q96BQ5 | 1.50E-05 | 0.39 | 1.31 |
| CCDC173 | Q0VFZ6 | 0.0015 | 0.71 | 1.64 |
| CCDC22 | O60826 | 1.60E-05 | 0.34 | 1.27 |
| CCDC87 | Q9NVE4 | 0.0046 | 0.27 | 1.21 |
| CCDC93 | Q567U6 | 3.50E-05 | 0.4 | 1.32 |
| CD63 | P08962 | 0.0025 | 0.41 | 1.33 |
| CD82 | P27701 | 0.042 | 0.32 | 1.25 |
| CD99 | P14209 | 6.10E-05 | 0.51 | 1.42 |

| | | | | |
|---|---|---|---|---|
| CDAN1 | Q8IWY9 | 1.30E-04 | 0.42 | 1.34 |
| CDKN2AIP | Q9NXV6 | 4.10E-05 | 0.33 | 1.25 |
| CDT1 | Q9H211 | 0.04 | 0.26 | 1.2 |
| CECR5_HUMAN | NA | 1.20E-04 | 0.29 | 1.23 |
| CEP112 | Q8N8E3 | 0.0011 | 0.39 | 1.31 |
| CEP44 | Q9C0F1 | 2.20E-04 | 0.28 | 1.22 |
| CEP63 | Q96MT8 | 0.035 | 0.27 | 1.2 |
| CEP70 | Q8NHQ1 | 0.035 | 0.35 | 1.27 |
| CERS1 | P27544 | 1.90E-05 | 0.34 | 1.27 |
| CFAP20 | Q9Y6A4 | 0.0023 | 0.28 | 1.21 |
| CFDP1 | Q9UEE9 | 0.00093 | 0.37 | 1.3 |
| CHD1L | Q86WJ1 | 6.60E-05 | 0.29 | 1.22 |
| CHD3 | Q12873 | 0.011 | 0.37 | 1.29 |
| CHD7 | Q9P2D1 | 2.50E-06 | 0.71 | 1.64 |
| CHDH | Q8NE62 | 7.90E-06 | 0.45 | 1.36 |
| CHGA | P10645 | 7.40E-04 | 0.93 | 1.91 |
| CHIC2 | Q9UKJ5 | 0.011 | 0.87 | 1.82 |
| CHID1 | Q9BWS9 | 1.30E-04 | 0.32 | 1.24 |
| CHURC1 | Q8WUH1 | 0.02 | 0.34 | 1.26 |
| CKS2 | P33552 | 0.0047 | 0.28 | 1.21 |
| CLDN3 | O15551 | 1.10E-04 | 1.24 | 2.36 |
| CLIC4 | Q9Y696 | 6.00E-04 | 0.3 | 1.23 |
| CLMN | Q96JQ2 | 8.60E-04 | 0.28 | 1.22 |
| CLU | P10909 | 3.20E-04 | 0.37 | 1.29 |
| CMBL | Q96DG6 | 0.0013 | 0.37 | 1.29 |
| CMC4 | P56277 | 7.70E-04 | 0.54 | 1.46 |
| CMPK1 | P30085 | 0.0013 | 0.36 | 1.29 |
| CMTM4 | Q8IZR5 | 0.0013 | 0.28 | 1.22 |
| CNTNAP1 | P78357 | 6.30E-07 | 1.35 | 2.56 |
| COL11A2 | P13942 | 0.0049 | 0.36 | 1.29 |
| COLEC10 | Q9Y6Z7 | 1.30E-04 | 0.31 | 1.24 |
| COMMD8 | Q9NX08 | 2.60E-05 | 0.55 | 1.46 |
| COMT | P21964 | 1.00E-06 | 0.72 | 1.64 |
| COQ2 | Q96H96 | 5.50E-04 | 0.49 | 1.41 |
| CPNE2 | Q96FN4 | 0.0043 | 0.32 | 1.25 |
| CRABP2 | P29373 | 1.80E-04 | 0.51 | 1.43 |
| CRB3 | Q9BUF7 | 0.0011 | 0.91 | 1.88 |
| CRIPT | Q9P021 | 0.0035 | 0.31 | 1.24 |
| CRTC3 | Q6UUV7 | 4.20E-04 | 0.6 | 1.52 |
| CSK | P41240 | 0.0033 | 0.27 | 1.21 |
| CTBS | Q01459 | 0.0062 | 0.34 | 1.27 |

| | | | | |
|---|---|---|---|---|
| CTH | P32929 | 2.90E-06 | 0.67 | 1.59 |
| CTSB | P07858 | 0.0013 | 0.38 | 1.3 |
| CTSH | P09668 | 6.70E-06 | 0.63 | 1.55 |
| CTSL | P07711 | 6.00E-04 | 0.58 | 1.49 |
| CTSZ | Q9UBR2 | 3.30E-06 | 0.48 | 1.4 |
| CUL5 | Q93034 | 1.40E-04 | 0.27 | 1.21 |
| CUZD1 | Q86UP6 | 1.50E-05 | 1.33 | 2.52 |
| CXorf23 | A2AJT9 | 0.0098 | 0.26 | 1.2 |
| CYP27A1 | Q02318 | 2.30E-06 | 0.89 | 1.86 |
| CYSTM1 | Q9H1C7 | 0.0015 | 0.57 | 1.48 |
| DCK | P27707 | 1.30E-04 | 0.52 | 1.43 |
| DCTD | P32321 | 3.80E-04 | 0.33 | 1.26 |
| DCUN1D1 | Q96GG9 | 0.0025 | 0.26 | 1.2 |
| DDHD2 | O94830 | 4.90E-06 | 0.93 | 1.91 |
| DDX59 | Q5T1V6 | 4.60E-04 | 0.31 | 1.24 |
| DEF6 | Q9H4E7 | 0.0033 | 0.31 | 1.24 |
| DEK | P35659 | 0.005 | 0.26 | 1.2 |
| DENND1A | Q8TEH3 | 1.20E-05 | 0.5 | 1.42 |
| DENND5B | Q6ZUT9 | 3.30E-04 | 0.92 | 1.9 |
| DENND6A | Q8IWF6 | 0.0013 | 0.31 | 1.24 |
| DEPDC5 | O75140 | 0.0018 | 0.31 | 1.24 |
| DERA | Q9Y315 | 4.80E-04 | 0.38 | 1.3 |
| DICER1 | Q9UPY3 | 5.00E-04 | 0.27 | 1.2 |
| DMXL1 | Q9Y485 | 5.30E-04 | 0.27 | 1.21 |
| DMXL2 | Q8TDJ6 | 0.04 | 0.36 | 1.29 |
| DNASE2 | O00115 | 1.60E-06 | 0.71 | 1.63 |
| DNPH1 | O43598 | 6.70E-04 | 0.42 | 1.33 |
| DOCK1 | Q14185 | 3.60E-04 | 0.28 | 1.22 |
| DOLPP1 | Q86YN1 | 0.002 | 0.3 | 1.23 |
| DPP7 | Q9UHL4 | 0.012 | 0.46 | 1.38 |
| DPY19L1 | Q2PZI1 | 8.20E-04 | 0.3 | 1.23 |
| DSP | P15924 | 2.10E-04 | 0.33 | 1.26 |
| DTD1 | Q8TEA8 | 0.012 | 0.38 | 1.3 |
| DUSP14 | O95147 | 0.0014 | 0.35 | 1.28 |
| ECHDC3 | Q96DC8 | 0.023 | 0.3 | 1.23 |
| EEF2KMT | Q96G04 | 0.0086 | 0.3 | 1.23 |
| EFNB1 | P98172 | 0.003 | 0.27 | 1.21 |
| EFS | O43281 | 0.001 | 0.37 | 1.29 |
| EGLN1 | Q9GZT9 | 4.50E-07 | 0.62 | 1.54 |
| EHBP1L1 | Q8N3D4 | 3.30E-05 | 0.74 | 1.67 |
| EIF2AK2 | P19525 | 1.20E-05 | 0.45 | 1.37 |

| | | | | |
|---|---|---|---|---|
| EIF4EBP1 | Q13541 | 0.0019 | 0.32 | 1.25 |
| ENGASE | Q8NFI3 | 0.0047 | 0.26 | 1.2 |
| ENO2 | P09104 | 0.017 | 0.42 | 1.34 |
| EPB42 | P16452 | 4.20E-06 | 0.58 | 1.49 |
| EPDR1 | Q9UM22 | 6.10E-04 | 0.84 | 1.79 |
| EPHA4 | P54764 | 0.047 | 0.33 | 1.26 |
| ERAP1 | Q9NZ08 | 3.90E-05 | 0.32 | 1.25 |
| ERCC6L2 | Q5T890 | 0.0032 | 0.33 | 1.25 |
| ESPN | B1AK53 | 6.50E-05 | 0.66 | 1.59 |
| EXOG | Q9Y2C4 | 0.01 | 0.26 | 1.2 |
| FA65B_HUMAN | NA | 0.025 | 0.3 | 1.23 |
| FABP6 | P51161 | 1.10E-04 | 0.73 | 1.66 |
| FAM114A2 | Q9NRY5 | 7.50E-04 | 0.26 | 1.2 |
| FAM160A2 | Q8N612 | 1.20E-04 | 0.39 | 1.31 |
| FAM160B1 | Q5W0V3 | 4.80E-06 | 0.45 | 1.36 |
| FAM162A | Q96A26 | 0.0011 | 0.39 | 1.31 |
| FAM173A | Q9BQD7 | 0.013 | 0.5 | 1.42 |
| FAM20B | O75063 | 0.0051 | 0.29 | 1.22 |
| FAM213A | Q9BRX8 | 2.10E-05 | 0.51 | 1.42 |
| FAM45A | Q8TCE6 | 6.20E-04 | 0.42 | 1.34 |
| FAM89B | Q8N5H3 | 0.019 | 0.29 | 1.22 |
| FAS | P25445 | 0.0092 | 0.34 | 1.26 |
| FASTK | Q14296 | 4.00E-05 | 0.6 | 1.51 |
| FBN2 | P35556 | 5.10E-04 | 0.37 | 1.29 |
| FBXL20 | Q96IG2 | 5.10E-04 | 0.32 | 1.25 |
| FBXO21 | O94952 | 0.0039 | 0.31 | 1.24 |
| FBXO3 | Q9UK99 | 2.90E-04 | 0.29 | 1.22 |
| FBXO31 | Q5XUX0 | 0.017 | 0.56 | 1.47 |
| FBXO42 | Q6P3S6 | 2.30E-04 | 0.36 | 1.28 |
| FGFR1 | P11362 | 1.90E-04 | 0.34 | 1.27 |
| FHAD1 | B1AJZ9 | 3.00E-04 | 0.91 | 1.88 |
| FILIP1 | Q7Z7B0 | 6.90E-05 | 1.26 | 2.4 |
| FKBP5 | Q13451 | 9.00E-04 | 0.26 | 1.2 |
| FN3K | Q9H479 | 1.20E-05 | 0.52 | 1.43 |
| FNTA | P49354 | 9.10E-04 | 0.38 | 1.3 |
| FNTB | P49356 | 4.20E-05 | 0.32 | 1.25 |
| FOXI3 | A8MTJ6 | 9.20E-05 | 0.44 | 1.36 |
| FRA10AC1 | Q70Z53 | 3.40E-04 | 0.32 | 1.25 |
| FRRS1L | Q9P0K9 | 0.037 | 0.36 | 1.28 |
| FST | P19883 | 1.70E-04 | 0.55 | 1.46 |
| FUCA1 | P04066 | 6.10E-05 | 0.34 | 1.26 |

| | | | | |
|---|---|---|---|---|
| FZD2 | Q14332 | 0.0076 | 0.42 | 1.34 |
| GAA | P10253 | 7.90E-05 | 0.45 | 1.37 |
| GABPB2 | Q8TAK5 | 0.0016 | 0.34 | 1.26 |
| GAL | P22466 | 1.50E-06 | 0.9 | 1.87 |
| GALC | P54803 | 1.10E-04 | 0.44 | 1.35 |
| GALE | Q14376 | 0.0012 | 0.29 | 1.22 |
| GCA | P28676 | 6.90E-04 | 0.36 | 1.28 |
| GCFC2 | P16383 | 0.0042 | 0.26 | 1.2 |
| GCHFR | P30047 | 0.021 | 0.31 | 1.24 |
| GCLM | P48507 | 0.0028 | 0.28 | 1.22 |
| GGH | Q92820 | 5.90E-05 | 0.71 | 1.64 |
| GGPS1 | O95749 | 7.60E-04 | 0.4 | 1.32 |
| GHITM | Q9H3K2 | 0.0015 | 0.34 | 1.26 |
| GKAP1 | Q5VSY0 | 5.20E-04 | 0.48 | 1.4 |
| GM2A | P17900 | 0.003 | 0.26 | 1.2 |
| GNPDA2 | Q8TDQ7 | 0.0029 | 0.38 | 1.31 |
| GNPNAT1 | Q96EK6 | 6.10E-05 | 0.39 | 1.31 |
| GOLT1B | Q9Y3E0 | 0.012 | 0.29 | 1.22 |
| GPCPD1 | Q9NPB8 | 9.60E-04 | 0.33 | 1.25 |
| GREB1 | Q4ZG55 | 0.0076 | 0.33 | 1.26 |
| GSTK1 | Q9Y2Q3 | 9.00E-05 | 0.34 | 1.27 |
| GTF2E1 | P29083 | 1.50E-04 | 0.26 | 1.2 |
| GYPC | P04921 | 0.0034 | 0.29 | 1.22 |
| H2AFY | O75367 | 7.00E-04 | 0.45 | 1.37 |
| H6PD | O95479 | 4.50E-06 | 0.64 | 1.55 |
| HACD3 | Q9P035 | 1.00E-04 | 0.36 | 1.28 |
| HAGH | Q16775 | 2.60E-04 | 0.32 | 1.25 |
| HDAC8 | Q9BY41 | 3.60E-04 | 0.46 | 1.38 |
| HECA | Q9UBI9 | 2.00E-04 | 0.43 | 1.35 |
| HERPUD2 | Q9BSE4 | 0.014 | 0.27 | 1.2 |
| HEXB | P07686 | 8.80E-04 | 0.29 | 1.22 |
| HIBADH | P31937 | 5.40E-05 | 0.51 | 1.42 |
| HIPK1 | Q86Z02 | 0.023 | 0.5 | 1.42 |
| HIST1H1T | P22492 | 0.029 | 0.27 | 1.2 |
| HIST1H2BD | P58876 | 0.0078 | 0.72 | 1.65 |
| HIST1H2BJ | P06899 | 0.025 | 0.57 | 1.48 |
| HMGB2 | P26583 | 9.40E-06 | 0.68 | 1.6 |
| HRC | P23327 | 0.0073 | 0.77 | 1.71 |
| HS1BP3 | Q53T59 | 5.20E-05 | 0.52 | 1.43 |
| HSD11B2 | P80365 | 1.90E-05 | 0.4 | 1.32 |
| HSP90AA4P | Q58FG1 | 0.0029 | 0.8 | 1.74 |

| | | | | |
|---|---|---|---|---|
| HSP90AB3P | Q58FF7 | 0.014 | 0.43 | 1.35 |
| HSP90AB4P | Q58FF6 | 0.0019 | 0.57 | 1.49 |
| HSPA12A | O43301 | 0.012 | 0.42 | 1.34 |
| HSPB8 | Q9UJY1 | 1.80E-05 | 0.65 | 1.57 |
| HSPBAP1 | Q96EW2 | 2.50E-05 | 0.46 | 1.38 |
| HTRA2 | O43464 | 9.80E-05 | 0.53 | 1.44 |
| IAH1 | Q2TAA2 | 2.00E-04 | 0.35 | 1.27 |
| IARS2 | Q9NSE4 | 7.50E-07 | 0.47 | 1.38 |
| ICAM1 | P05362 | 5.10E-04 | 0.59 | 1.51 |
| IDE | P14735 | 3.20E-04 | 0.41 | 1.33 |
| IDH2 | P48735 | 6.40E-05 | 0.43 | 1.35 |
| IDO1 | P14902 | 1.30E-06 | 0.52 | 1.44 |
| IFI35 | P80217 | 0.0032 | 0.36 | 1.29 |
| IFIT5 | Q13325 | 5.70E-04 | 0.39 | 1.31 |
| IFT57 | Q9NWB7 | 0.0025 | 0.29 | 1.23 |
| IGF2BP2 | Q9Y6M1 | 5.00E-04 | 0.26 | 1.2 |
| IKBKB | O14920 | 1.90E-04 | 0.3 | 1.23 |
| IL27RA | Q6UWB1 | 0.0096 | 0.32 | 1.25 |
| INPP4A | Q96PE3 | 3.50E-04 | 0.38 | 1.3 |
| INPPL1 | O15357 | 6.30E-04 | 0.32 | 1.25 |
| IPPK | Q9H8X2 | 0.027 | 0.41 | 1.33 |
| IRF2 | P14316 | 0.027 | 0.38 | 1.3 |
| IRF7 | Q92985 | 0.0084 | 0.39 | 1.31 |
| ISOC1 | Q96CN7 | 3.60E-04 | 0.33 | 1.25 |
| KATNAL1 | Q9BW62 | 6.30E-05 | 0.34 | 1.26 |
| KBTBD11 | O94819 | 1.10E-04 | 0.6 | 1.52 |
| KCNAB2 | Q13303 | 0.0065 | 0.46 | 1.37 |
| KCNH8 | Q96L42 | 2.30E-04 | 0.71 | 1.63 |
| KHDRBS2 | Q5VWX1 | 4.70E-04 | 0.65 | 1.57 |
| KHNYN | O15037 | 6.80E-04 | 0.31 | 1.24 |
| KIAA0232 | Q92628 | 0.019 | 0.32 | 1.25 |
| KIAA0355 | O15063 | 2.90E-04 | 0.37 | 1.29 |
| KIAA0556 | O60303 | 0.013 | 0.3 | 1.23 |
| KIAA1211 | Q6ZU35 | 1.60E-04 | 0.32 | 1.25 |
| KIFC3 | Q9BVG8 | 9.40E-05 | 0.56 | 1.48 |
| KLF13 | Q9Y2Y9 | 9.80E-04 | 0.27 | 1.21 |
| KNDC1 | Q76NI1 | 0.004 | 0.33 | 1.26 |
| KRTCAP3 | Q53RY4 | 0.0057 | 0.29 | 1.23 |
| KYAT1 | Q16773 | 1.30E-04 | 0.38 | 1.3 |
| KYAT3 | Q6YP21 | 7.70E-05 | 0.42 | 1.34 |
| L2HGDH | Q9H9P8 | 1.30E-04 | 0.32 | 1.24 |

| | | | | |
|---|---|---|---|---|
| LAMP2 | P13473 | 3.60E-04 | 0.3 | 1.23 |
| LARP6 | Q9BRS8 | 1.70E-04 | 0.34 | 1.27 |
| LDAH | Q9H6V9 | 1.70E-04 | 0.29 | 1.22 |
| LDHD | Q86WU2 | 0.0051 | 0.48 | 1.39 |
| LEG1 | Q6P5S2 | 2.30E-06 | 0.47 | 1.38 |
| LEPROT | O15243 | 0.0072 | 0.79 | 1.73 |
| LGALS3BP | Q08380 | 5.40E-07 | 0.72 | 1.65 |
| LGMN | Q99538 | 0.0025 | 0.45 | 1.37 |
| LHFPL5 | Q8TAF8 | 8.10E-04 | 0.46 | 1.38 |
| LHPP | Q9H008 | 0.0015 | 0.53 | 1.45 |
| LIMCH1 | Q9UPQ0 | 0.021 | 0.28 | 1.21 |
| LIN52 | Q52LA3 | 7.10E-04 | 0.33 | 1.26 |
| LOXL3 | P58215 | 4.40E-04 | 0.28 | 1.21 |
| LPGAT1 | Q92604 | 3.30E-04 | 0.46 | 1.37 |
| LPIN2 | Q92539 | 0.0021 | 0.35 | 1.27 |
| LRIG1 | Q96JA1 | 7.60E-05 | 0.3 | 1.23 |
| LRP11 | Q86VZ4 | 0.031 | 0.31 | 1.24 |
| LRRC41 | Q15345 | 0.0011 | 0.26 | 1.2 |
| LRTOMT | Q8WZ04 | 0.024 | 0.42 | 1.34 |
| LSM11 | P83369 | 7.40E-05 | 0.47 | 1.39 |
| LTBP4 | Q8N2S1 | 0.015 | 0.38 | 1.3 |
| LYRM9 | A8MSI8 | 0.023 | 0.57 | 1.48 |
| MACC1 | Q6ZN28 | 7.40E-05 | 0.78 | 1.71 |
| MACROD1 | Q9BQ69 | 1.80E-06 | 0.63 | 1.55 |
| MAL2 | Q969L2 | 0.0029 | 0.26 | 1.2 |
| MAP1LC3A | Q9H492 | 1.20E-04 | 0.53 | 1.45 |
| MAP1LC3B | Q9GZQ8 | 0.0066 | 0.27 | 1.21 |
| MAP2K5 | Q13163 | 5.30E-05 | 0.45 | 1.37 |
| MAP4K2 | Q12851 | 0.0055 | 0.31 | 1.24 |
| MAP7D1 | Q3KQU3 | 0.0094 | 0.26 | 1.2 |
| MAP7D2 | Q96T17 | 0.019 | 0.33 | 1.26 |
| MAPK8IP3 | Q9UPT6 | 0.0027 | 0.44 | 1.35 |
| MARCH2 | Q969Z3 | 3.20E-05 | 0.4 | 1.32 |
| MARCKS | P29966 | 0.012 | 0.31 | 1.24 |
| MAT1A | Q00266 | 0.0074 | 0.3 | 1.23 |
| MBLAC2 | Q68D91 | 0.0024 | 0.33 | 1.25 |
| MCM9 | Q9NXL9 | 0.0083 | 0.27 | 1.2 |
| MCPH1 | Q8NEM0 | 0.0099 | 0.3 | 1.23 |
| MDK | P21741 | 0.003 | 0.41 | 1.33 |
| ME1 | P48163 | 0.0038 | 0.33 | 1.26 |
| MED12L | Q86YW9 | 0.028 | 0.38 | 1.3 |

| | | | | |
|---|---|---|---|---|
| MED30 | Q96HR3 | 3.00E-04 | 0.28 | 1.21 |
| MED31 | Q9Y3C7 | 0.014 | 0.46 | 1.37 |
| MEGF8 | Q7Z7M0 | 5.50E-04 | 0.27 | 1.2 |
| MEIOC | A2RUB1 | 0.001 | 0.35 | 1.27 |
| METTL5 | Q9NRN9 | 2.30E-04 | 0.42 | 1.34 |
| MGARP | Q8TDB4 | 9.80E-04 | 0.67 | 1.59 |
| MGMT | P16455 | 1.90E-04 | 0.45 | 1.36 |
| MIF4GD | A9UHW6 | 3.40E-04 | 0.32 | 1.25 |
| MINOS1 | Q5TGZ0 | 0.034 | 0.35 | 1.28 |
| MIPOL1 | Q8TD10 | 3.20E-04 | 0.29 | 1.22 |
| MLTK_HUMAN | NA | 6.60E-04 | 0.29 | 1.22 |
| MMS22L | Q6ZRQ5 | 2.30E-05 | 0.44 | 1.36 |
| MOB1A | Q9H8S9 | 7.60E-04 | 0.26 | 1.2 |
| MOB3B | Q86TA1 | 0.0011 | 0.42 | 1.34 |
| MOCOS | Q96EN8 | 0.0048 | 0.42 | 1.34 |
| MOCS2 | O96007 | 4.00E-04 | 0.3 | 1.23 |
| MON1B | Q7L1V2 | 0.018 | 0.31 | 1.24 |
| MSI1 | O43347 | 0.0042 | 0.41 | 1.32 |
| MSI2 | Q96DH6 | 9.60E-04 | 0.39 | 1.31 |
| MSN | P26038 | 2.70E-04 | 0.32 | 1.25 |
| MTBP | Q96DY7 | 0.0025 | 0.33 | 1.26 |
| MTFMT | Q96DP5 | 0.016 | 0.26 | 1.2 |
| MTG2 | Q9H4K7 | 4.50E-04 | 0.29 | 1.23 |
| MXRA7 | P84157 | 3.80E-04 | 0.49 | 1.4 |
| MYNN | Q9NPC7 | 0.0073 | 0.45 | 1.37 |
| N6AMT1 | Q9Y5N5 | 0.0037 | 0.27 | 1.21 |
| N6MT2_HUMAN | NA | 0.005 | 0.68 | 1.6 |
| NAB2 | Q15742 | 8.20E-05 | 0.45 | 1.37 |
| NADK | O95544 | 1.20E-04 | 0.39 | 1.31 |
| NADK2 | Q4G0N4 | 2.80E-05 | 0.4 | 1.32 |
| NAGA | P17050 | 7.70E-04 | 0.32 | 1.25 |
| NALCN | Q8IZF0 | 0.014 | 0.55 | 1.47 |
| NAPB | Q9H115 | 0.0023 | 0.27 | 1.2 |
| NAT1 | P18440 | 0.0036 | 0.27 | 1.21 |
| NAT9 | Q9BTE0 | 0.0073 | 0.42 | 1.34 |
| NAXE | Q8NCW5 | 1.30E-05 | 0.45 | 1.36 |
| NBEA | Q8NFP9 | 4.20E-04 | 0.38 | 1.3 |
| NCOA7 | Q8NI08 | 0.0052 | 0.32 | 1.25 |
| NCS1 | P62166 | 0.029 | 0.34 | 1.27 |
| NDUFAF8 | A1L188 | 0.025 | 0.29 | 1.22 |
| NECAP2 | Q9NVZ3 | 0.013 | 0.29 | 1.22 |

| | | | | |
|---|---|---|---|---|
| NEK9 | Q8TD19 | 0.0012 | 0.27 | 1.21 |
| NFKB1 | P19838 | 7.50E-05 | 0.35 | 1.27 |
| NID1 | P14543 | 0.0022 | 0.59 | 1.5 |
| NLN | Q9BYT8 | 3.90E-05 | 0.29 | 1.22 |
| NLRX1 | Q86UT6 | 4.40E-06 | 0.42 | 1.34 |
| NME2P1 | O60361 | 2.70E-04 | 1.15 | 2.23 |
| NME4 | O00746 | 3.10E-04 | 0.47 | 1.39 |
| NOS1AP | O75052 | 0.0019 | 0.26 | 1.2 |
| NR2C2AP | Q86WQ0 | 0.002 | 0.26 | 1.2 |
| NR3C1 | P04150 | 0.0047 | 0.43 | 1.35 |
| NR6A1 | Q15406 | 0.018 | 0.29 | 1.22 |
| NRF1 | Q16656 | 0.002 | 0.28 | 1.21 |
| NSMCE1 | Q8WV22 | 6.10E-05 | 0.38 | 1.3 |
| NSMCE2 | Q96MF7 | 0.001 | 0.31 | 1.24 |
| NSMCE3 | Q96MG7 | 4.70E-05 | 0.35 | 1.28 |
| NSMCE4A | Q9NXX6 | 7.30E-04 | 0.3 | 1.23 |
| NT5DC1 | Q5TFE4 | 0.0012 | 0.26 | 1.2 |
| NUB1 | Q9Y5A7 | 4.30E-05 | 0.36 | 1.28 |
| NUCKS1 | Q9H1E3 | 8.20E-04 | 0.69 | 1.62 |
| NUDT11 | Q96G61 | 0.0051 | 0.26 | 1.2 |
| NUDT17 | P0C025 | 0.016 | 0.84 | 1.8 |
| NUDT6 | P53370 | 4.40E-04 | 0.44 | 1.36 |
| OARD1 | Q9Y530 | 0.0089 | 0.28 | 1.22 |
| OAT | P04181 | 0.0013 | 0.29 | 1.22 |
| OPLAH | O14841 | 2.00E-04 | 0.31 | 1.24 |
| ORC3 | Q9UBD5 | 0.0041 | 0.26 | 1.2 |
| OSBPL1A | Q9BXW6 | 0.0025 | 0.34 | 1.27 |
| OTX2 | P32243 | 0.006 | 0.57 | 1.48 |
| OXLD1 | Q5BKU9 | 0.02 | 0.31 | 1.24 |
| P3H4 | Q92791 | 5.00E-04 | 0.27 | 1.2 |
| PAAF1 | Q9BRP4 | 2.00E-05 | 0.42 | 1.34 |
| PAFAH1B1 | P43034 | 3.50E-04 | 0.32 | 1.24 |
| PAFAH1B2 | P68402 | 0.0022 | 0.29 | 1.22 |
| PAPD4 | Q6PIY7 | 0.003 | 0.27 | 1.2 |
| PARP12 | Q9H0J9 | 0.0019 | 0.36 | 1.28 |
| PAX6 | P26367 | 4.00E-05 | 1.26 | 2.39 |
| PBK | Q96KB5 | 2.30E-04 | 0.33 | 1.26 |
| PCCB | P05166 | 0.0016 | 0.31 | 1.24 |
| PCDH11X | Q9BZA7 | 1.10E-04 | 1.37 | 2.58 |
| PCGF3 | Q3KNV8 | 0.001 | 0.7 | 1.63 |
| PCK2 | Q16822 | 1.80E-06 | 0.76 | 1.69 |

| | | | | |
|---|---|---|---|---|
| PCYOX1 | Q9UHG3 | 1.60E-04 | 0.35 | 1.27 |
| PDCD4 | Q53EL6 | 6.70E-07 | 0.71 | 1.64 |
| PDE6D | O43924 | 0.0088 | 0.26 | 1.2 |
| PDGFA | P04085 | 9.40E-06 | 2.27 | 4.81 |
| PDK2 | Q15119 | 0.0012 | 0.27 | 1.21 |
| PDS5A | Q29RF7 | 0.0052 | 0.26 | 1.2 |
| PEX14 | O75381 | 4.00E-04 | 0.28 | 1.21 |
| PFKFB2 | O60825 | 3.20E-04 | 0.26 | 1.2 |
| PHGDH | O43175 | 7.40E-04 | 0.37 | 1.3 |
| PHYH | O14832 | 6.90E-04 | 0.48 | 1.4 |
| PHYHIP | Q92561 | 8.40E-04 | 0.29 | 1.22 |
| PIK3R6 | Q5UE93 | 0.0011 | 0.43 | 1.34 |
| PIR | O00625 | 2.80E-04 | 0.56 | 1.47 |
| PKIA | P61925 | 4.40E-04 | 0.42 | 1.33 |
| PKIB | Q9C010 | 1.90E-04 | 0.55 | 1.47 |
| PKLR | P30613 | 0.047 | 0.54 | 1.45 |
| PLCG2 | P16885 | 1.50E-05 | 0.48 | 1.39 |
| PLCL2 | Q9UPR0 | 1.30E-06 | 0.58 | 1.49 |
| PLCXD1 | Q9NUJ7 | 0.0016 | 0.31 | 1.24 |
| PLEKHA4 | Q9H4M7 | 0.0095 | 0.49 | 1.4 |
| PLGRKT | Q9HBL7 | 1.50E-04 | 0.31 | 1.24 |
| PLIN2 | Q99541 | 2.60E-06 | 0.79 | 1.73 |
| PLPP6 | Q8IY26 | 0.0018 | 0.38 | 1.3 |
| PLSCR1 | O15162 | 0.0033 | 0.68 | 1.6 |
| PLSCR3 | Q9NRY6 | 0.001 | 0.32 | 1.25 |
| PLXND1 | Q9Y4D7 | 3.10E-04 | 0.45 | 1.37 |
| PM20D2 | Q8IYS1 | 7.20E-04 | 0.32 | 1.25 |
| PNPLA2 | Q96AD5 | 0.0015 | 0.3 | 1.23 |
| PNPLA8 | Q9NP80 | 0.0044 | 0.32 | 1.24 |
| PNPO | Q9NVS9 | 3.70E-07 | 0.84 | 1.79 |
| POLQ | O75417 | 0.0013 | 0.3 | 1.23 |
| POLR2M | P0CAP2 | 0.0017 | 0.26 | 1.2 |
| POMP | Q9Y244 | 8.00E-05 | 0.52 | 1.44 |
| PPAT | Q06203 | 0.0015 | 0.3 | 1.23 |
| PPCS | Q9HAB8 | 3.40E-05 | 0.32 | 1.25 |
| PPDPF | Q9H3Y8 | 6.90E-04 | 0.3 | 1.23 |
| PPIP5K1 | Q6PFW1 | 0.026 | 0.57 | 1.49 |
| PPM1F | P49593 | 6.20E-04 | 0.29 | 1.22 |
| PPP1R9B | Q96SB3 | 0.0018 | 0.31 | 1.24 |
| PPP4R4 | Q6NUP7 | 0.00093 | 0.4 | 1.32 |
| PPT2 | Q9UMR5 | 2.20E-04 | 0.3 | 1.23 |

| | | | | |
|---|---|---|---|---|
| PRDX4 | Q13162 | 5.40E-06 | 0.52 | 1.43 |
| PREX1 | Q8TCU6 | 5.10E-04 | 0.42 | 1.34 |
| PRICKLE1 | Q96MT3 | 2.60E-04 | 0.98 | 1.98 |
| PRMT3 | O60678 | 0.0019 | 0.29 | 1.23 |
| PRPF40B | Q6NWY9 | 0.0033 | 0.32 | 1.25 |
| PRRT3 | Q5FWE3 | 0.023 | 0.44 | 1.36 |
| PRSS1 | P07477 | 0.0047 | 0.6 | 1.51 |
| PRSS8 | Q16651 | 5.30E-05 | 0.44 | 1.36 |
| PSAP | P07602 | 0.009 | 0.42 | 1.34 |
| PSAT1 | Q9Y617 | 5.70E-05 | 0.72 | 1.65 |
| PSMB10 | P40306 | 0.0034 | 0.3 | 1.23 |
| PSMB8 | P28062 | 0.0015 | 0.27 | 1.21 |
| PSMB9 | P28065 | 0.0089 | 0.32 | 1.25 |
| PSMC3IP | Q9P2W1 | 0.0041 | 0.26 | 1.2 |
| PSPH | P78330 | 1.50E-04 | 0.66 | 1.58 |
| PTBP2 | Q9UKA9 | 6.20E-05 | 0.38 | 1.3 |
| PTGR1 | Q14914 | 1.50E-05 | 0.5 | 1.41 |
| PTGR2 | Q8N8N7 | 2.50E-04 | 0.39 | 1.31 |
| PTMA | P06454 | 0.03 | 0.43 | 1.35 |
| PTMS | P20962 | 8.80E-04 | 0.66 | 1.58 |
| PTP4A1 | Q93096 | 0.0054 | 0.26 | 1.2 |
| PTPRR | Q15256 | 0.0041 | 0.56 | 1.47 |
| PXK | Q7Z7A4 | 4.90E-04 | 0.38 | 1.3 |
| PYCARD | Q9ULZ3 | 1.40E-04 | 0.41 | 1.33 |
| PYCR1 | P32322 | 8.30E-06 | 0.55 | 1.46 |
| PYROXD1 | Q8WU10 | 0.023 | 0.38 | 1.3 |
| QRICH1 | Q2TAL8 | 0.002 | 0.32 | 1.25 |
| RAB12 | Q6IQ22 | 1.70E-04 | 0.28 | 1.22 |
| RAB17 | Q9H0T7 | 5.30E-04 | 0.46 | 1.37 |
| RAB19 | A4D1S5 | 0.0053 | 0.28 | 1.22 |
| RAB23 | Q9ULC3 | 7.30E-05 | 0.37 | 1.29 |
| RAB24 | Q969Q5 | 6.00E-04 | 0.32 | 1.25 |
| RAB25 | P57735 | 0.0046 | 0.27 | 1.21 |
| RAB27A | P51159 | 0.0012 | 0.39 | 1.31 |
| RAB3C | Q96E17 | 1.10E-04 | 0.42 | 1.34 |
| RAB42 | Q8N4Z0 | 0.001 | 0.32 | 1.25 |
| RAB4B | P61018 | 0.038 | 0.29 | 1.23 |
| RAD54B | Q9Y620 | 1.90E-04 | 0.33 | 1.26 |
| RAD9A | Q99638 | 0.0089 | 0.35 | 1.28 |
| RAP1GAP | P47736 | 0.0037 | 0.36 | 1.29 |
| RASA2 | Q15283 | 1.50E-04 | 0.26 | 1.2 |

| | | | | |
|---|---|---|---|---|
| RBAKDN | A6NC62 | 1.80E-04 | 0.44 | 1.35 |
| RBM20 | Q5T481 | 0.05 | 0.42 | 1.34 |
| RBSN | Q9H1K0 | 5.70E-05 | 0.27 | 1.2 |
| RCN1 | Q15293 | 0.0013 | 0.3 | 1.23 |
| RCOR1 | Q9UKL0 | 1.70E-04 | 0.26 | 1.2 |
| RDX | P35241 | 0.004 | 0.37 | 1.29 |
| RHOB | P62745 | 2.40E-04 | 0.46 | 1.38 |
| RHPN2 | Q8IUC4 | 2.10E-05 | 0.39 | 1.31 |
| RIMKLB | Q9ULI2 | 0.015 | 0.28 | 1.21 |
| RIMS3 | Q9UJD0 | 0.022 | 0.41 | 1.33 |
| RNASET2 | O00584 | 0.016 | 0.27 | 1.2 |
| RNF10 | Q8N5U6 | 4.20E-05 | 0.53 | 1.44 |
| RNF123 | Q5XPI4 | 2.30E-04 | 0.28 | 1.21 |
| RNF170 | Q96K19 | 7.10E-04 | 0.41 | 1.33 |
| RNMT | O43148 | 9.50E-05 | 0.27 | 1.21 |
| ROS1 | P08922 | 2.40E-04 | 0.41 | 1.33 |
| RPRD1A | Q96P16 | 1.50E-04 | 0.31 | 1.24 |
| RPS6KA3 | P51812 | 0.001 | 0.28 | 1.21 |
| RPS6KA4 | O75676 | 4.80E-04 | 0.26 | 1.2 |
| RPS6KA5 | O75582 | 8.80E-08 | 0.66 | 1.58 |
| RUFY3 | Q7L099 | 2.50E-04 | 0.31 | 1.24 |
| RXRB | P28702 | 0.0038 | 0.36 | 1.28 |
| S100A11 | P31949 | 0.0046 | 0.3 | 1.23 |
| S100A13 | Q99584 | 0.01 | 0.71 | 1.64 |
| S100A16 | Q96FQ6 | 0.0013 | 0.36 | 1.28 |
| S100A4 | P26447 | 3.30E-04 | 0.52 | 1.43 |
| SARG | Q9BW04 | 7.80E-04 | 0.32 | 1.25 |
| SARS | P49591 | 0.001 | 0.3 | 1.23 |
| SAT2 | Q96F10 | 7.70E-05 | 0.6 | 1.52 |
| SBF2 | Q86WG5 | 0.0041 | 0.26 | 1.2 |
| SCAPER | Q9BY12 | 2.00E-04 | 0.44 | 1.36 |
| SCARB2 | Q14108 | 1.80E-04 | 0.5 | 1.41 |
| SCCPDH | Q8NBX0 | 4.20E-05 | 0.38 | 1.3 |
| SCLY | Q96I15 | 0.0096 | 0.31 | 1.24 |
| SCPEP1 | Q9HB40 | 0.0068 | 0.29 | 1.22 |
| SCYL3 | Q8IZE3 | 5.80E-04 | 0.36 | 1.28 |
| SDHAF4 | Q5VUM1 | 0.0042 | 0.32 | 1.25 |
| SDHC | Q99643 | 1.70E-04 | 0.45 | 1.36 |
| SEC61A2 | Q9H9S3 | 0.0018 | 0.27 | 1.2 |
| SERAC1 | Q96JX3 | 0.0011 | 0.32 | 1.25 |
| SESN2 | P58004 | 0.033 | 0.71 | 1.64 |

| | | | | |
|---|---|---|---|---|
| SESN3 | P58005 | 0.0015 | 0.32 | 1.25 |
| SFRP1 | Q8N474 | 3.30E-04 | 0.57 | 1.48 |
| SFRP2 | Q96HF1 | 0.0063 | 0.28 | 1.22 |
| SGCG | Q13326 | 0.008 | 0.6 | 1.52 |
| SH3BGRL2 | Q9UJC5 | 0.0016 | 0.37 | 1.29 |
| SH3TC1 | Q8TE82 | 0.042 | 0.27 | 1.2 |
| SH3TC2 | Q8TF17 | 0.0055 | 0.32 | 1.25 |
| SHMT1 | P34896 | 0.011 | 0.26 | 1.2 |
| SHMT2 | P34897 | 4.20E-05 | 0.44 | 1.35 |
| SIX5 | Q8N196 | 0.0036 | 0.31 | 1.24 |
| SLC11A2 | P49281 | 1.80E-06 | 0.74 | 1.67 |
| SLC1A3 | P43003 | 4.50E-04 | 0.75 | 1.68 |
| SLC25A21 | Q9BQT8 | 3.20E-04 | 0.26 | 1.2 |
| SLC25A36 | Q96CQ1 | 5.20E-04 | 0.26 | 1.2 |
| SLC27A6 | Q9Y2P4 | 4.80E-04 | 0.35 | 1.27 |
| SLC29A3 | Q9BZD2 | 8.10E-04 | 0.3 | 1.23 |
| SLC2A12 | Q8TD20 | 9.60E-04 | 0.75 | 1.68 |
| SLC2A13 | Q96QE2 | 0.034 | 0.27 | 1.21 |
| SLC2A4 | P14672 | 2.70E-04 | 0.83 | 1.78 |
| SLC35F6 | Q8N357 | 1.10E-05 | 0.41 | 1.33 |
| SLC37A4 | O43826 | 0.0014 | 0.36 | 1.28 |
| SLC44A2 | Q8IWA5 | 1.30E-06 | 0.83 | 1.78 |
| SLC4A7 | Q9Y6M7 | 0.026 | 0.26 | 1.2 |
| SLC6A9 | P48067 | 8.10E-05 | 0.51 | 1.42 |
| SLC7A11 | Q9UPY5 | 1.50E-05 | 0.83 | 1.78 |
| SLF1 | Q9BQI6 | 0.028 | 0.28 | 1.21 |
| SLFN13 | Q68D06 | 1.30E-04 | 0.42 | 1.34 |
| SLIT2 | O94813 | 0.029 | 0.33 | 1.25 |
| SLX4IP | Q5VYV7 | 0.0021 | 0.31 | 1.24 |
| SMAD5 | Q99717 | 7.60E-04 | 0.26 | 1.2 |
| SMARCAL1 | Q9NZC9 | 8.10E-04 | 0.3 | 1.23 |
| SMC5 | Q8IY18 | 1.10E-04 | 0.32 | 1.25 |
| SMCHD1 | A6NHR9 | 2.10E-05 | 0.38 | 1.3 |
| SMCO4 | Q9NRQ5 | 0.0077 | 0.3 | 1.23 |
| SMDT1 | Q9H4I9 | 0.0044 | 0.38 | 1.3 |
| SMIM19 | Q96E16 | 0.0014 | 0.39 | 1.31 |
| SMIM20 | Q8N5G0 | 0.018 | 0.31 | 1.24 |
| SMIM4 | Q8WVI0 | 1.10E-04 | 0.56 | 1.48 |
| SMOC2 | Q9H3U7 | 0.034 | 0.3 | 1.23 |
| SMPDL3A | Q92484 | 0.0016 | 0.75 | 1.68 |
| SMS | P52788 | 1.40E-04 | 0.39 | 1.31 |

| | | | | |
|---|---|---|---|---|
| SMURF2 | Q9HAU4 | 0.0028 | 0.27 | 1.21 |
| SMYD3 | Q9H7B4 | 7.50E-04 | 0.27 | 1.2 |
| SNCG | O76070 | 0.0011 | 0.36 | 1.28 |
| SNF8 | Q96H20 | 0.0095 | 0.26 | 1.2 |
| SNRK | Q9NRH2 | 0.0072 | 0.31 | 1.24 |
| SNRNP35 | Q16560 | 1.20E-04 | 0.32 | 1.25 |
| SNX30 | Q5VWJ9 | 2.50E-04 | 0.38 | 1.3 |
| SNX4 | O95219 | 3.30E-05 | 0.31 | 1.24 |
| SNX7 | Q9UNH6 | 6.20E-04 | 0.31 | 1.24 |
| SOAT1 | P35610 | 5.80E-06 | 0.96 | 1.95 |
| SOCS2 | O14508 | 6.60E-04 | 0.61 | 1.53 |
| SOCS6 | O14544 | 0.0064 | 0.45 | 1.37 |
| SOX21 | Q9Y651 | 5.40E-05 | 0.37 | 1.29 |
| SPATS2 | Q86XZ4 | 3.20E-05 | 0.37 | 1.3 |
| SPG21 | Q9NZD8 | 0.0016 | 0.26 | 1.2 |
| SRSF12 | Q8WXF0 | 0.0012 | 0.36 | 1.29 |
| SRSF4 | Q08170 | 5.70E-04 | 0.26 | 1.2 |
| SSBP2 | P81877 | 0.0042 | 0.34 | 1.27 |
| SSBP4 | Q9BWG4 | 0.026 | 0.27 | 1.2 |
| SSH3 | Q8TE77 | 3.10E-05 | 0.36 | 1.28 |
| SSTR2 | P30874 | 1.30E-04 | 0.47 | 1.39 |
| STAP2 | Q9UGK3 | 2.50E-04 | 0.44 | 1.36 |
| STAT1 | P42224 | 2.00E-04 | 0.31 | 1.24 |
| STAT5B | P51692 | 1.40E-04 | 0.32 | 1.25 |
| STAT6 | P42226 | 0.025 | 0.36 | 1.28 |
| STC2 | O76061 | 1.40E-05 | 0.9 | 1.87 |
| STK17B | O94768 | 3.30E-04 | 0.39 | 1.31 |
| STK3 | Q13188 | 2.90E-04 | 0.34 | 1.26 |
| STK39 | Q9UEW8 | 0.0015 | 0.28 | 1.22 |
| STN1 | Q9H668 | 0.026 | 0.27 | 1.21 |
| STON1 | Q9Y6Q2 | 0.0011 | 0.39 | 1.31 |
| STX12 | Q86Y82 | 2.60E-04 | 0.28 | 1.21 |
| STX17 | P56962 | 1.50E-04 | 0.27 | 1.2 |
| SULF1 | Q8IWU6 | 0.025 | 0.32 | 1.24 |
| SULT1A1 | P50225 | 0.0084 | 0.67 | 1.59 |
| SUN2 | Q9UH99 | 9.60E-05 | 0.39 | 1.31 |
| SVIL | O95425 | 2.80E-05 | 0.37 | 1.29 |
| SYG_HUMAN | NA | 0.0052 | 0.27 | 1.2 |
| SYN3 | O14994 | 0.0012 | 0.48 | 1.4 |
| SYNE1 | Q8NF91 | 5.60E-05 | 0.68 | 1.6 |
| SYNE3 | Q6ZMZ3 | 0.013 | 0.64 | 1.56 |

| | | | | |
|---|---|---|---|---|
| SYNE4 | Q8N205 | 0.016 | 0.27 | 1.21 |
| SYNGR2 | O43760 | 0.0011 | 0.27 | 1.21 |
| SYNGR3 | O43761 | 2.10E-04 | 0.4 | 1.32 |
| SYNJ2 | O15056 | 0.0021 | 0.32 | 1.25 |
| SYPL1 | Q16563 | 0.0016 | 0.32 | 1.25 |
| SYT7 | O43581 | 0.011 | 0.37 | 1.29 |
| SZT2 | Q5T011 | 0.034 | 0.28 | 1.22 |
| TARSL2 | A2RTX5 | 2.20E-04 | 0.37 | 1.3 |
| TATDN2 | Q93075 | 0.019 | 0.34 | 1.26 |
| TBC1D20 | Q96BZ9 | 0.043 | 0.29 | 1.22 |
| TBC1D25 | Q3MII6 | 0.018 | 0.38 | 1.3 |
| TBL2 | Q9Y4P3 | 4.90E-04 | 0.29 | 1.22 |
| TCAF1 | Q9Y4C2 | 2.60E-04 | 0.28 | 1.22 |
| TCEAL4 | Q96EI5 | 9.60E-04 | 0.28 | 1.22 |
| TCHP | Q9BT92 | 0.0033 | 0.49 | 1.41 |
| TCIRG1 | Q13488 | 0.019 | 0.3 | 1.23 |
| TEAD1 | P28347 | 1.60E-04 | 0.63 | 1.55 |
| TEAD3 | Q99594 | 1.50E-04 | 0.55 | 1.46 |
| TERF2 | Q15554 | 1.10E-05 | 0.42 | 1.34 |
| TESK2 | Q96S53 | 0.0046 | 0.49 | 1.41 |
| TFEB | P19484 | 0.043 | 0.71 | 1.63 |
| TFEC | O14948 | 0.012 | 0.57 | 1.48 |
| TGM3 | Q08188 | 0.0077 | 0.52 | 1.43 |
| THAP12 | O43422 | 6.70E-05 | 0.3 | 1.23 |
| THUMPD2 | Q9BTF0 | 1.30E-04 | 0.36 | 1.28 |
| TKFC | Q3LXA3 | 2.10E-04 | 0.32 | 1.25 |
| TMED1 | Q13445 | 3.20E-04 | 0.3 | 1.23 |
| TMED8 | Q6PL24 | 0.0028 | 0.26 | 1.2 |
| TMEM120A | Q9BXJ8 | 0.001 | 0.59 | 1.51 |
| TMEM160 | Q9NX00 | 4.40E-04 | 0.36 | 1.28 |
| TMEM164 | Q5U3C3 | 0.0024 | 0.48 | 1.4 |
| TMEM179B | Q7Z7N9 | 0.032 | 0.31 | 1.24 |
| TMEM181 | Q9P2C4 | 7.40E-04 | 0.27 | 1.2 |
| TMEM184C | Q9NVA4 | 0.0043 | 0.36 | 1.28 |
| TMEM26 | Q6ZUK4 | 0.0089 | 0.59 | 1.5 |
| TMEM267 | Q0VDI3 | 0.0038 | 0.47 | 1.38 |
| TMEM268 | Q5VZI3 | 8.10E-04 | 0.31 | 1.24 |
| TMEM35B | Q8NCS4 | 0.0037 | 0.35 | 1.28 |
| TMEM65 | Q6PI78 | 1.00E-04 | 0.32 | 1.25 |
| TMEM9 | Q9P0T7 | 0.0026 | 0.28 | 1.22 |
| TMLHE | Q9NVH6 | 3.00E-06 | 0.57 | 1.48 |

| | | | | |
|---|---|---|---|---|
| TMSB4X | P62328 | 0.0014 | 0.34 | 1.26 |
| TMTC4 | Q5T4D3 | 0.029 | 0.28 | 1.21 |
| TNFAIP2 | Q03169 | 9.50E-08 | 0.99 | 1.98 |
| TNFRSF11A | Q9Y6Q6 | 0.0025 | 0.9 | 1.86 |
| TOM1L1 | O75674 | 9.70E-04 | 0.3 | 1.23 |
| TOMM40L | Q969M1 | 0.002 | 0.38 | 1.3 |
| TOPAZ1 | Q8N9V7 | 0.018 | 0.71 | 1.63 |
| TPP1 | O14773 | 3.30E-04 | 0.36 | 1.28 |
| TPPP3 | Q9BW30 | 0.0028 | 0.74 | 1.67 |
| TRAPPC3 | O43617 | 4.10E-05 | 0.48 | 1.4 |
| TRAPPC5 | Q8IUR0 | 0.0025 | 0.3 | 1.23 |
| TRAPPC6A | O75865 | 4.80E-06 | 0.66 | 1.58 |
| TRIM25 | Q14258 | 4.30E-04 | 0.27 | 1.21 |
| TRIM47 | Q96LD4 | 3.00E-05 | 0.31 | 1.24 |
| TRIM72 | Q6ZMU5 | 1.60E-05 | 0.64 | 1.56 |
| TRIQK | Q629K1 | 0.0012 | 0.54 | 1.45 |
| TRMT12 | Q53H54 | 0.0095 | 0.39 | 1.31 |
| TRNT1 | Q96Q11 | 3.20E-04 | 0.27 | 1.21 |
| TSC1 | Q92574 | 5.40E-04 | 0.29 | 1.22 |
| TSPAN3 | O60637 | 7.90E-05 | 0.45 | 1.37 |
| TST | Q16762 | 1.20E-04 | 0.3 | 1.23 |
| TSTD1 | Q8NFU3 | 0.0032 | 0.42 | 1.33 |
| TTC12 | Q9H892 | 1.90E-05 | 1.11 | 2.15 |
| TTC38 | Q5R3I4 | 3.60E-04 | 0.37 | 1.29 |
| TTLL10 | Q6ZVT0 | 3.50E-04 | 1.13 | 2.19 |
| TUBAL3 | A6NHL2 | 2.50E-06 | 1.01 | 2.02 |
| TUSC3 | Q13454 | 1.80E-05 | 0.48 | 1.4 |
| TXNRD2 | Q9NNW7 | 6.50E-04 | 0.28 | 1.21 |
| TXNRD3 | Q86VQ6 | 3.70E-07 | 0.8 | 1.75 |
| UAP1L1 | Q3KQV9 | 0.017 | 0.35 | 1.28 |
| UBE2D1 | P51668 | 0.009 | 0.29 | 1.22 |
| UBE2D3 | P61077 | 0.042 | 0.31 | 1.24 |
| UBE2E2 | Q96LR5 | 0.042 | 0.54 | 1.45 |
| UBE2E3 | Q969T4 | 0.0017 | 1.31 | 2.49 |
| UBE2G1 | P62253 | 8.40E-04 | 0.37 | 1.29 |
| UBE2K | P61086 | 0.003 | 0.28 | 1.21 |
| UBE2T | Q9NPD8 | 0.002 | 0.32 | 1.25 |
| UBOX5 | O94941 | 4.70E-04 | 0.37 | 1.29 |
| UGP2 | Q16851 | 7.00E-04 | 0.3 | 1.23 |
| UHRF1BP1 | Q6BDS2 | 2.80E-05 | 0.42 | 1.34 |
| UHRF2 | Q96PU4 | 0.0023 | 0.34 | 1.27 |

| | | | | |
|---|---|---|---|---|
| UNC50 | Q53HI1 | 0.035 | 0.27 | 1.2 |
| UNK | Q9C0B0 | 0.0065 | 0.33 | 1.26 |
| USP24 | Q9UPU5 | 2.90E-04 | 0.26 | 1.2 |
| USP3 | Q9Y6I4 | 3.20E-05 | 0.75 | 1.68 |
| UTY | O14607 | 7.80E-04 | 0.39 | 1.31 |
| VAMP2 | P63027 | 0.014 | 0.26 | 1.2 |
| VAMP4 | O75379 | 0.0026 | 0.37 | 1.29 |
| VARS2 | Q5ST30 | 2.00E-04 | 0.34 | 1.27 |
| VAT1L | Q9HCJ6 | 0.0056 | 0.33 | 1.26 |
| VAV1 | P15498 | 0.032 | 0.54 | 1.46 |
| VCPKMT | Q9H867 | 3.90E-04 | 0.27 | 1.2 |
| VGF | O15240 | 4.60E-06 | 1.22 | 2.32 |
| VGLL3 | A8MV65 | 0.002 | 0.47 | 1.38 |
| VIPAS39 | Q9H9C1 | 1.90E-04 | 0.27 | 1.21 |
| VPS45 | Q9NRW7 | 4.70E-04 | 0.29 | 1.22 |
| VWA1 | Q6PCB0 | 0.008 | 0.27 | 1.21 |
| VWA3B | Q502W6 | 1.50E-05 | 1.06 | 2.09 |
| WARS | P23381 | 0.0034 | 0.35 | 1.27 |
| WASF3 | Q9UPY6 | 0.012 | 0.27 | 1.21 |
| WDR7 | Q9Y4E6 | 5.60E-04 | 0.27 | 1.21 |
| WDTC1 | Q8N5D0 | 0.0032 | 0.5 | 1.42 |
| WIPI2 | Q9Y4P8 | 0.0011 | 0.26 | 1.2 |
| WTIP | A6NIX2 | 0.0049 | 0.28 | 1.21 |
| XPA | P23025 | 9.00E-04 | 0.3 | 1.23 |
| XPOT | O43592 | 4.20E-05 | 0.4 | 1.32 |
| YAF2 | Q8IY57 | 0.0054 | 0.67 | 1.59 |
| YARS | P54577 | 8.90E-04 | 0.35 | 1.28 |
| YIPF2 | Q9BWQ6 | 0.026 | 0.47 | 1.39 |
| ZBED8 | Q8IZ13 | 6.20E-04 | 0.42 | 1.34 |
| ZBTB10 | Q96DT7 | 5.50E-05 | 0.46 | 1.38 |
| ZBTB33 | Q86T24 | 6.70E-04 | 0.37 | 1.29 |
| ZBTB43 | O43298 | 0.017 | 0.26 | 1.2 |
| ZBTB45 | Q96K62 | 0.0087 | 0.46 | 1.37 |
| ZC3H6 | P61129 | 9.00E-04 | 0.44 | 1.36 |
| ZCCHC4 | Q9H5U6 | 0.0082 | 0.27 | 1.2 |
| ZDHHC4 | Q9NPG8 | 7.70E-04 | 0.62 | 1.54 |
| ZFHX2 | Q9C0A1 | 5.70E-04 | 0.31 | 1.24 |
| ZFX | P17010 | 0.0022 | 0.29 | 1.22 |
| ZFYVE9 | O95405 | 2.60E-04 | 0.28 | 1.21 |
| ZHX2 | Q9Y6X8 | 4.40E-04 | 0.32 | 1.24 |
| ZIC2 | O95409 | 4.80E-04 | 0.5 | 1.41 |

| | | | | |
|---|---|---|---|---|
| ZIC5 | Q96T25 | 2.00E-04 | 0.42 | 1.33 |
| ZMYND11 | Q15326 | 0.0071 | 0.48 | 1.39 |
| ZNF132 | P52740 | 7.90E-05 | 1.15 | 2.21 |
| ZNF177 | Q13360 | 5.80E-04 | 0.29 | 1.23 |
| ZNF185 | O15231 | 7.30E-04 | 0.36 | 1.28 |
| ZNF516 | Q92618 | 1.40E-04 | 0.31 | 1.24 |
| ZNF521 | Q96K83 | 0.0017 | 0.47 | 1.39 |
| ZNF548 | Q8NEK5 | 0.017 | 0.3 | 1.23 |
| ZNF778 | Q96MU6 | 0.002 | 0.45 | 1.37 |
| ZSCAN26 | Q16670 | 0.019 | 0.26 | 1.2 |

| **Supplementary Table 2B:** ***KDM6A*****KO downregulated protein expression by mass spec** [pVal = p-value; logFC = log of the fold change; FC = fold change in expression] | | | | |
|---|---|---|---|---|
| **Protein** | **Accession** | **pVal** | **logFC** | **FC** |
| ABCB1 | P08183 | 3.80E-04 | -0.62 | -1.53 |
| ABCC10 | Q5T3U5 | 0.0015 | -0.29 | -1.22 |
| ABCC4 | O15439 | 8.60E-05 | -0.32 | -1.25 |
| ABCG2 | Q9UNQ0 | 8.20E-04 | -0.56 | -1.47 |
| ABLIM1 | O14639 | 3.00E-04 | -0.46 | -1.37 |
| ACAA1 | P09110 | 1.20E-04 | -0.27 | -1.21 |
| ACAP1 | Q15027 | 0.0013 | -0.29 | -1.22 |
| ACSBG1 | Q96GR2 | 0.0025 | -0.6 | -1.51 |
| ACTG1 | P63261 | 0.0016 | -0.29 | -1.22 |
| ACVR1 | Q04771 | 3.90E-04 | -0.38 | -1.3 |
| ADAMTS7 | Q9UKP4 | 8.10E-04 | -0.35 | -1.28 |
| ADCY8 | P40145 | 1.70E-04 | -1.13 | -2.19 |
| ADCY9 | O60503 | 2.80E-04 | -0.51 | -1.42 |
| ADCYAP1R1 | P41586 | 0.0043 | -0.41 | -1.33 |
| ADGRL3 | Q9HAR2 | 8.00E-06 | -0.74 | -1.67 |
| AEN | Q8WTP8 | 0.0031 | -0.51 | -1.43 |
| AFAP1L2 | Q8N4X5 | 0.0024 | -0.99 | -1.99 |
| AGPAT2 | O15120 | 0.0024 | -0.28 | -1.21 |
| AGRN | O00468 | 0.0012 | -0.3 | -1.23 |
| AK5 | Q9Y6K8 | 1.90E-04 | -0.78 | -1.71 |
| AKAP7 | O43687 | 0.0018 | -0.88 | -1.84 |
| AKNA | Q7Z591 | 0.0075 | -0.4 | -1.32 |
| AKR1D1 | P51857 | 0.0069 | -0.37 | -1.29 |
| ALAS1 | P13196 | 1.70E-04 | -0.34 | -1.26 |
| ALOX15 | P16050 | 7.70E-05 | -1.03 | -2.04 |
| ALPL | P05186 | 8.50E-04 | -0.26 | -1.2 |
| AMD1 | P17707 | 0.0059 | -0.26 | -1.2 |
| AMIGO3 | Q86WK7 | 0.021 | -0.4 | -1.32 |
| ANKRD20A8P | Q5CZ79 | 3.90E-05 | -0.59 | -1.5 |
| ANXA1 | P04083 | 0.037 | -0.74 | -1.67 |
| ANXA2 | P07355 | 5.80E-04 | -0.65 | -1.57 |
| APC | P25054 | 0.0014 | -0.26 | -1.2 |
| AR | P10275 | 0.0083 | -0.34 | -1.26 |
| ARHGAP22 | Q7Z5H3 | 0.0023 | -0.36 | -1.28 |
| ARHGEF17 | Q96PE2 | 0.0015 | -0.27 | -1.21 |
| ARMCX4 | Q5H9R4 | 0.0012 | -0.27 | -1.2 |
| ARV1 | Q9H2C2 | 0.013 | -0.26 | -1.2 |
| ASAP3 | Q8TDY4 | 1.20E-04 | -0.29 | -1.22 |

| | | | | |
|---|---|---|---|---|
| ASPHD2 | Q6ICH7 | 0.0029 | -0.32 | -1.25 |
| ASTN2 | O75129 | 9.60E-06 | -0.61 | -1.53 |
| ATAD3B | Q5T9A4 | 3.10E-06 | -0.47 | -1.39 |
| ATAD5 | Q96QE3 | 0.026 | -0.49 | -1.41 |
| ATG14 | Q6ZNE5 | 0.0042 | -0.26 | -1.2 |
| ATG3 | Q9NT62 | 7.90E-05 | -0.33 | -1.26 |
| ATP1A3 | P13637 | 1.10E-06 | -0.51 | -1.42 |
| ATP1B1 | P05026 | 0.005 | -0.28 | -1.22 |
| ATP1B2 | P14415 | 2.00E-04 | -0.39 | -1.31 |
| ATP2A3 | Q93084 | 0.0038 | -0.31 | -1.24 |
| ATP5G1 | P05496 | 2.70E-05 | -1.54 | -2.91 |
| ATP5S | Q99766 | 0.0061 | -0.26 | -1.2 |
| ATPIF1 | Q9UII2 | 6.40E-04 | -0.35 | -1.27 |
| ATXN7L3 | Q14CW9 | 8.70E-06 | -1.05 | -2.06 |
| AURKAIP1 | Q9NWT8 | 6.30E-05 | -0.35 | -1.27 |
| BACE1 | P56817 | 0.0011 | -0.37 | -1.3 |
| BAG1 | Q99933 | 0.0028 | -0.31 | -1.24 |
| BAHCC1 | Q9P281 | 0.0037 | -0.58 | -1.5 |
| BAIAP2 | Q9UQB8 | 2.50E-05 | -0.47 | -1.38 |
| BCAM | P50895 | 0.001 | -0.35 | -1.28 |
| BCL11A | Q9H165 | 0.0024 | -0.35 | -1.27 |
| BCL2L12 | Q9HB09 | 3.20E-06 | -0.45 | -1.36 |
| BDP1 | A6H8Y1 | 0.012 | -0.3 | -1.23 |
| BHLHB9 | Q6PI77 | 0.0073 | -0.27 | -1.21 |
| BHMT | Q93088 | 5.50E-04 | -0.39 | -1.31 |
| BICC1 | Q9H694 | 5.60E-05 | -0.57 | -1.49 |
| BMS1 | Q14692 | 1.20E-04 | -0.3 | -1.23 |
| BOLA3 | Q53S33 | 1.50E-04 | -0.39 | -1.31 |
| BOP1 | Q14137 | 2.90E-05 | -0.32 | -1.25 |
| C12orf29 | Q8N999 | 0.0021 | -0.38 | -1.3 |
| C18orf21 | Q32NC0 | 0.0023 | -0.44 | -1.35 |
| C19orf43 | Q9BQ61 | 1.80E-04 | -0.3 | -1.23 |
| C19orf53 | Q9UNZ5 | 1.30E-04 | -0.65 | -1.57 |
| C1orf106 | Q3KP66 | 0.0051 | -0.27 | -1.2 |
| C1orf21 | Q9H246 | 7.50E-05 | -0.48 | -1.39 |
| C2CD3 | Q4AC94 | 0.0023 | -0.39 | -1.31 |
| C4A | P0C0L4 | 0.018 | -0.3 | -1.23 |
| C4orf3 | Q8WVX3 | 0.017 | -0.51 | -1.43 |
| C4orf32 | Q8N8J7 | 0.003 | -0.35 | -1.27 |
| C5orf46 | Q6UWT4 | 0.013 | -0.48 | -1.4 |
| C8orf59 | Q8N0T1 | 5.30E-05 | -0.38 | -1.3 |

| | | | | |
|---|---|---|---|---|
| C9orf16 | Q9BUW7 | 0.022 | -0.34 | -1.27 |
| C9orf85 | Q96MD7 | 9.60E-05 | -0.4 | -1.32 |
| CA3 | P07451 | 6.60E-06 | -0.45 | -1.37 |
| CA4 | P22748 | 7.60E-04 | -0.63 | -1.55 |
| CABP1 | Q9NZU7 | 0.018 | -0.45 | -1.37 |
| CABYR | O75952 | 0.027 | -0.32 | -1.25 |
| CADPS | Q9ULU8 | 2.70E-05 | -0.44 | -1.36 |
| CALB2 | P22676 | 4.30E-04 | -0.56 | -1.48 |
| CALD1 | Q05682 | 3.40E-04 | -0.51 | -1.43 |
| CALR | P27797 | 8.20E-04 | -0.28 | -1.21 |
| CAMKK1 | Q8N5S9 | 0.0031 | -0.3 | -1.23 |
| CAMKV | Q8NCB2 | 4.00E-04 | -0.3 | -1.23 |
| CAPN5 | O15484 | 2.70E-04 | -0.27 | -1.21 |
| CAPN6 | Q9Y6Q1 | 4.20E-04 | -0.49 | -1.4 |
| CARD10 | Q9BWT7 | 0.0089 | -0.27 | -1.21 |
| CARD11 | Q9BXL7 | 0.001 | -0.3 | -1.23 |
| CASC3 | O15234 | 5.00E-04 | -0.31 | -1.24 |
| CASKIN2 | Q8WXE0 | 0.0015 | -0.3 | -1.24 |
| CAV1 | Q03135 | 2.60E-04 | -0.83 | -1.78 |
| CBR3 | O75828 | 9.60E-05 | -0.7 | -1.63 |
| CCDC138 | Q96M89 | 0.0021 | -0.29 | -1.22 |
| CCDC14 | Q49A88 | 9.90E-05 | -0.29 | -1.22 |
| CCDC177 | Q9NQR7 | 7.80E-04 | -0.46 | -1.37 |
| CCDC59 | Q9P031 | 6.80E-04 | -0.45 | -1.37 |
| CCDC65 | Q8IXS2 | 3.30E-04 | -0.32 | -1.24 |
| CCDC86 | Q9H6F5 | 1.90E-04 | -0.32 | -1.25 |
| CD151 | P48509 | 0.034 | -0.28 | -1.22 |
| CD3EAP | O15446 | 1.50E-04 | -0.34 | -1.26 |
| CD44 | P16070 | 1.80E-04 | -0.69 | -1.62 |
| CD9 | P21926 | 6.10E-04 | -0.57 | -1.48 |
| CDC25A | P30304 | 0.0068 | -0.26 | -1.2 |
| CDC45 | O75419 | 0.004 | -0.27 | -1.2 |
| CDCA2 | Q69YH5 | 1.50E-04 | -0.31 | -1.24 |
| CDCA5 | Q96FF9 | 0.0013 | -0.32 | -1.25 |
| CDCA7 | Q9BWT1 | 0.0017 | -0.38 | -1.3 |
| CDCA7L | Q96GN5 | 4.70E-05 | -0.36 | -1.28 |
| CDCP1 | Q9H5V8 | 0.0016 | -0.33 | -1.26 |
| CDH11 | P55287 | 0.0037 | -0.53 | -1.45 |
| CDH13 | P55290 | 1.30E-04 | -0.96 | -1.94 |
| CDH2 | P19022 | 1.30E-06 | -0.88 | -1.84 |
| CDH3 | P22223 | 0.0019 | -0.26 | -1.2 |

| | | | | |
|---|---|---|---|---|
| CDIP1 | Q9H305 | 0.0041 | -0.3 | -1.24 |
| CDK14 | O94921 | 0.01 | -0.27 | -1.2 |
| CDKN2A | P42771 | 0.0025 | -0.6 | -1.52 |
| CDRT15 | Q96T59 | 7.60E-05 | -1.29 | -2.44 |
| CDS1 | Q92903 | 6.60E-05 | -0.36 | -1.29 |
| CEBPZ | Q03701 | 0.0012 | -0.28 | -1.21 |
| CELSR1 | Q9NYQ6 | 0.0032 | -0.4 | -1.32 |
| CEND1 | Q8N111 | 4.90E-05 | -0.5 | -1.42 |
| CENPS | Q8N2Z9 | 7.10E-04 | -0.32 | -1.25 |
| CEP131 | Q9UPN4 | 2.00E-05 | -0.35 | -1.27 |
| CERK | Q8TCT0 | 0.017 | -0.34 | -1.27 |
| CERS4 | Q9HA82 | 0.0036 | -0.31 | -1.24 |
| CHCHD10 | Q8WYQ3 | 0.0037 | -0.28 | -1.21 |
| CHCHD3 | Q9NX63 | 1.50E-04 | -0.29 | -1.22 |
| CHPF | Q8IZ52 | 0.0015 | -0.27 | -1.21 |
| CHST9 | Q7L1S5 | 0.015 | -0.38 | -1.3 |
| CLDN10 | P78369 | 0.041 | -0.3 | -1.23 |
| CLIP3 | Q96DZ5 | 0.0096 | -0.28 | -1.22 |
| CLN8 | Q9UBY8 | 1.00E-04 | -0.48 | -1.39 |
| CLSTN3 | Q9BQT9 | 5.50E-04 | -0.82 | -1.77 |
| CNKSR2 | Q8WXI2 | 1.30E-04 | -0.31 | -1.24 |
| CNN1 | P51911 | 0.0056 | -0.32 | -1.25 |
| CNPY4 | Q8N129 | 8.90E-04 | -0.36 | -1.29 |
| CNRIP1 | Q96F85 | 1.00E-04 | -0.57 | -1.48 |
| CNTN1 | Q12860 | 2.80E-05 | -0.47 | -1.38 |
| COBL | O75128 | 0.0011 | -0.3 | -1.23 |
| COL12A1 | Q99715 | 0.011 | -0.34 | -1.27 |
| COL14A1 | Q05707 | 3.90E-05 | -0.56 | -1.48 |
| COL1A1 | P02452 | 5.70E-05 | -0.69 | -1.61 |
| COL1A2 | P08123 | 0.0012 | -0.38 | -1.3 |
| COL5A2 | P05997 | 0.022 | -0.32 | -1.25 |
| COL6A2 | P12110 | 5.10E-05 | -0.39 | -1.31 |
| COL9A3 | Q14050 | 0.045 | -0.33 | -1.26 |
| COLEC12 | Q5KU26 | 0.0015 | -0.53 | -1.44 |
| COQ4 | Q9Y3A0 | 3.00E-05 | -0.67 | -1.59 |
| CORO1A | P31146 | 1.30E-06 | -0.76 | -1.69 |
| CORO2A | Q92828 | 0.0051 | -0.29 | -1.22 |
| COX16 | Q9P0S2 | 8.20E-05 | -0.41 | -1.32 |
| COX6C | P09669 | 0.0013 | -0.29 | -1.23 |
| COX7C | P15954 | 0.0096 | -0.26 | -1.2 |
| CPNE7 | Q9UBL6 | 8.00E-04 | -0.46 | -1.37 |

| | | | | |
|---|---|---|---|---|
| CPT1A | P50416 | 1.90E-04 | -0.75 | -1.69 |
| CR1L | Q2VPA4 | 0.014 | -0.43 | -1.35 |
| CRELD2 | Q6UXH1 | 3.60E-04 | -0.34 | -1.27 |
| CRIP1 | P50238 | 0.048 | -0.27 | -1.21 |
| CRYAB | P02511 | 4.10E-05 | -0.53 | -1.45 |
| CSDE1 | O75534 | 6.50E-05 | -0.34 | -1.26 |
| CSRP1 | P21291 | 2.40E-04 | -0.85 | -1.81 |
| CTGF | P29279 | 6.10E-06 | -0.83 | -1.78 |
| CTHRC1 | Q96CG8 | 0.0023 | -0.39 | -1.31 |
| CXCL11 | O14625 | 0.044 | -0.5 | -1.42 |
| CXorf57 | Q6NSI4 | 1.50E-05 | -0.41 | -1.33 |
| CYR61 | O00622 | 1.30E-04 | -0.7 | -1.62 |
| DAAM1 | Q9Y4D1 | 9.00E-05 | -0.41 | -1.33 |
| DAB1 | O75553 | 5.40E-08 | -1.03 | -2.04 |
| DAG1 | Q14118 | 0.0059 | -0.27 | -1.21 |
| DDX18 | Q9NVP1 | 4.30E-04 | -0.37 | -1.3 |
| DDX21 | Q9NR30 | 0.023 | -0.37 | -1.29 |
| DDX24 | Q9GZR7 | 3.20E-04 | -0.53 | -1.44 |
| DDX27 | Q96GQ7 | 5.60E-04 | -0.28 | -1.22 |
| DDX52 | Q9Y2R4 | 0.003 | -0.51 | -1.42 |
| DEFA4 | P12838 | 0.0058 | -0.46 | -1.38 |
| DES | P17661 | 0.0015 | -0.42 | -1.33 |
| DGUOK | Q16854 | 5.20E-04 | -0.5 | -1.41 |
| DHRS11 | Q6UWP2 | 0.0072 | -0.26 | -1.2 |
| DIDO1 | Q9BTC0 | 2.20E-04 | -0.31 | -1.24 |
| DKC1 | O60832 | 7.60E-04 | -0.28 | -1.21 |
| DLG2 | Q15700 | 0.0015 | -0.56 | -1.47 |
| DLG5 | Q8TDM6 | 1.20E-04 | -0.26 | -1.2 |
| DMKN | Q6E0U4 | 0.0055 | -0.4 | -1.32 |
| DNAH8 | Q96JB1 | 0.018 | -1.16 | -2.24 |
| DNAJA4 | Q8WW22 | 4.40E-06 | -0.47 | -1.39 |
| DNAJC14 | Q6Y2X3 | 0.0056 | -0.33 | -1.25 |
| DNAJC15 | Q9Y5T4 | 5.80E-05 | -0.52 | -1.43 |
| DNLZ | Q5SXM8 | 1.40E-05 | -0.56 | -1.47 |
| DNMT3B | Q9UBC3 | 0.017 | -0.6 | -1.52 |
| DPH3 | Q96FX2 | 2.30E-07 | -0.93 | -1.91 |
| DPPA2 | Q7Z7J5 | 2.30E-04 | -0.68 | -1.6 |
| DPPA4 | Q7L190 | 4.40E-05 | -0.52 | -1.43 |
| DTNA | Q9Y4J8 | 5.80E-05 | -0.56 | -1.47 |
| DUOXA2 | Q1HG44 | 0.001 | -0.96 | -1.94 |
| DUSP11 | O75319 | 0.0039 | -0.27 | -1.21 |

| | | | | |
|---|---|---|---|---|
| DUSP16 | Q9BY84 | 9.70E-04 | -0.27 | -1.2 |
| DVL1 | O14640 | 0.0016 | -0.26 | -1.2 |
| DYSF | O75923 | 1.70E-04 | -0.29 | -1.22 |
| ECHDC1 | Q9NTX5 | 2.50E-05 | -0.33 | -1.25 |
| EHD1 | Q9H4M9 | 1.40E-05 | -0.61 | -1.53 |
| EHD2 | Q9NZN4 | 7.00E-05 | -0.4 | -1.32 |
| EHD4 | Q9H223 | 1.40E-05 | -0.49 | -1.41 |
| EHHADH | Q08426 | 0.017 | -0.34 | -1.27 |
| ELAVL2 | Q12926 | 1.10E-04 | -0.36 | -1.28 |
| ELOA | Q14241 | 0.026 | -0.42 | -1.34 |
| ELOF1 | P60002 | 0.012 | -0.28 | -1.22 |
| EMC6 | Q9BV81 | 9.80E-04 | -0.41 | -1.33 |
| ENAH | Q8N8S7 | 3.10E-04 | -0.26 | -1.2 |
| ENPP1 | P22413 | 0.0069 | -0.43 | -1.34 |
| EPB41L1 | Q9H4G0 | 6.60E-08 | -0.86 | -1.81 |
| EPHB6 | O15197 | 1.30E-04 | -0.55 | -1.46 |
| EPPK1 | P58107 | 2.30E-04 | -0.32 | -1.24 |
| ERICH5 | Q6P6B1 | 0.014 | -0.38 | -1.3 |
| ERRFI1 | Q9UJM3 | 1.80E-04 | -0.52 | -1.44 |
| ERVH48-1 | M5A8F1 | 1.60E-05 | -0.95 | -1.94 |
| ERVK-6 | Q7LDI9 | 3.70E-04 | -0.59 | -1.51 |
| ERVMER34-1 | Q9H9K5 | 5.10E-05 | -0.34 | -1.27 |
| ESYT1 | Q9BSJ8 | 5.10E-04 | -0.26 | -1.2 |
| EVPL | Q92817 | 3.40E-06 | -1.47 | -2.77 |
| EXOSC6 | Q5RKV6 | 4.40E-04 | -0.28 | -1.22 |
| EXOSC8 | Q96B26 | 3.30E-04 | -0.29 | -1.22 |
| F101B_HUMAN | NA | 4.50E-06 | -1.01 | -2.02 |
| F11R | Q9Y624 | 3.40E-04 | -0.33 | -1.26 |
| F212A_HUMAN | NA | 1.70E-04 | -0.34 | -1.27 |
| FA2H | Q7L5A8 | 3.30E-04 | -0.62 | -1.54 |
| FABP5 | Q01469 | 0.02 | -0.26 | -1.2 |
| FAM126A | Q9BYI3 | 3.10E-04 | -0.26 | -1.2 |
| FAM171A2 | A8MVW0 | 0.0052 | -0.31 | -1.24 |
| FAM174A | Q8TBP5 | 0.021 | -0.39 | -1.31 |
| FAM207A | Q9NSI2 | 1.20E-04 | -0.26 | -1.2 |
| FAM20A | Q96MK3 | 0.017 | -0.76 | -1.69 |
| FAM210B | Q96KR6 | 0.0043 | -0.45 | -1.36 |
| FAM49A | Q9H0Q0 | 7.40E-05 | -0.27 | -1.21 |
| FAM69B | Q5VUD6 | 1.90E-04 | -0.57 | -1.48 |
| FAR1 | Q8WVX9 | 2.70E-04 | -0.28 | -1.21 |
| FARP1 | Q9Y4F1 | 0.005 | -0.27 | -1.21 |

| | | | | |
|---|---|---|---|---|
| FAU | P35544 | 2.10E-04 | -0.63 | -1.55 |
| FBXO9 | Q9UK97 | 0.001 | -0.29 | -1.22 |
| FERMT1 | Q9BQL6 | 2.30E-04 | -0.34 | -1.27 |
| FGF2 | P09038 | 0.0021 | -0.74 | -1.67 |
| FHL1 | Q13642 | 0.0016 | -0.48 | -1.4 |
| FHL2 | Q14192 | 9.90E-05 | -0.4 | -1.32 |
| FKBP11 | Q9NYL4 | 1.40E-04 | -0.32 | -1.24 |
| FKBP14 | Q9NWM8 | 0.0017 | -0.29 | -1.23 |
| FKRP | Q9H9S5 | 7.40E-05 | -0.37 | -1.29 |
| FLNB | O75369 | 2.80E-04 | -0.29 | -1.22 |
| FLNC | Q14315 | 0.0056 | -0.52 | -1.43 |
| FN1 | P02751 | 0.0011 | -0.58 | -1.5 |
| FOXH1 | O75593 | 6.70E-05 | -1.2 | -2.3 |
| FOXK1 | P85037 | 2.00E-05 | -0.47 | -1.38 |
| FOXM1 | Q08050 | 3.00E-05 | -0.64 | -1.56 |
| FOXO4 | P98177 | 5.30E-07 | -1.09 | -2.12 |
| FOXRED2 | Q8IWF2 | 6.80E-04 | -0.42 | -1.33 |
| FTSJ3 | Q8IY81 | 9.80E-05 | -0.35 | -1.27 |
| FUNDC1 | Q8IVP5 | 3.00E-05 | -0.44 | -1.36 |
| FUT4 | P22083 | 0.0082 | -0.31 | -1.24 |
| FZD5 | Q13467 | 0.0022 | -0.39 | -1.31 |
| FZD7 | O75084 | 3.40E-04 | -0.34 | -1.27 |
| GABRA3 | P34903 | 3.90E-05 | -0.46 | -1.37 |
| GALNT14 | Q96FL9 | 0.0051 | -0.48 | -1.4 |
| GAP43 | P17677 | 1.60E-04 | -0.9 | -1.86 |
| GAR1 | Q9NY12 | 3.10E-04 | -0.31 | -1.24 |
| GAS2L3 | Q86XJ1 | 6.20E-04 | -0.67 | -1.59 |
| GDF1 | P27539 | 0.001 | -0.73 | -1.66 |
| GDF3 | Q9NR23 | 9.90E-06 | -0.82 | -1.76 |
| GGCT | O75223 | 0.0015 | -0.26 | -1.2 |
| GJC2 | Q5T442 | 0.021 | -0.45 | -1.37 |
| GLB1L2 | Q8IW92 | 0.0015 | -0.26 | -1.2 |
| GLB1L3 | Q8NCI6 | 0.0053 | -0.37 | -1.29 |
| GLMP | Q8WWB7 | 0.011 | -0.45 | -1.37 |
| GNA14 | O95837 | 8.30E-04 | -0.37 | -1.29 |
| GNAS | Q5JWF2 | 0.0017 | -0.35 | -1.28 |
| GNG2 | P59768 | 1.00E-04 | -0.26 | -1.2 |
| GNG3 | P63215 | 0.0057 | -0.46 | -1.38 |
| GNL2 | Q13823 | 7.30E-04 | -0.36 | -1.28 |
| GNL3 | Q9BVP2 | 3.20E-04 | -0.28 | -1.22 |
| GOT2 | P00505 | 8.60E-05 | -0.8 | -1.74 |

| | | | | |
|---|---|---|---|---|
| GPATCH4 | Q5T3I0 | 2.00E-04 | -0.36 | -1.29 |
| GPC3 | P51654 | 0.0014 | -0.42 | -1.34 |
| GPC4 | O75487 | 4.30E-05 | -0.65 | -1.57 |
| GPC6 | Q9Y625 | 0.0021 | -0.34 | -1.27 |
| GPRIN1 | Q7Z2K8 | 3.70E-05 | -0.39 | -1.31 |
| GPX1 | P07203 | 6.80E-04 | -0.29 | -1.22 |
| GSCR2_HUMAN | NA | 0.0034 | -0.28 | -1.21 |
| GSG2 | Q8TF76 | 0.025 | -0.28 | -1.21 |
| GSTM4 | Q03013 | 0.011 | -0.35 | -1.28 |
| GTF3A | Q92664 | 2.20E-04 | -0.27 | -1.21 |
| GTPBP4 | Q9BZE4 | 6.10E-04 | -0.42 | -1.34 |
| GTSF1 | Q8WW33 | 4.90E-05 | -0.58 | -1.49 |
| GUK1 | Q16774 | 0.0095 | -0.26 | -1.2 |
| H2AFX | P16104 | 5.80E-04 | -0.33 | -1.25 |
| HACE1 | Q8IYU2 | 0.027 | -0.28 | -1.21 |
| HAO1 | Q9UJM8 | 1.80E-05 | -0.35 | -1.28 |
| HAS3 | O00219 | 0.0026 | -0.4 | -1.32 |
| HEG1 | Q9ULI3 | 0.0096 | -0.27 | -1.21 |
| HIST1H4A | P62805 | 0.045 | -0.71 | -1.64 |
| HIST3H2BB | Q8N257 | 5.00E-05 | -0.5 | -1.42 |
| HLA-C | Q07000 | 0.02 | -0.27 | -1.21 |
| HMGCR | P04035 | 0.0031 | -0.44 | -1.35 |
| HMGN4 | O00479 | 3.70E-04 | -1.02 | -2.03 |
| HMGN5 | P82970 | 1.30E-04 | -0.39 | -1.31 |
| HN1_HUMAN | NA | 0.0019 | -0.26 | -1.2 |
| HNMT | P50135 | 7.70E-04 | -0.35 | -1.28 |
| HNRNPCL1 | O60812 | 4.70E-05 | -0.43 | -1.34 |
| HPDL | Q96IR7 | 5.90E-06 | -0.59 | -1.5 |
| HPSE | Q9Y251 | 1.90E-04 | -0.41 | -1.33 |
| HRSL3_HUMAN | NA | 0.014 | -0.36 | -1.28 |
| HSPA1L | P34931 | 1.50E-05 | -0.39 | -1.31 |
| HSPA2 | P54652 | 0.0021 | -0.33 | -1.26 |
| HSPA5 | P11021 | 3.60E-06 | -0.4 | -1.32 |
| HSPB1 | P04792 | 4.70E-06 | -0.5 | -1.42 |
| HTR1A | P08908 | 3.70E-06 | -1.86 | -3.64 |
| HYOU1 | Q9Y4L1 | 2.00E-05 | -0.39 | -1.31 |
| IGF1R | P08069 | 0.0013 | -0.28 | -1.21 |
| IGSF8 | Q969P0 | 4.20E-04 | -0.39 | -1.31 |
| IL17RD | Q8NFM7 | 7.10E-05 | -0.45 | -1.36 |
| IMUP | Q9GZP8 | 6.10E-04 | -0.53 | -1.45 |
| INA | Q16352 | 2.60E-05 | -0.58 | -1.49 |

| | | | | |
|---|---|---|---|---|
| IQCE | Q6IPM2 | 0.0058 | -0.86 | -1.81 |
| IQGAP1 | P46940 | 5.40E-04 | -0.27 | -1.21 |
| IQSEC2 | Q5JU85 | 5.80E-06 | -0.67 | -1.59 |
| IRF4 | Q15306 | 0.0098 | -0.27 | -1.2 |
| ISG20 | Q96AZ6 | 1.60E-04 | -0.59 | -1.5 |
| ISG20L2 | Q9H9L3 | 3.20E-04 | -0.38 | -1.3 |
| ITGA1 | P56199 | 4.30E-05 | -0.4 | -1.32 |
| ITGA3 | P26006 | 0.0017 | -0.3 | -1.23 |
| ITGA5 | P08648 | 1.80E-04 | -0.31 | -1.24 |
| ITGA9 | Q13797 | 0.0017 | -0.58 | -1.49 |
| ITGAV | P06756 | 0.0023 | -0.27 | -1.2 |
| ITGB1 | P05556 | 1.40E-05 | -0.63 | -1.55 |
| ITGB4 | P16144 | 0.04 | -0.3 | -1.23 |
| ITGB5 | P18084 | 1.50E-05 | -0.52 | -1.43 |
| ITM2A | O43736 | 1.20E-05 | -0.68 | -1.61 |
| JAM2 | P57087 | 3.20E-05 | -0.49 | -1.41 |
| JARID2 | Q92833 | 0.0055 | -0.76 | -1.69 |
| JAZF1 | Q86VZ6 | 3.20E-04 | -0.59 | -1.51 |
| JPH1 | Q9HDC5 | 0.0015 | -0.32 | -1.25 |
| JSRP1 | Q96MG2 | 7.50E-05 | -1.41 | -2.65 |
| KCNK5 | O95279 | 2.80E-04 | -0.29 | -1.22 |
| KDM2B | Q8NHM5 | 6.10E-04 | -0.35 | -1.27 |
| KDM6A | O15550 | 3.50E-07 | -1.27 | -2.41 |
| KIAA0100 | Q14667 | 0.017 | -0.28 | -1.22 |
| KIAA0754 | O94854 | 9.70E-05 | -0.33 | -1.26 |
| KIAA1522 | Q9P206 | 0.0024 | -0.29 | -1.22 |
| KIAA2013 | Q8IYS2 | 0.0083 | -0.27 | -1.2 |
| KIF27 | Q86VH2 | 1.50E-05 | -0.54 | -1.45 |
| KIT | P10721 | 4.80E-04 | -0.47 | -1.38 |
| KLF4 | O43474 | 0.00093 | -0.91 | -1.88 |
| KLHL21 | Q9UJP4 | 0.0023 | -0.33 | -1.25 |
| KMT2C | Q8NEZ4 | 8.40E-04 | -0.39 | -1.31 |
| KRI1 | Q8N9T8 | 3.10E-05 | -0.38 | -1.3 |
| KRR1 | Q13601 | 0.0011 | -0.36 | -1.29 |
| L1CAM | P32004 | 0.0062 | -0.3 | -1.23 |
| L1RE1 | Q9UN81 | 2.10E-05 | -0.73 | -1.66 |
| L1TD1 | Q5T7N2 | 1.40E-04 | -0.26 | -1.2 |
| LACTB | P83111 | 0.0024 | -0.28 | -1.22 |
| LAD1 | O00515 | 5.50E-04 | -0.32 | -1.25 |
| LAMA2 | P24043 | 0.003 | -0.32 | -1.24 |
| LAMC2 | Q13753 | 1.20E-05 | -0.97 | -1.96 |

| | | | | |
|---|---|---|---|---|
| LAMC3 | Q9Y6N6 | 6.80E-05 | -0.57 | -1.49 |
| LARP1B | Q659C4 | 0.0042 | -0.28 | -1.21 |
| LCP1 | P13796 | 0.0091 | -0.38 | -1.3 |
| LDLR | P01130 | 0.012 | -0.31 | -1.24 |
| LECT1_HUMAN | NA | 1.20E-04 | -0.34 | -1.27 |
| LEFTY1 | O75610 | 0.0019 | -0.55 | -1.47 |
| LEMD1 | Q68G75 | 0.0082 | -0.44 | -1.36 |
| LIMS2 | Q7Z4I7 | 0.001 | -0.54 | -1.46 |
| LIN28A | Q9H9Z2 | 3.00E-04 | -0.29 | -1.22 |
| LIPE | Q05469 | 7.80E-04 | -0.3 | -1.23 |
| LITAF | Q99732 | 1.10E-07 | -1.39 | -2.62 |
| LMBR1 | Q8WVP7 | 0.017 | -0.3 | -1.23 |
| LMCD1 | Q9NZU5 | 4.00E-07 | -0.65 | -1.57 |
| LMNA | P02545 | 0.0026 | -0.26 | -1.2 |
| LPL | P06858 | 4.20E-05 | -0.59 | -1.51 |
| LRP2 | P98164 | 0.0027 | -0.45 | -1.37 |
| LRPAP1 | P30533 | 5.40E-05 | -0.34 | -1.27 |
| LRRC59 | Q96AG4 | 4.20E-05 | -0.48 | -1.4 |
| LRRC8C | Q8TDW0 | 0.0045 | -0.6 | -1.51 |
| LRRC8D | Q7L1W4 | 0.0018 | -0.27 | -1.2 |
| LSAMP | Q13449 | 0.015 | -0.28 | -1.21 |
| LUZP1 | Q86V48 | 9.90E-05 | -0.27 | -1.2 |
| LYAR | Q9NX58 | 0.0024 | -0.37 | -1.3 |
| LYPD3 | O95274 | 1.20E-06 | -0.94 | -1.91 |
| MAGEL2 | Q9UJ55 | 0.003 | -0.38 | -1.3 |
| MAP1A | P78559 | 9.40E-05 | -0.39 | -1.31 |
| MAPK13 | O15264 | 0.0015 | -0.34 | -1.27 |
| MAST4 | O15021 | 4.70E-04 | -0.7 | -1.62 |
| MATK | P42679 | 9.10E-05 | -0.36 | -1.29 |
| MB21D1 | Q8N884 | 0.025 | -0.5 | -1.42 |
| MBD2 | Q9UBB5 | 0.029 | -0.42 | -1.34 |
| MBD4 | O95243 | 0.0018 | -0.32 | -1.24 |
| MBD6 | Q96DN6 | 0.03 | -0.35 | -1.27 |
| MCAM | P43121 | 1.60E-06 | -1.08 | -2.11 |
| MCM10 | Q7L590 | 0.0074 | -0.3 | -1.23 |
| MCUB | Q9NWR8 | 1.40E-04 | -0.35 | -1.27 |
| MDM2 | Q00987 | 0.0016 | -0.4 | -1.32 |
| MDN1 | Q9NU22 | 4.60E-04 | -0.29 | -1.22 |
| MET | P08581 | 0.0019 | -0.4 | -1.32 |
| METTL7A | Q9H8H3 | 7.10E-08 | -1.06 | -2.09 |
| MFAP3L | O75121 | 0.022 | -0.38 | -1.3 |

| | | | | |
|---|---|---|---|---|
| MGST2 | Q99735 | 0.0013 | -0.34 | -1.27 |
| MICA | Q29983 | 0.039 | -0.28 | -1.21 |
| MIDN | Q504T8 | 8.10E-04 | -0.33 | -1.26 |
| MKNK1 | Q9BUB5 | 0.018 | -0.29 | -1.22 |
| MLEC | Q14165 | 0.0022 | -0.26 | -1.2 |
| MME | P08473 | 0.0016 | -0.53 | -1.44 |
| MMP15 | P51511 | 0.002 | -0.27 | -1.2 |
| MORF4L1 | Q9UBU8 | 0.0098 | -0.33 | -1.26 |
| MORF4L2 | Q15014 | 0.014 | -0.3 | -1.23 |
| MPC2 | O95563 | 9.90E-04 | -0.27 | -1.2 |
| MPP1 | Q00013 | 0.013 | -0.32 | -1.24 |
| MRPL11 | Q9Y3B7 | 2.70E-04 | -0.26 | -1.2 |
| MRPL12 | P52815 | 0.0024 | -0.28 | -1.22 |
| MRPL32 | Q9BYC8 | 0.005 | -0.42 | -1.33 |
| MRPL46 | Q9H2W6 | 2.20E-04 | -0.28 | -1.22 |
| MRPL54 | Q6P161 | 4.40E-05 | -0.36 | -1.28 |
| MRPL9 | Q9BYD2 | 4.20E-05 | -0.29 | -1.22 |
| MRPS16 | Q9Y3D3 | 6.60E-04 | -0.26 | -1.2 |
| MRPS17 | Q9Y2R5 | 0.0065 | -0.3 | -1.23 |
| MRPS36 | P82909 | 1.40E-04 | -0.31 | -1.24 |
| MRRF | Q96E11 | 1.00E-05 | -0.35 | -1.28 |
| MSRA | Q9UJ68 | 7.80E-04 | -0.29 | -1.22 |
| MT1B | P07438 | 0.039 | -0.92 | -1.89 |
| MT1F | P04733 | 8.70E-04 | -0.92 | -1.89 |
| MT1G | P13640 | 0.015 | -0.42 | -1.34 |
| MT1H | P80294 | 0.013 | -1.28 | -2.43 |
| MT1X | P80297 | 1.90E-04 | -1.24 | -2.35 |
| MTCH1 | Q9NZJ7 | 0.0027 | -0.3 | -1.23 |
| MTIF2 | P46199 | 0.0018 | -0.28 | -1.22 |
| MXRA8 | Q9BRK3 | 0.023 | -0.36 | -1.28 |
| MYH1 | P12882 | 1.00E-05 | -1.33 | -2.51 |
| MYH10 | P35580 | 5.40E-05 | -0.28 | -1.22 |
| MYH11 | P35749 | 0.0081 | -0.29 | -1.23 |
| MYL12A | P19105 | 7.50E-04 | -0.5 | -1.42 |
| MYOF | Q9NZM1 | 0.0023 | -0.46 | -1.38 |
| MYSM1 | Q5VVJ2 | 8.00E-04 | -0.3 | -1.23 |
| MZT2B | Q6NZ67 | 0.0027 | -0.55 | -1.46 |
| NACAD | O15069 | 6.40E-04 | -0.35 | -1.27 |
| NANOG | Q9H9S0 | 0.035 | -0.49 | -1.41 |
| NBEAL1 | Q6ZS30 | 0.018 | -0.35 | -1.27 |
| NCOA4 | Q13772 | 9.00E-04 | -0.28 | -1.21 |

| | | | | |
|---|---|---|---|---|
| NCR3LG1 | Q68D85 | 0.01 | -0.28 | -1.21 |
| NDUFA2 | O43678 | 0.0061 | -0.26 | -1.2 |
| NDUFA3 | O95167 | 2.60E-04 | -0.35 | -1.27 |
| NDUFB9 | Q9Y6M9 | 0.0018 | -0.27 | -1.21 |
| NDUFS4 | O43181 | 0.0067 | -0.27 | -1.2 |
| NDUFS6 | O75380 | 1.30E-05 | -0.41 | -1.33 |
| NDUFV1 | P49821 | 0.0033 | -0.27 | -1.2 |
| NDUFV2 | P19404 | 0.0019 | -0.26 | -1.2 |
| NEFH | P12036 | 7.30E-04 | -0.94 | -1.92 |
| NEFL | P07196 | 4.50E-06 | -0.97 | -1.95 |
| NEFM | P07197 | 5.30E-08 | -0.77 | -1.7 |
| NEPRO | Q6NW34 | 0.0015 | -0.35 | -1.27 |
| NES | P48681 | 4.50E-06 | -0.72 | -1.65 |
| NFE2L3 | Q9Y4A8 | 3.90E-04 | -0.53 | -1.44 |
| NFU1 | Q9UMS0 | 0.002 | -0.29 | -1.22 |
| NFXL1 | Q6ZNB6 | 5.00E-04 | -0.54 | -1.45 |
| NGFR | P08138 | 0.019 | -0.44 | -1.35 |
| NIFK | Q9BYG3 | 8.40E-04 | -0.38 | -1.31 |
| NLGN3 | Q9NZ94 | 4.40E-05 | -0.61 | -1.53 |
| NMRK2 | Q9NPI5 | 3.30E-04 | -0.71 | -1.63 |
| NOC3L | Q8WTT2 | 0.0063 | -0.37 | -1.3 |
| NOL7 | Q9UMY1 | 6.20E-05 | -0.31 | -1.24 |
| NOP16 | Q9Y3C1 | 3.30E-05 | -0.32 | -1.25 |
| NOP2 | P46087 | 0.0024 | -0.41 | -1.33 |
| NOP56 | O00567 | 0.035 | -0.32 | -1.25 |
| NOTCH2 | Q04721 | 0.0027 | -0.27 | -1.21 |
| NOTCH3 | Q9UM47 | 6.70E-05 | -0.36 | -1.29 |
| NOVA2 | Q9UNW9 | 7.00E-04 | -0.29 | -1.22 |
| NPM1 | P06748 | 2.70E-04 | -0.26 | -1.2 |
| NPTX2 | P47972 | 7.80E-05 | -0.74 | -1.67 |
| NPTXR | O95502 | 3.40E-04 | -0.37 | -1.29 |
| NRBF2 | Q96F24 | 4.50E-04 | -0.26 | -1.2 |
| NRK | Q7Z2Y5 | 4.10E-06 | -1.15 | -2.22 |
| NRXN2 | Q9P2S2 | 8.20E-04 | -0.45 | -1.37 |
| NSA2 | O95478 | 4.50E-05 | -0.46 | -1.37 |
| NSD1 | Q96L73 | 1.40E-04 | -0.31 | -1.24 |
| NSG1 | P42857 | 0.0041 | -0.38 | -1.3 |
| NT5C3A | Q9H0P0 | 1.70E-04 | -0.31 | -1.24 |
| NUDT8 | Q8WV74 | 0.0063 | -0.3 | -1.23 |
| NUS1 | Q96E22 | 3.00E-04 | -0.32 | -1.25 |
| OGFOD2 | Q6N063 | 0.0013 | -0.37 | -1.3 |

| | | | | |
|---|---|---|---|---|
| OLFM2 | O95897 | 0.0028 | -0.33 | -1.25 |
| ORMDL2 | Q53FV1 | 1.70E-05 | -0.46 | -1.38 |
| OSBPL10 | Q9BXB5 | 2.40E-04 | -0.26 | -1.2 |
| P3H1 | Q32P28 | 7.20E-05 | -0.29 | -1.22 |
| P4HB | P07237 | 3.10E-04 | -0.27 | -1.2 |
| PACSIN1 | Q9BY11 | 2.80E-04 | -0.33 | -1.25 |
| PALM2 | Q8IXS6 | 3.30E-05 | -0.43 | -1.34 |
| PALM3 | A6NDB9 | 4.00E-04 | -0.31 | -1.24 |
| PARP8 | Q8N3A8 | 0.0019 | -0.31 | -1.24 |
| PCDHB2 | Q9Y5E7 | 5.90E-04 | -0.39 | -1.31 |
| PCDHGB4 | Q9UN71 | 9.10E-05 | -0.46 | -1.38 |
| PCDHGC3 | Q9UN70 | 0.003 | -0.35 | -1.27 |
| PCGF2 | P35227 | 2.20E-05 | -0.39 | -1.31 |
| PCOLCE | Q15113 | 1.30E-04 | -0.47 | -1.39 |
| PCP4L1 | A6NKN8 | 2.60E-04 | -0.35 | -1.28 |
| PCSK5 | Q92824 | 3.20E-05 | -0.88 | -1.85 |
| PDCD2 | Q16342 | 5.10E-06 | -0.5 | -1.42 |
| PDE12 | Q6L8Q7 | 6.70E-05 | -0.42 | -1.34 |
| PDLIM3 | Q53GG5 | 1.40E-05 | -0.51 | -1.42 |
| PDLIM7 | Q9NR12 | 5.40E-04 | -0.44 | -1.36 |
| PDZD4 | Q76G19 | 3.90E-06 | -0.76 | -1.7 |
| PES1 | O00541 | 8.00E-04 | -0.35 | -1.28 |
| PET117 | Q6UWS5 | 0.031 | -0.45 | -1.37 |
| PINX1 | Q96BK5 | 2.20E-05 | -0.46 | -1.38 |
| PKDREJ | Q9NTG1 | 2.00E-04 | -0.27 | -1.2 |
| PKN1 | Q16512 | 3.20E-05 | -0.3 | -1.23 |
| PKP2 | Q99959 | 0.0023 | -0.38 | -1.3 |
| PKP3 | Q9Y446 | 1.40E-04 | -0.35 | -1.28 |
| PLA2G3 | Q9NZ20 | 4.60E-05 | -0.39 | -1.31 |
| PLA2G4A | P47712 | 0.024 | -0.39 | -1.31 |
| PLEKHF2 | Q9H8W4 | 1.40E-04 | -0.29 | -1.22 |
| PLIN4 | Q96Q06 | 0.04 | -0.29 | -1.22 |
| PLXNB1 | O43157 | 0.0027 | -0.28 | -1.21 |
| PNMA3 | Q9UL41 | 3.40E-04 | -0.43 | -1.34 |
| PNMA6A | P0CW24 | 8.50E-04 | -0.35 | -1.28 |
| PNRC2 | Q9NPJ4 | 0.0066 | -0.38 | -1.3 |
| POLR1E | Q9GZS1 | 0.004 | -0.33 | -1.26 |
| POM121 | Q96HA1 | 0.0018 | -0.31 | -1.24 |
| POP1 | Q99575 | 5.90E-06 | -0.41 | -1.33 |
| POTEF | A5A3E0 | 0.012 | -1.13 | -2.19 |
| POU2F2 | P09086 | 6.80E-04 | -0.44 | -1.35 |

| | | | | |
|---|---|---|---|---|
| PPAN | Q9NQ55 | 4.50E-04 | -0.46 | -1.37 |
| PPIB | P23284 | 7.10E-05 | -0.32 | -1.25 |
| PPP1R14A | Q96A00 | 3.70E-06 | -0.65 | -1.57 |
| PPP1R17 | O96001 | 0.0085 | -0.42 | -1.34 |
| PPP1R3D | O95685 | 0.0065 | -0.31 | -1.24 |
| PPTC7 | Q8NI37 | 0.016 | -0.29 | -1.22 |
| PRAF2 | O60831 | 5.90E-04 | -0.43 | -1.35 |
| PRDBP_HUMAN | NA | 3.50E-04 | -0.65 | -1.57 |
| PRDM14 | Q9GZV8 | 8.00E-04 | -0.28 | -1.22 |
| PRDM5 | Q9NQX1 | 0.022 | -0.39 | -1.31 |
| PRKG1 | Q13976 | 2.40E-04 | -0.71 | -1.64 |
| PROCR | Q9UNN8 | 0.0038 | -0.41 | -1.33 |
| PRR11 | Q96HE9 | 5.30E-05 | -0.44 | -1.36 |
| PRRG3 | Q9BZD7 | 0.012 | -0.58 | -1.5 |
| PTRF_HUMAN | NA | 2.60E-04 | -0.79 | -1.73 |
| PTRHD1 | Q6GMV3 | 2.90E-08 | -0.92 | -1.9 |
| PVR | P15151 | 0.0073 | -0.31 | -1.24 |
| PWWP2A | Q96N64 | 2.00E-04 | -0.35 | -1.28 |
| PXN | P49023 | 0.0014 | -0.27 | -1.21 |
| PYURF | Q96I23 | 8.60E-05 | -0.47 | -1.38 |
| R3HDM4 | Q96D70 | 2.40E-04 | -0.56 | -1.47 |
| RAB15 | P59190 | 0.0065 | -0.39 | -1.31 |
| RAB20 | Q9NX57 | 0.0013 | -0.32 | -1.24 |
| RAP1GAP2 | Q684P5 | 4.90E-05 | -0.38 | -1.3 |
| RAPH1 | Q70E73 | 1.10E-04 | -0.36 | -1.29 |
| RASL11B | Q9BPW5 | 0.036 | -0.37 | -1.29 |
| RASSF8 | Q8NHQ8 | 0.0022 | -0.32 | -1.25 |
| RBM28 | Q9NW13 | 6.90E-04 | -0.68 | -1.6 |
| RBM34 | P42696 | 2.50E-04 | -0.56 | -1.47 |
| RCN3 | Q96D15 | 2.80E-04 | -0.52 | -1.43 |
| RFTN1 | Q14699 | 3.70E-04 | -0.63 | -1.54 |
| RGS10 | O43665 | 3.90E-04 | -0.43 | -1.35 |
| RHOC | P08134 | 3.70E-04 | -0.8 | -1.74 |
| ROR2 | Q01974 | 4.40E-05 | -0.76 | -1.69 |
| RPF2 | Q9H7B2 | 0.0022 | -0.58 | -1.49 |
| RPIA | P49247 | 2.80E-05 | -0.33 | -1.25 |
| RPL10A | P62906 | 5.10E-04 | -0.51 | -1.42 |
| RPL17 | P18621 | 2.50E-04 | -0.42 | -1.34 |
| RPL23 | P62829 | 5.20E-05 | -0.47 | -1.38 |
| RPL23A | P62750 | 3.30E-04 | -0.29 | -1.22 |
| RPL27A | P46776 | 2.80E-04 | -0.33 | -1.26 |

| | | | | |
|---|---|---|---|---|
| RPL29 | P47914 | 0.001 | -0.43 | -1.35 |
| RPL34 | P49207 | 0.024 | -0.36 | -1.28 |
| RPL36 | Q9Y3U8 | 0.0012 | -0.28 | -1.21 |
| RPL36AL | Q969Q0 | 0.0057 | -0.35 | -1.27 |
| RPL4 | P36578 | 0.0013 | -0.27 | -1.21 |
| RPL7A | P62424 | 0.0044 | -0.48 | -1.39 |
| RPL7L1 | Q6DKI1 | 0.046 | -0.26 | -1.2 |
| RPL8 | P62917 | 0.0012 | -0.36 | -1.28 |
| RPLP1 | P05386 | 0.0041 | -0.32 | -1.25 |
| RPS18 | P62269 | 4.40E-04 | -0.26 | -1.2 |
| RPS21 | P63220 | 9.20E-04 | -0.33 | -1.25 |
| RPS23 | P62266 | 9.40E-04 | -0.44 | -1.36 |
| RPS6 | P62753 | 0.0086 | -0.41 | -1.32 |
| RRBP1 | Q9P2E9 | 3.20E-04 | -0.32 | -1.25 |
| RRP1 | P56182 | 0.0052 | -0.29 | -1.22 |
| RRP15 | Q9Y3B9 | 6.00E-05 | -0.33 | -1.26 |
| RRP1B | Q14684 | 7.90E-06 | -0.53 | -1.44 |
| RRS1 | Q15050 | 2.20E-05 | -0.36 | -1.29 |
| RSL1D1 | O76021 | 2.00E-04 | -0.56 | -1.47 |
| RTN3 | O95197 | 1.40E-05 | -0.46 | -1.38 |
| RUNX1T1 | Q06455 | 0.011 | -0.32 | -1.25 |
| S100A10 | P60903 | 2.10E-04 | -0.6 | -1.52 |
| SAFB2 | Q14151 | 5.70E-05 | -0.37 | -1.29 |
| SARAF | Q96BY9 | 0.0053 | -0.26 | -1.2 |
| SCD | O00767 | 1.40E-04 | -0.41 | -1.33 |
| SCIN | Q9Y6U3 | 0.0011 | -0.47 | -1.38 |
| SDAD1 | Q9NVU7 | 0.0061 | -0.41 | -1.33 |
| SDC1 | P18827 | 0.0012 | -0.6 | -1.52 |
| SDC2 | P34741 | 3.40E-04 | -0.55 | -1.47 |
| SDHAF3 | Q9NRP4 | 3.50E-06 | -0.53 | -1.44 |
| SEC61B | P60468 | 6.50E-04 | -0.33 | -1.25 |
| SEC62 | Q99442 | 0.0013 | -0.29 | -1.23 |
| SELENOF | O60613 | 2.00E-05 | -0.43 | -1.35 |
| SELENOH | Q8IZQ5 | 3.60E-04 | -0.29 | -1.22 |
| SELENOK | Q9Y6D0 | 0.0017 | -0.34 | -1.27 |
| SELENOM | Q8WWX9 | 0.0072 | -0.31 | -1.24 |
| SELENOT | P62341 | 1.90E-04 | -0.29 | -1.23 |
| SEMA4A | Q9H3S1 | 2.50E-06 | -0.56 | -1.47 |
| SEMA6A | Q9H2E6 | 0.0026 | -0.33 | -1.26 |
| 6-Sep | Q14141 | 0.0022 | -0.29 | -1.22 |
| SERF2 | P84101 | 0.0096 | -0.37 | -1.29 |

| | | | | |
|---|---|---|---|---|
| SERPINB3 | P29508 | 5.40E-04 | -0.56 | -1.47 |
| SERPINB4 | P48594 | 3.60E-07 | -0.51 | -1.43 |
| SGCB | Q16585 | 0.042 | -0.27 | -1.2 |
| SGO2 | Q562F6 | 0.022 | -0.3 | -1.23 |
| SH2D3A | Q9BRG2 | 3.50E-04 | -0.3 | -1.23 |
| SH2D4A | Q9H788 | 9.00E-06 | -0.46 | -1.37 |
| SH3RF1 | Q7Z6J0 | 0.0051 | -0.35 | -1.28 |
| SHB | Q15464 | 0.0037 | -0.27 | -1.2 |
| SHROOM4 | Q9ULL8 | 4.00E-04 | -0.55 | -1.46 |
| SIPA1L2 | Q9P2F8 | 7.30E-06 | -0.79 | -1.74 |
| SLC16A10 | Q8TF71 | 1.90E-04 | -0.54 | -1.45 |
| SLC20A2 | Q08357 | 0.0051 | -0.29 | -1.22 |
| SLC25A5 | P05141 | 1.00E-05 | -0.54 | -1.46 |
| SLC26A6 | Q9BXS9 | 0.0022 | -0.28 | -1.22 |
| SLC29A1 | Q99808 | 0.0084 | -0.46 | -1.37 |
| SLC29A2 | Q14542 | 0.0047 | -0.33 | -1.26 |
| SLC35A2 | P78381 | 8.70E-04 | -0.28 | -1.21 |
| SLC37A1 | P57057 | 5.50E-05 | -0.38 | -1.3 |
| SLC38A5 | Q8WUX1 | 5.30E-06 | -0.43 | -1.35 |
| SLC39A14 | Q15043 | 1.60E-04 | -0.37 | -1.29 |
| SLC39A8 | Q9C0K1 | 0.041 | -0.33 | -1.25 |
| SLC4A11 | Q8NBS3 | 2.00E-04 | -0.51 | -1.43 |
| SLC4A2 | P04920 | 4.00E-04 | -0.3 | -1.24 |
| SLC6A6 | P31641 | 0.034 | -0.28 | -1.22 |
| SLC7A3 | Q8WY07 | 0.0079 | -0.37 | -1.29 |
| SLCO4A1 | Q96BD0 | 0.016 | -0.32 | -1.25 |
| SLIT3 | O75094 | 7.80E-05 | -0.65 | -1.57 |
| SLITRK4 | Q8IW52 | 1.40E-04 | -0.39 | -1.31 |
| SNAPC1 | Q16533 | 0.0016 | -0.4 | -1.32 |
| SNCAIP | Q9Y6H5 | 0.0019 | -0.43 | -1.35 |
| SNTA1 | Q13424 | 0.0029 | -0.42 | -1.34 |
| SNX18 | Q96RF0 | 4.40E-04 | -0.28 | -1.21 |
| SOD2 | P04179 | 0.018 | -0.3 | -1.23 |
| SOX15 | O60248 | 5.00E-04 | -0.71 | -1.64 |
| SOX3 | P41225 | 0.014 | -0.34 | -1.27 |
| SOX4 | Q06945 | 1.20E-05 | -0.38 | -1.3 |
| SPARC | P09486 | 3.20E-04 | -0.36 | -1.28 |
| SPATA2 | Q9UM82 | 0.0025 | -0.27 | -1.2 |
| SPIDR | Q14159 | 0.0013 | -0.26 | -1.2 |
| SPIN2A | Q99865 | 0.0067 | -0.31 | -1.24 |
| SPINT1 | O43278 | 1.80E-04 | -0.32 | -1.25 |

| | | | | |
|---|---|---|---|---|
| SPRY2 | O43597 | 9.30E-05 | -0.48 | -1.4 |
| SPRY4 | Q9C004 | 4.50E-05 | -0.54 | -1.45 |
| SPTY2D1 | Q68D10 | 0.0012 | -0.76 | -1.69 |
| SREBF1 | P36956 | 0.0088 | -0.27 | -1.21 |
| SRP19 | P09132 | 0.0082 | -0.26 | -1.2 |
| ST14 | Q9Y5Y6 | 2.40E-04 | -0.33 | -1.25 |
| ST8SIA3 | O43173 | 7.00E-04 | -0.26 | -1.2 |
| STAB2 | Q8WWQ8 | 0.03 | -0.27 | -1.21 |
| STK17A | Q9UEE5 | 0.0061 | -0.32 | -1.25 |
| STK32C | Q86UX6 | 9.00E-04 | -0.54 | -1.45 |
| STOM | P27105 | 0.0076 | -0.36 | -1.28 |
| STXBP6 | Q8NFX7 | 4.80E-04 | -0.42 | -1.34 |
| SUCLG1 | P53597 | 2.00E-04 | -0.34 | -1.27 |
| SUCLG2 | Q96I99 | 0.0015 | -0.26 | -1.2 |
| SYAP1 | Q96A49 | 3.10E-04 | -0.27 | -1.21 |
| SYDE1 | Q6ZW31 | 0.0014 | -0.28 | -1.22 |
| SYNPO2 | Q9UMS6 | 0.0056 | -0.31 | -1.24 |
| SYT13 | Q7L8C5 | 0.043 | -0.28 | -1.21 |
| SYT2 | Q8N9I0 | 6.80E-07 | -0.71 | -1.64 |
| TACC2 | O95359 | 2.60E-04 | -0.28 | -1.22 |
| TAGLN | Q01995 | 0.041 | -0.29 | -1.22 |
| TAGLN2 | P37802 | 1.90E-05 | -0.4 | -1.32 |
| TANC2 | Q9HCD6 | 5.00E-04 | -0.54 | -1.46 |
| TBC1D2 | Q9BYX2 | 0.0025 | -0.32 | -1.25 |
| TCEAL2 | Q9H3H9 | 5.10E-07 | -2.21 | -4.63 |
| TCEAL7 | Q9BRU2 | 0.0021 | -0.27 | -1.21 |
| TCF25 | Q9BQ70 | 6.20E-05 | -0.3 | -1.23 |
| TCF7L1 | Q9HCS4 | 8.40E-04 | -0.33 | -1.25 |
| TCL1A | P56279 | 3.60E-04 | -0.81 | -1.76 |
| TCTN3 | Q6NUS6 | 6.70E-05 | -0.45 | -1.36 |
| TEK | Q02763 | 3.00E-05 | -0.43 | -1.35 |
| TFPI2 | P48307 | 0.014 | -0.46 | -1.38 |
| TFRC | P02786 | 5.80E-05 | -0.36 | -1.29 |
| THAP4 | Q8WY91 | 3.80E-04 | -0.28 | -1.21 |
| THSD7B | Q9C0I4 | 0.021 | -0.59 | -1.5 |
| THYN1 | Q9P016 | 6.10E-05 | -0.3 | -1.23 |
| TIE1 | P35590 | 6.70E-04 | -0.65 | -1.57 |
| TMA7 | Q9Y2S6 | 0.0051 | -0.45 | -1.36 |
| TMEM132A | Q24JP5 | 0.0017 | -0.28 | -1.21 |
| TMEM151B | Q8IW70 | 7.30E-04 | -0.35 | -1.28 |
| TMEM2 | Q9UHN6 | 4.00E-04 | -0.3 | -1.23 |

| | | | | |
|---|---|---|---|---|
| TMEM229A | B2RXF0 | 0.005 | -0.39 | -1.31 |
| TMEM55B | Q86T03 | 0.0012 | -0.35 | -1.28 |
| TMEM63A | O94886 | 0.0017 | -0.35 | -1.28 |
| TMPPE | Q6ZT21 | 4.20E-04 | -0.44 | -1.35 |
| TMSB10 | P63313 | 0.0021 | -0.27 | -1.21 |
| TMSB4Y | O14604 | 3.30E-05 | -0.71 | -1.63 |
| TMX2 | Q9Y320 | 1.30E-04 | -0.28 | -1.22 |
| TNFRSF10C | O14798 | 0.0061 | -0.28 | -1.21 |
| TNFRSF12A | Q9NP84 | 3.90E-04 | -0.38 | -1.3 |
| TNFRSF8 | P28908 | 0.0018 | -0.47 | -1.39 |
| TNFSF9 | P41273 | 0.014 | -0.45 | -1.37 |
| TNRC6A | Q8NDV7 | 1.80E-05 | -0.34 | -1.26 |
| TNS1 | Q9HBL0 | 0.0012 | -0.31 | -1.24 |
| TOMM20 | Q15388 | 2.60E-04 | -0.35 | -1.28 |
| TOMM70 | O94826 | 1.10E-04 | -0.28 | -1.21 |
| TOP1 | P11387 | 0.0043 | -0.5 | -1.42 |
| TPM1 | P09493 | 6.40E-04 | -0.36 | -1.28 |
| TPM2 | P07951 | 1.50E-04 | -0.38 | -1.3 |
| TRA2B | P62995 | 3.60E-04 | -0.33 | -1.26 |
| TRANK1 | O15050 | 0.012 | -0.5 | -1.41 |
| TRIM41 | Q8WV44 | 0.0097 | -0.26 | -1.2 |
| TRIM6 | Q9C030 | 2.90E-04 | -0.43 | -1.35 |
| TRO | Q12816 | 1.40E-04 | -0.36 | -1.28 |
| TSHZ3 | Q63HK5 | 9.20E-04 | -0.27 | -1.21 |
| TSR3 | Q9UJK0 | 9.70E-05 | -0.47 | -1.38 |
| TTC39B | Q5VTQ0 | 0.0087 | -0.4 | -1.32 |
| TTC39C | Q8N584 | 1.50E-07 | -0.91 | -1.88 |
| TTC7A | Q9ULT0 | 0.0062 | -0.26 | -1.2 |
| TTF1 | Q15361 | 1.10E-04 | -0.34 | -1.26 |
| TUBB3 | Q13509 | 0.0076 | -0.27 | -1.2 |
| TUBB6 | Q9BUF5 | 4.20E-04 | -0.32 | -1.25 |
| TWISTNB | Q3B726 | 7.30E-05 | -0.31 | -1.24 |
| UBA52 | P62987 | 0.04 | -0.28 | -1.22 |
| UBIAD1 | Q9Y5Z9 | 8.20E-05 | -0.45 | -1.36 |
| UGT3A2 | Q3SY77 | 1.30E-04 | -0.39 | -1.31 |
| UNC5B | Q8IZJ1 | 0.014 | -0.31 | -1.24 |
| UPP1 | Q16831 | 0.025 | -0.26 | -1.2 |
| UROD | P06132 | 0.0034 | -0.27 | -1.21 |
| USP35 | Q9P2H5 | 0.0049 | -0.39 | -1.31 |
| UTF1 | Q5T230 | 0.0053 | -0.55 | -1.46 |
| UTP11 | Q9Y3A2 | 0.012 | -0.51 | -1.42 |

| | | | | |
|---|---|---|---|---|
| UXS1 | Q8NBZ7 | 2.90E-04 | -0.31 | -1.24 |
| VCAN | P13611 | 0.001 | -0.29 | -1.22 |
| VCY | O14598 | 0.0087 | -0.28 | -1.22 |
| VIM | P08670 | 2.00E-06 | -0.87 | -1.83 |
| VOPP1 | Q96AW1 | 0.0038 | -0.29 | -1.22 |
| VSIG10 | Q8N0Z9 | 5.50E-04 | -0.36 | -1.28 |
| WDR74 | Q6RFH5 | 4.30E-04 | -0.28 | -1.21 |
| WFS1 | O76024 | 2.20E-04 | -0.33 | -1.26 |
| XPC | Q01831 | 0.0071 | -0.72 | -1.65 |
| XYLT1 | Q86Y38 | 2.00E-04 | -0.41 | -1.33 |
| YME1L1 | Q96TA2 | 2.40E-04 | -0.27 | -1.2 |
| ZBTB3 | Q9H5J0 | 8.50E-05 | -0.33 | -1.25 |
| ZC3H3 | Q8IXZ2 | 0.0051 | -0.36 | -1.28 |
| ZCCHC17 | Q9NP64 | 4.80E-05 | -0.31 | -1.24 |
| ZCCHC9 | Q8N567 | 4.50E-05 | -0.32 | -1.24 |
| ZDBF2 | Q9HCK1 | 0.0097 | -0.36 | -1.28 |
| ZDHHC12 | Q96GR4 | 0.0085 | -0.32 | -1.25 |
| ZDHHC15 | Q96MV8 | 0.0044 | -0.62 | -1.54 |
| ZDHHC22 | Q8N966 | 2.80E-05 | -0.72 | -1.65 |
| ZDHHC3 | Q9NYG2 | 4.60E-04 | -0.34 | -1.27 |
| ZFP42 | Q96MM3 | 1.20E-04 | -0.5 | -1.41 |
| ZFP62 | Q8NB50 | 0.049 | -0.41 | -1.33 |
| ZIC3 | O60481 | 0.0024 | -0.38 | -1.3 |
| ZIK1 | Q3SY52 | 0.0015 | -0.3 | -1.23 |
| ZMAT2 | Q96NC0 | 0.0012 | -0.36 | -1.28 |
| ZMPSTE24 | O75844 | 6.00E-04 | -0.27 | -1.21 |
| ZMYND19 | Q96E35 | 0.0034 | -0.28 | -1.21 |
| ZNF107 | Q9UII5 | 0.016 | -0.3 | -1.23 |
| ZNF112 | Q9UJU3 | 0.022 | -0.36 | -1.28 |
| ZNF124 | Q15973 | 1.10E-04 | -0.41 | -1.33 |
| ZNF136 | P52737 | 0.0041 | -0.31 | -1.24 |
| ZNF141 | Q15928 | 1.90E-04 | -0.47 | -1.39 |
| ZNF208 | O43345 | 0.026 | -1.65 | -3.15 |
| ZNF22 | P17026 | 0.028 | -0.37 | -1.29 |
| ZNF232 | Q9UNY5 | 0.0097 | -0.53 | -1.44 |
| ZNF282 | Q9UDV7 | 0.034 | -0.33 | -1.26 |
| ZNF296 | Q8WUU4 | 4.50E-04 | -0.37 | -1.3 |
| ZNF354A | O60765 | 4.60E-04 | -0.65 | -1.57 |
| ZNF397 | Q8NF99 | 0.0023 | -0.37 | -1.29 |
| ZNF398 | Q8TD17 | 5.10E-06 | -0.43 | -1.34 |
| ZNF414 | Q96IQ9 | 0.029 | -0.28 | -1.21 |

| | | | | |
|---|---|---|---|---|
| ZNF429 | Q86V71 | 0.0013 | -0.67 | -1.59 |
| ZNF460 | Q14592 | 0.0038 | -0.35 | -1.28 |
| ZNF48 | Q96MX3 | 0.0013 | -0.41 | -1.33 |
| ZNF485 | Q8NCK3 | 0.029 | -0.39 | -1.31 |
| ZNF512 | Q96ME7 | 0.004 | -1.22 | -2.33 |
| ZNF512B | Q96KM6 | 0.02 | -0.9 | -1.87 |
| ZNF525 | Q8N782 | 0.011 | -0.6 | -1.51 |
| ZNF552 | Q9H707 | 0.0035 | -0.44 | -1.35 |
| ZNF561 | Q8N587 | 0.0045 | -0.36 | -1.28 |
| ZNF587 | Q96SQ5 | 0.034 | -0.33 | -1.25 |
| ZNF589 | Q86UQ0 | 6.00E-04 | -0.28 | -1.21 |
| ZNF595 | Q8IYB9 | 0.012 | -0.39 | -1.31 |
| ZNF644 | Q9H582 | 0.001 | -0.29 | -1.22 |
| ZNF66 | Q6ZN08 | 0.0087 | -0.61 | -1.52 |
| ZNF660 | Q6AZW8 | 0.0012 | -0.42 | -1.34 |
| ZNF668 | Q96K58 | 0.019 | -0.27 | -1.21 |
| ZNF708 | P17019 | 0.026 | -0.56 | -1.47 |
| ZNF71 | Q9NQZ8 | 0.015 | -0.28 | -1.21 |
| ZNF724 | A8MTY0 | 0.024 | -0.39 | -1.31 |
| ZNF75A | Q96N20 | 0.044 | -0.29 | -1.23 |
| ZNF761 | Q86XN6 | 0.0074 | -0.4 | -1.32 |
| ZNF768 | Q9H5H4 | 1.60E-04 | -0.3 | -1.23 |
| ZNF770 | Q6IQ21 | 0.018 | -0.75 | -1.68 |
| ZNF785 | A8K8V0 | 0.018 | -0.31 | -1.24 |
| ZNF787 | Q6DD87 | 2.00E-04 | -0.38 | -1.3 |
| ZNF792 | Q3KQV3 | 0.029 | -0.26 | -1.2 |
| ZNF805 | Q5CZA5 | 0.0054 | -0.3 | -1.23 |
| ZNF808 | Q8N4W9 | 0.024 | -0.37 | -1.3 |
| ZNF813 | Q6ZN06 | 6.50E-04 | -0.34 | -1.26 |
| ZNF814 | B7Z6K7 | 0.033 | -0.62 | -1.54 |
| ZNF845 | Q96IR2 | 0.0055 | -0.4 | -1.32 |
| ZNF850 | A8MQ14 | 0.0048 | -0.68 | -1.6 |
| ZNF90 | Q03938 | 0.0093 | -0.6 | -1.52 |
| ZNF93 | P35789 | 0.037 | -0.6 | -1.51 |
| ZNHIT1 | O43257 | 0.0038 | -0.43 | -1.35 |
| ZNHIT3 | Q15649 | 0.013 | -0.46 | -1.38 |
| ZNRD1 | Q9P1U0 | 6.30E-04 | -0.43 | -1.35 |
| ZSCAN10 | Q96SZ4 | 0.0021 | -0.35 | -1.28 |
| ZYX | Q15942 | 1.30E-04 | -0.32 | -1.25 |

| **Supplementary Table 3A: *KDM6A*KO upregulated phospho-protein expression by mass spec**<br>[pVal = p-value; logFC = log of the fold change; FC = fold change in expression] | | | | |
|---|---|---|---|---|
| **Protein** | **Accession** | **pVal** | **logFC** | **FC** |
| AAK1 | Q2M2I8 | 4.50E-05 | 0.55 | 1.47 |
| ABCC4 | O15439 | 0.005 | 0.38 | 1.3 |
| ABCD3 | P28288 | 0.044 | 0.32 | 1.25 |
| ABI1 | Q8IZP0 | 0.003 | 0.68 | 1.61 |
| ABLIM2 | Q6H8Q1 | 5.70E-05 | 0.52 | 1.43 |
| ADAM22 | Q9P0K1 | 0.0037 | 0.4 | 1.32 |
| ADAR | P55265 | 0.03 | 0.36 | 1.28 |
| ADRM1 | Q16186 | 9.80E-04 | 0.94 | 1.92 |
| AFF4 | Q9UHB7 | 1.60E-04 | 0.45 | 1.37 |
| AGAP1 | Q9UPQ3 | 0.0055 | 0.49 | 1.41 |
| AGAP3 | Q96P47 | 1.00E-04 | 0.62 | 1.53 |
| AGFG1 | P52594 | 1.40E-04 | 1.22 | 2.33 |
| AHNAK | Q09666 | 0.0032 | 0.74 | 1.67 |
| AIF1L | Q9BQI0 | 4.50E-05 | 1.24 | 2.37 |
| AJUBA | Q96IF1 | 0.0016 | 0.35 | 1.28 |
| AKAP1 | Q92667 | 0.02 | 0.4 | 1.32 |
| AKAP10 | O43572 | 0.003 | 0.44 | 1.36 |
| AKAP9 | Q99996 | 9.70E-05 | 1.16 | 2.23 |
| AKT1 | P31749 | 0.0098 | 0.28 | 1.22 |
| AKT2 | P31751 | 0.025 | 0.51 | 1.42 |
| ALG3 | Q92685 | 0.0088 | 0.48 | 1.4 |
| ALMS1 | Q8TCU4 | 0.033 | 0.27 | 1.2 |
| ALPK3 | Q96L96 | 0.011 | 0.36 | 1.29 |
| AMMECR1L | Q6DCA0 | 4.40E-04 | 0.6 | 1.51 |
| AMOTL1 | Q8IY63 | 0.0056 | 0.33 | 1.26 |
| ANK1 | P16157 | 9.50E-05 | 0.71 | 1.64 |
| ANK3 | Q12955 | 0.013 | 0.26 | 1.2 |
| ANKHD1 | Q8IWZ3 | 0.0031 | 0.43 | 1.34 |
| ANKRD54 | Q6NXT1 | 1.60E-06 | 0.9 | 1.87 |
| ANP32A | P39687 | 0.013 | 0.35 | 1.27 |
| ANP32B | Q92688 | 0.017 | 0.64 | 1.56 |
| ANTXR1 | Q9H6X2 | 4.60E-04 | 0.56 | 1.48 |
| AP3D1 | O14617 | 2.40E-04 | 0.78 | 1.72 |
| APBA3 | O96018 | 0.019 | 0.26 | 1.2 |
| APBB2 | Q92870 | 0.0041 | 0.67 | 1.6 |
| ARAF | P10398 | 0.0076 | 0.36 | 1.29 |
| ARFGAP2 | Q8N6H7 | 0.014 | 0.47 | 1.39 |
| ARFGEF1 | Q9Y6D6 | 0.0035 | 0.39 | 1.31 |

| | | | | |
|---|---|---|---|---|
| ARFGEF2 | Q9Y6D5 | 2.30E-05 | 0.82 | 1.77 |
| ARFIP1 | P53367 | 0.0088 | 0.31 | 1.24 |
| ARHGAP1 | Q07960 | 0.0037 | 0.61 | 1.52 |
| ARHGAP23 | Q9P227 | 2.20E-04 | 0.45 | 1.37 |
| ARHGAP32 | A7KAX9 | 0.0025 | 0.48 | 1.39 |
| ARHGAP39 | Q9C0H5 | 0.004 | 0.5 | 1.41 |
| ARHGEF1 | Q92888 | 0.0011 | 1.46 | 2.76 |
| ARHGEF12 | Q9NZN5 | 7.20E-04 | 0.65 | 1.57 |
| ARHGEF16 | Q5VV41 | 0.0045 | 0.38 | 1.3 |
| ARHGEF18 | Q6ZSZ5 | 0.003 | 0.4 | 1.32 |
| ARHGEF25 | Q86VW2 | 6.50E-06 | 1.45 | 2.74 |
| ARHGEF5 | Q12774 | 7.80E-06 | 0.75 | 1.68 |
| ARID2 | Q68CP9 | 0.026 | 0.33 | 1.26 |
| ARL2 | P36404 | 0.002 | 0.54 | 1.45 |
| ASAP1 | Q9ULH1 | 0.029 | 0.34 | 1.26 |
| ASCC3 | Q8N3C0 | 2.30E-04 | 0.7 | 1.63 |
| ASF1B | Q9NVP2 | 0.048 | 0.39 | 1.31 |
| ASPSCR1 | Q9BZE9 | 0.011 | 0.26 | 1.2 |
| ATAT1 | Q5SQI0 | 5.20E-06 | 0.96 | 1.95 |
| ATF6 | P18850 | 0.0076 | 0.77 | 1.7 |
| ATG13 | O75143 | 0.035 | 0.59 | 1.51 |
| ATP13A1 | Q9HD20 | 0.01 | 0.41 | 1.32 |
| ATR | Q13535 | 0.031 | 0.3 | 1.23 |
| AUNIP | Q9H7T9 | 7.00E-04 | 0.74 | 1.67 |
| AUTS2 | Q8WXX7 | 0.0019 | 0.47 | 1.39 |
| BAG3 | O95817 | 1.60E-04 | 0.71 | 1.63 |
| BAG6 | P46379 | 0.046 | 0.27 | 1.21 |
| BAP18 | Q8IXM2 | 0.024 | 0.29 | 1.22 |
| BBX | Q8WY36 | 0.0094 | 0.32 | 1.25 |
| BCAP31 | P51572 | 5.80E-04 | 1.15 | 2.22 |
| BCL9 | O00512 | 7.40E-04 | 0.4 | 1.32 |
| BET1L | Q9NYM9 | 0.0018 | 0.48 | 1.4 |
| BICDL2 | A1A5D9 | 0.028 | 0.36 | 1.29 |
| BMI1 | P35226 | 3.10E-04 | 0.71 | 1.63 |
| BMP2K | Q9NSY1 | 0.0033 | 0.4 | 1.32 |
| BNIP2 | Q12982 | 0.0025 | 0.79 | 1.73 |
| BRF1 | Q92994 | 0.011 | 0.52 | 1.44 |
| BRSK2 | Q8IWQ3 | 0.011 | 0.43 | 1.35 |
| BSN | Q9UPA5 | 3.10E-05 | 0.94 | 1.92 |
| BUB1 | O43683 | 6.20E-06 | 0.74 | 1.68 |
| C12orf45 | Q8N5I9 | 0.018 | 0.63 | 1.54 |

| | | | | |
|---|---|---|---|---|
| C1orf226 | A1L170 | 0.0023 | 0.54 | 1.45 |
| C2CD4C | Q8TF44 | 0.0027 | 0.52 | 1.43 |
| C2orf69 | Q8N8R5 | 0.007 | 0.85 | 1.8 |
| C2orf74 | A8MZ97 | 0.0093 | 0.46 | 1.37 |
| C6orf132 | Q5T0Z8 | 3.80E-04 | 0.54 | 1.45 |
| C7orf43 | Q8WVR3 | 0.006 | 0.39 | 1.31 |
| C9orf142 | Q9BUH6 | 0.028 | 0.32 | 1.25 |
| CA8 | P35219 | 0.018 | 0.47 | 1.39 |
| CAMKK2 | Q96RR4 | 0.0012 | 0.79 | 1.73 |
| CAMSAP1 | Q5T5Y3 | 0.0015 | 0.58 | 1.5 |
| CAMSAP3 | Q9P1Y5 | 0.0018 | 0.33 | 1.26 |
| CARMIL1 | Q5VZK9 | 0.0083 | 0.4 | 1.32 |
| CASC5_HUMAN | NA | 0.021 | 0.33 | 1.26 |
| CASP3 | P42574 | 0.0035 | 0.46 | 1.38 |
| CBX5 | P45973 | 1.90E-05 | 1.84 | 3.59 |
| CCDC117 | Q8IWD4 | 1.60E-06 | 1.26 | 2.39 |
| CCDC88A | Q3V6T2 | 0.0029 | 0.33 | 1.26 |
| CCDC9 | Q9Y3X0 | 0.0073 | 0.32 | 1.24 |
| CCM2 | Q9BSQ5 | 6.80E-04 | 1.03 | 2.04 |
| CCNE1 | P24864 | 4.80E-04 | 0.77 | 1.7 |
| CDC20 | Q12834 | 2.20E-04 | 0.89 | 1.85 |
| CDC25A | P30304 | 0.032 | 0.31 | 1.24 |
| CDC25C | P30307 | 0.035 | 0.41 | 1.32 |
| CDC42BPA | Q5VT25 | 0.0024 | 0.97 | 1.96 |
| CDC42EP1 | Q00587 | 1.80E-04 | 1.13 | 2.18 |
| CDC42EP3 | Q9UKI2 | 0.0038 | 0.78 | 1.72 |
| CDC6 | Q99741 | 0.0046 | 0.59 | 1.51 |
| CDCA3 | Q99618 | 0.0033 | 0.54 | 1.45 |
| CDCA8 | Q53HL2 | 0.019 | 0.32 | 1.25 |
| CDK16 | Q00536 | 0.0036 | 0.46 | 1.38 |
| CDK17 | Q00537 | 0.02 | 0.98 | 1.97 |
| CDK9 | P50750 | 0.0014 | 0.69 | 1.61 |
| CDKL5 | O76039 | 0.013 | 0.28 | 1.21 |
| CDR2 | Q01850 | 0.0019 | 0.95 | 1.93 |
| CENPF | P49454 | 0.0075 | 0.31 | 1.24 |
| CENPI | Q92674 | 0.031 | 0.32 | 1.25 |
| CENPT | Q96BT3 | 0.0052 | 0.89 | 1.85 |
| CEP112 | Q8N8E3 | 5.90E-04 | 0.51 | 1.42 |
| CEP128 | Q6ZU80 | 0.029 | 0.49 | 1.41 |
| CEP170 | Q5SW79 | 0.039 | 0.28 | 1.21 |
| CEP295 | Q9C0D2 | 0.017 | 0.28 | 1.21 |

| | | | | |
|---|---|---|---|---|
| CEP41 | Q9BYV8 | 0.0036 | 1.4 | 2.64 |
| CEP97 | Q8IW35 | 0.046 | 0.28 | 1.22 |
| CFAP97 | Q9P2B7 | 1.30E-05 | 0.8 | 1.74 |
| CFL2 | Q9Y281 | 0.0012 | 1.85 | 3.6 |
| CHD1 | O14646 | 0.028 | 0.35 | 1.27 |
| CHD7 | Q9P2D1 | 8.80E-04 | 0.65 | 1.57 |
| CHEK1 | O14757 | 0.0049 | 0.52 | 1.43 |
| CHGA | P10645 | 0.021 | 0.44 | 1.35 |
| CISD2 | Q8N5K1 | 0.045 | 0.52 | 1.44 |
| CLASP1 | Q7Z460 | 1.50E-04 | 0.69 | 1.61 |
| CLASP2 | O75122 | 0.0013 | 0.78 | 1.71 |
| CLDN3 | O15551 | 2.00E-04 | 0.56 | 1.47 |
| CLIC4 | Q9Y696 | 4.60E-04 | 0.49 | 1.41 |
| CLIP1 | P30622 | 2.50E-05 | 1.23 | 2.34 |
| CLIP2 | Q9UDT6 | 0.0027 | 0.54 | 1.46 |
| CNNM3 | Q8NE01 | 0.0031 | 0.42 | 1.34 |
| CNNM4 | Q6P4Q7 | 0.0012 | 0.54 | 1.45 |
| CNOT2 | Q9NZN8 | 5.40E-05 | 0.81 | 1.76 |
| CNTLN | Q9NXG0 | 0.003 | 0.44 | 1.36 |
| CRMP1 | Q14194 | 5.40E-04 | 0.98 | 1.97 |
| CRTC1 | Q6UUV9 | 0.0041 | 0.39 | 1.31 |
| CSNK1A1 | P48729 | 6.60E-04 | 0.9 | 1.86 |
| CSNK1D | P48730 | 0.0025 | 1.11 | 2.17 |
| CSNK1G3 | Q9Y6M4 | 0.0024 | 0.4 | 1.32 |
| CTCF | P49711 | 0.012 | 0.38 | 1.3 |
| CTDSPL2 | Q05D32 | 6.20E-05 | 0.84 | 1.79 |
| CTNNB1 | P35222 | 0.011 | 0.35 | 1.27 |
| CU025_HUMAN | NA | 7.90E-04 | 0.96 | 1.95 |
| CXorf38 | Q8TB03 | 1.30E-04 | 0.78 | 1.72 |
| CYP2S1 | Q96SQ9 | 0.024 | 0.5 | 1.41 |
| DAGLB | Q8NCG7 | 0.0043 | 0.38 | 1.3 |
| DCAF10 | Q5QP82 | 0.0029 | 0.3 | 1.24 |
| DCAF11 | Q8TEB1 | 0.0036 | 0.51 | 1.42 |
| DCAF5 | Q96JK2 | 0.032 | 0.34 | 1.27 |
| DCUN1D4 | Q92564 | 3.80E-04 | 0.56 | 1.47 |
| DCX | O43602 | 0.0028 | 0.56 | 1.48 |
| DDX11L8 | A8MPP1 | 0.023 | 0.48 | 1.4 |
| DDX17 | Q92841 | 2.90E-05 | 0.78 | 1.72 |
| DDX39A | O00148 | 0.0013 | 0.41 | 1.33 |
| DDX3X | O00571 | 0.0013 | 0.81 | 1.76 |
| DDX5 | P17844 | 0.042 | 0.64 | 1.55 |

| | | | | |
|---|---|---|---|---|
| DDX52 | Q9Y2R4 | 0.0018 | 1.41 | 2.65 |
| DDX59 | Q5T1V6 | 2.50E-04 | 0.85 | 1.8 |
| DDX6 | P26196 | 1.90E-04 | 0.8 | 1.74 |
| DEF6 | Q9H4E7 | 0.0063 | 0.51 | 1.42 |
| DENND1A | Q8TEH3 | 4.70E-06 | 0.97 | 1.96 |
| DENND5B | Q6ZUT9 | 5.50E-04 | 0.38 | 1.3 |
| DENND6A | Q8IWF6 | 0.002 | 0.98 | 1.97 |
| DHX57 | Q6P158 | 0.0078 | 0.73 | 1.65 |
| DICER1 | Q9UPY3 | 0.046 | 0.29 | 1.22 |
| DIP2C | Q9Y2E4 | 0.0043 | 0.62 | 1.53 |
| DMAP1 | Q9NPF5 | 0.014 | 0.44 | 1.35 |
| DMXL1 | Q9Y485 | 0.011 | 0.41 | 1.33 |
| DNM1L | O00429 | 1.70E-04 | 0.8 | 1.75 |
| DNPH1 | O43598 | 0.013 | 0.35 | 1.27 |
| DOCK6 | Q96HP0 | 0.0056 | 0.57 | 1.48 |
| DOCK7 | Q96N67 | 0.011 | 0.49 | 1.4 |
| DOK1 | Q99704 | 0.003 | 0.42 | 1.34 |
| DPF2 | Q92785 | 0.016 | 0.26 | 1.2 |
| DPYSL2 | Q16555 | 7.00E-06 | 1.17 | 2.25 |
| DPYSL3 | Q14195 | 0.027 | 0.29 | 1.22 |
| DSA2D_HUMAN | C9JQL5 | 0.022 | 0.51 | 1.43 |
| DSC2 | Q02487 | 0.006 | 0.57 | 1.48 |
| DSN1 | Q9H410 | 0.024 | 0.35 | 1.28 |
| DYDC1 | Q8WWB3 | 0.034 | 0.61 | 1.52 |
| DYNC1LI2 | O43237 | 0.0018 | 0.43 | 1.35 |
| ECM29 | Q5VYK3 | 3.70E-04 | 0.69 | 1.61 |
| EDC3 | Q96F86 | 0.0072 | 0.32 | 1.25 |
| EFHD2 | Q96C19 | 2.90E-04 | 0.51 | 1.42 |
| EFS | O43281 | 0.05 | 0.47 | 1.39 |
| EGLN1 | Q9GZT9 | 8.90E-05 | 1.05 | 2.07 |
| EIF2B4 | Q9UI10 | 0.026 | 0.31 | 1.24 |
| EIF3A | Q14152 | 3.80E-04 | 0.66 | 1.58 |
| EIF4ENIF1 | Q9NRA8 | 1.90E-04 | 0.76 | 1.69 |
| EIF4G2 | P78344 | 1.10E-04 | 0.9 | 1.86 |
| ENSA | O43768 | 2.10E-04 | 1.13 | 2.19 |
| EPB41 | P11171 | 0.0034 | 0.37 | 1.29 |
| EPB41L2 | O43491 | 0.0018 | 0.45 | 1.37 |
| EPB41L4B | Q9H329 | 0.015 | 0.49 | 1.4 |
| EPHA1 | P21709 | 0.0044 | 0.54 | 1.46 |
| EPHX3 | Q9H6B9 | 0.032 | 0.26 | 1.2 |
| EPN3 | Q9H201 | 0.0046 | 0.85 | 1.8 |

| | | | | |
|---|---|---|---|---|
| ERC1 | Q8IUD2 | 0.0097 | 1.09 | 2.12 |
| ERC2 | O15083 | 0.002 | 0.51 | 1.42 |
| ERF | P50548 | 4.70E-04 | 0.91 | 1.88 |
| ERG | P11308 | 0.011 | 0.73 | 1.66 |
| ESCO2 | Q56NI9 | 8.90E-04 | 0.51 | 1.42 |
| ESPN | B1AK53 | 0.043 | 0.52 | 1.44 |
| ETV3 | P41162 | 1.50E-05 | 0.92 | 1.9 |
| EXOC2 | Q96KP1 | 4.70E-06 | 1.73 | 3.32 |
| EXOC7 | Q9UPT5 | 2.40E-04 | 1.33 | 2.52 |
| EYA3 | Q99504 | 0.03 | 0.29 | 1.22 |
| EZH2 | Q15910 | 0.05 | 0.29 | 1.22 |
| FA65B_HUMAN | NA | 0.002 | 0.63 | 1.55 |
| FAM109A | Q8N4B1 | 8.00E-05 | 0.58 | 1.5 |
| FAM117A | Q9C073 | 0.026 | 0.26 | 1.2 |
| FAM122A | Q96E09 | 9.10E-04 | 0.37 | 1.29 |
| FAM129A | Q9BZQ8 | 0.02 | 0.27 | 1.21 |
| FAM53B | Q14153 | 0.002 | 0.44 | 1.35 |
| FAM83H | Q6ZRV2 | 0.0052 | 0.38 | 1.3 |
| FBXO28 | Q9NVF7 | 0.034 | 0.32 | 1.25 |
| FBXO42 | Q6P3S6 | 2.90E-04 | 0.63 | 1.54 |
| FBXW5 | Q969U6 | 0.021 | 0.29 | 1.22 |
| FHOD1 | Q9Y613 | 0.021 | 0.3 | 1.23 |
| FHOD3 | Q2V2M9 | 0.013 | 1.31 | 2.48 |
| FIGN | Q5HY92 | 0.0013 | 0.45 | 1.36 |
| FKBP8 | Q14318 | 0.028 | 0.62 | 1.54 |
| FLYWCH2 | Q96CP2 | 0.0081 | 0.49 | 1.41 |
| FNBP1L | Q5T0N5 | 0.0042 | 0.43 | 1.34 |
| FOXJ3 | Q9UPW0 | 0.0025 | 0.53 | 1.44 |
| FRMD4A | Q9P2Q2 | 0.0047 | 0.69 | 1.61 |
| FRMD8 | Q9BZ67 | 0.0066 | 0.51 | 1.43 |
| FRY | Q5TBA9 | 6.60E-04 | 0.64 | 1.55 |
| FSD1 | Q9BTV5 | 1.60E-05 | 0.75 | 1.68 |
| FUBP3 | Q96I24 | 2.40E-04 | 0.53 | 1.45 |
| G0S2 | P27469 | 0.0044 | 0.93 | 1.9 |
| GAB2 | Q9UQC2 | 0.0091 | 0.36 | 1.28 |
| GAPVD1 | Q14C86 | 9.00E-05 | 0.75 | 1.69 |
| GIPC1 | O14908 | 0.0068 | 0.37 | 1.3 |
| GIT1 | Q9Y2X7 | 0.001 | 0.44 | 1.36 |
| GKAP1 | Q5VSY0 | 0.0011 | 0.53 | 1.45 |
| GLI2 | P10070 | 0.0064 | 0.35 | 1.28 |
| GLI3 | P10071 | 1.20E-04 | 0.84 | 1.79 |

| | | | | |
|---|---|---|---|---|
| GMPS | P49915 | 0.01 | 0.43 | 1.34 |
| GORASP2 | Q9H8Y8 | 3.10E-04 | 0.48 | 1.4 |
| GPBP1 | Q86WP2 | 0.0015 | 0.36 | 1.28 |
| GPRC5B | Q9NZH0 | 0.0054 | 0.82 | 1.77 |
| GRAMD1B | Q3KR37 | 5.70E-04 | 1.03 | 2.04 |
| GRB10 | Q13322 | 1.10E-05 | 1.19 | 2.28 |
| GSPT2 | Q8IYD1 | 0.023 | 0.63 | 1.55 |
| GTF2F1 | P35269 | 0.029 | 0.49 | 1.4 |
| GTF2F2 | P13984 | 0.032 | 1.13 | 2.19 |
| GYS1 | P13807 | 6.50E-04 | 0.52 | 1.44 |
| H2AFX | P16104 | 7.60E-04 | 0.65 | 1.57 |
| H2AFY | O75367 | 0.007 | 0.34 | 1.26 |
| HADH | Q16836 | 0.02 | 0.93 | 1.9 |
| HAUS6 | Q7Z4H7 | 0.0033 | 0.39 | 1.31 |
| HAUS8 | Q9BT25 | 4.20E-04 | 1.65 | 3.14 |
| HEATR6 | Q6AI08 | 0.01 | 0.57 | 1.48 |
| HECTD1 | Q9ULT8 | 0.018 | 0.28 | 1.21 |
| HERPUD2 | Q9BSE4 | 0.03 | 0.6 | 1.51 |
| HIST3H3 | Q16695 | 0.0063 | 0.99 | 1.99 |
| HIVEP1 | P15822 | 0.0095 | 0.3 | 1.23 |
| HN1L_HUMAN | NA | 4.00E-04 | 0.85 | 1.81 |
| HNRNPA1 | P09651 | 0.0015 | 0.59 | 1.51 |
| HNRNPA2B1 | P22626 | 0.028 | 0.27 | 1.21 |
| HNRNPH1 | P31943 | 0.0017 | 0.93 | 1.91 |
| HPS5 | Q9UPZ3 | 0.026 | 0.4 | 1.32 |
| HSPA12B | Q96MM6 | 0.0078 | 0.49 | 1.4 |
| ID2 | Q02363 | 0.0014 | 1.09 | 2.13 |
| IFITM2 | Q01629 | 0.047 | 0.36 | 1.29 |
| IFT88 | Q13099 | 0.0021 | 0.67 | 1.59 |
| IGF2BP3 | O00425 | 0.015 | 0.26 | 1.2 |
| IGF2R | P11717 | 0.011 | 0.38 | 1.3 |
| IKBKAP | O95163 | 0.023 | 0.26 | 1.2 |
| IKZF4 | Q9H2S9 | 0.047 | 0.4 | 1.32 |
| ILKAP | Q9H0C8 | 0.0065 | 0.3 | 1.23 |
| INF2 | Q27J81 | 0.003 | 0.45 | 1.36 |
| INPP5F | Q9Y2H2 | 0.0078 | 0.34 | 1.27 |
| INPPL1 | O15357 | 2.10E-04 | 1.18 | 2.26 |
| INTS12 | Q96CB8 | 0.0049 | 0.44 | 1.36 |
| INTS3 | Q68E01 | 0.012 | 0.35 | 1.28 |
| IQGAP3 | Q86VI3 | 0.014 | 0.79 | 1.73 |
| IRS2 | Q9Y4H2 | 0.025 | 0.27 | 1.2 |

| | | | | |
|---|---|---|---|---|
| ITPR1 | Q14643 | 0.0019 | 0.59 | 1.51 |
| ITPR3 | Q14573 | 9.40E-04 | 0.66 | 1.58 |
| IVNS1ABP | Q9Y6Y0 | 5.60E-04 | 0.45 | 1.36 |
| JMY | Q8N9B5 | 0.03 | 0.29 | 1.22 |
| KANSL2 | Q9H9L4 | 0.015 | 0.39 | 1.31 |
| KAT14 | Q9H8E8 | 9.70E-04 | 1.02 | 2.03 |
| KAT6A | Q92794 | 0.021 | 0.37 | 1.29 |
| KCNAB2 | Q13303 | 0.013 | 0.47 | 1.38 |
| KCNN3 | Q9UGI6 | 1.30E-05 | 1.07 | 2.1 |
| KCTD15 | Q96SI1 | 0.0012 | 1.46 | 2.75 |
| KDR | P35968 | 0.0042 | 0.63 | 1.55 |
| KHDRBS1 | Q07666 | 2.80E-04 | 0.56 | 1.47 |
| KIAA0556 | O60303 | 0.021 | 0.33 | 1.25 |
| KIAA0753 | Q2KHM9 | 8.20E-04 | 0.48 | 1.39 |
| KIAA1191 | Q96A73 | 9.40E-05 | 0.73 | 1.66 |
| KIAA1211 | Q6ZU35 | 0.033 | 0.51 | 1.43 |
| KIAA1217 | Q5T5P2 | 0.0066 | 0.32 | 1.25 |
| KIAA1614 | Q5VZ46 | 0.0067 | 0.63 | 1.55 |
| KIF13B | Q9NQT8 | 0.021 | 0.36 | 1.28 |
| KIF14 | Q15058 | 0.039 | 0.27 | 1.21 |
| KIF18B | Q86Y91 | 0.0034 | 0.31 | 1.24 |
| KIF20A | O95235 | 9.90E-04 | 0.71 | 1.63 |
| KIF22 | Q14807 | 0.0064 | 0.66 | 1.58 |
| KIF2A | O00139 | 0.0021 | 0.34 | 1.26 |
| KIF3A | Q9Y496 | 0.0024 | 1.45 | 2.73 |
| KIF4A | O95239 | 0.0026 | 0.36 | 1.28 |
| KIF7 | Q2M1P5 | 0.0028 | 0.6 | 1.51 |
| KLF13 | Q9Y2Y9 | 0.0015 | 0.98 | 1.98 |
| KLRG2 | A4D1S0 | 0.0022 | 0.76 | 1.7 |
| KPNA6 | O60684 | 2.40E-04 | 0.6 | 1.51 |
| KRT8 | P05787 | 0.013 | 0.66 | 1.58 |
| KRTCAP2 | Q8N6L1 | 0.0013 | 1.06 | 2.08 |
| LARP6 | Q9BRS8 | 1.40E-05 | 0.97 | 1.96 |
| LASP1 | Q14847 | 0.0063 | 1.09 | 2.14 |
| LATS2 | Q9NRM7 | 0.0019 | 0.64 | 1.56 |
| LDB2 | O43679 | 4.00E-04 | 1.14 | 2.21 |
| LENG8 | Q96PV6 | 0.0064 | 0.64 | 1.56 |
| LIMCH1 | Q9UPQ0 | 0.034 | 0.52 | 1.44 |
| LIMD1 | Q9UGP4 | 0.0073 | 0.29 | 1.23 |
| LIMK2 | P53671 | 4.00E-04 | 0.95 | 1.93 |
| LINGO1 | Q96FE5 | 1.10E-04 | 1.41 | 2.65 |

| | | | | |
|---|---|---|---|---|
| LMTK2 | Q8IWU2 | 0.011 | 0.38 | 1.3 |
| LRCH3 | Q96II8 | 0.0031 | 0.39 | 1.31 |
| LRP1 | Q07954 | 5.80E-04 | 1.64 | 3.12 |
| LRP4 | O75096 | 0.0053 | 0.32 | 1.24 |
| LRRC1 | Q9BTT6 | 2.00E-04 | 1.08 | 2.11 |
| LRRC40 | Q9H9A6 | 0.0043 | 0.3 | 1.23 |
| LRRC41 | Q15345 | 1.50E-04 | 0.68 | 1.6 |
| LRRC8A | Q8IWT6 | 0.0072 | 0.49 | 1.4 |
| LRRFIP2 | Q9Y608 | 1.50E-04 | 0.43 | 1.34 |
| LRRTM1 | Q86UE6 | 0.048 | 0.68 | 1.6 |
| LSM14B | Q9BX40 | 0.0096 | 0.4 | 1.32 |
| LUZP1 | Q86V48 | 3.10E-04 | 0.53 | 1.44 |
| LYAR | Q9NX58 | 8.50E-05 | 0.94 | 1.91 |
| LZTS1 | Q9Y250 | 0.0052 | 0.53 | 1.44 |
| LZTS3 | O60299 | 7.60E-05 | 0.77 | 1.71 |
| MAP2K4 | P45985 | 0.004 | 1.09 | 2.12 |
| MAP3K3 | Q99759 | 7.20E-05 | 0.74 | 1.67 |
| MAP4K3 | Q8IVH8 | 0.001 | 1.4 | 2.63 |
| MAP4K5 | Q9Y4K4 | 0.03 | 0.93 | 1.91 |
| MAP7D1 | Q3KQU3 | 0.0014 | 0.44 | 1.36 |
| MAPK6 | Q16659 | 0.027 | 1.11 | 2.16 |
| MAPKAPK5 | Q8IW41 | 9.80E-04 | 0.41 | 1.33 |
| MARCKS | P29966 | 4.50E-05 | 0.69 | 1.61 |
| MARK1 | Q9P0L2 | 0.0027 | 0.36 | 1.28 |
| MARK2 | Q7KZI7 | 0.011 | 0.39 | 1.31 |
| MARK3 | P27448 | 0.0029 | 0.42 | 1.34 |
| MARK4 | Q96L34 | 0.045 | 0.37 | 1.29 |
| MCC | P23508 | 7.30E-04 | 0.4 | 1.32 |
| MCM9 | Q9NXL9 | 0.004 | 0.45 | 1.36 |
| MDH1 | P40925 | 0.039 | 0.28 | 1.21 |
| MEN1 | O00255 | 5.70E-05 | 0.77 | 1.7 |
| METTL3 | Q86U44 | 2.00E-04 | 1.26 | 2.39 |
| MEX3D | Q86XN8 | 2.30E-04 | 1.54 | 2.91 |
| MFF | Q9GZY8 | 9.20E-04 | 0.48 | 1.39 |
| MGMT | P16455 | 7.80E-07 | 1.29 | 2.45 |
| MIB2 | Q96AX9 | 0.043 | 0.39 | 1.31 |
| MICALL1 | Q8N3F8 | 4.30E-05 | 1.05 | 2.07 |
| MID1 | O15344 | 2.00E-04 | 0.64 | 1.56 |
| MIIP | Q5JXC2 | 0.0057 | 0.38 | 1.31 |
| MKNK1 | Q9BUB5 | 5.50E-05 | 0.99 | 1.99 |
| MLLT10 | P55197 | 0.014 | 0.26 | 1.2 |

| | | | | |
|---|---|---|---|---|
| MLTK_HUMAN | NA | 6.50E-04 | 0.39 | 1.31 |
| MOCOS | Q96EN8 | 0.024 | 0.44 | 1.35 |
| MPG | P29372 | 0.0038 | 0.88 | 1.84 |
| MPLKIP | Q8TAP9 | 0.0037 | 0.52 | 1.44 |
| MSI1 | O43347 | 0.05 | 0.3 | 1.23 |
| MSN | P26038 | 8.20E-04 | 0.91 | 1.88 |
| MTBP | Q96DY7 | 0.0061 | 0.38 | 1.3 |
| MTDH | Q86UE4 | 0.037 | 0.44 | 1.36 |
| MTM1 | Q13496 | 9.80E-04 | 0.59 | 1.5 |
| MTMR12 | Q9C0I1 | 0.04 | 1.51 | 2.84 |
| MTMR2 | Q13614 | 0.033 | 0.31 | 1.24 |
| MUTYH | Q9UIF7 | 6.00E-05 | 1.29 | 2.44 |
| MYCBP2 | O75592 | 0.0017 | 0.62 | 1.54 |
| MYL12A | P19105 | 0.0018 | 1.35 | 2.56 |
| MYL5 | Q02045 | 0.006 | 1.09 | 2.12 |
| MYO10 | Q9HD67 | 0.016 | 0.45 | 1.37 |
| MYO19 | Q96H55 | 0.0032 | 0.52 | 1.44 |
| MZT2B | Q6NZ67 | 0.013 | 0.43 | 1.35 |
| N4BP1 | O75113 | 0.01 | 0.29 | 1.22 |
| NAB1 | Q13506 | 5.30E-04 | 0.85 | 1.81 |
| NAB2 | Q15742 | 0.0035 | 0.54 | 1.46 |
| NAP1L4 | Q99733 | 0.025 | 0.36 | 1.28 |
| NAV2 | Q8IVL1 | 0.037 | 0.27 | 1.2 |
| NBN | O60934 | 6.60E-04 | 0.76 | 1.7 |
| NCBP2 | P52298 | 0.0014 | 0.51 | 1.42 |
| NCK1 | P16333 | 0.0065 | 0.57 | 1.48 |
| NCKAP5L | Q9HCH0 | 0.0023 | 0.54 | 1.45 |
| NCOA3 | Q9Y6Q9 | 0.027 | 0.28 | 1.21 |
| NDE1 | Q9NXR1 | 0.0012 | 0.35 | 1.27 |
| NECTIN2 | Q92692 | 0.017 | 0.27 | 1.21 |
| NEK3 | P51956 | 0.013 | 0.52 | 1.43 |
| NELFA | Q9H3P2 | 0.0033 | 0.48 | 1.4 |
| NEPRO | Q6NW34 | 0.023 | 0.3 | 1.23 |
| NF1 | P21359 | 8.80E-04 | 0.57 | 1.48 |
| NFATC1 | O95644 | 8.70E-07 | 1.58 | 2.98 |
| NIPBL | Q6KC79 | 0.015 | 0.33 | 1.26 |
| NKAP | Q8N5F7 | 0.049 | 0.29 | 1.22 |
| NKIRAS2 | Q9NYR9 | 0.0025 | 1.39 | 2.63 |
| NME2 | P22392 | 0.014 | 0.31 | 1.24 |
| NOP16 | Q9Y3C1 | 0.0075 | 0.29 | 1.22 |
| NPAT | Q14207 | 0.002 | 0.43 | 1.35 |

| | | | | |
|---|---|---|---|---|
| NR3C1 | P04150 | 0.0037 | 0.98 | 1.97 |
| NRDC | O43847 | 0.0022 | 0.9 | 1.86 |
| NUMA1 | Q14980 | 3.90E-04 | 0.41 | 1.33 |
| NUP133 | Q8WUM0 | 0.017 | 0.68 | 1.6 |
| NUP153 | P49790 | 0.0087 | 0.28 | 1.21 |
| NUP35 | Q8NFH5 | 0.011 | 0.29 | 1.22 |
| NUP50 | Q9UKX7 | 1.40E-05 | 0.81 | 1.76 |
| NUSAP1 | Q9BXS6 | 0.0014 | 1.01 | 2.01 |
| OPLAH | O14841 | 0.0048 | 0.43 | 1.35 |
| OPTN | Q96CV9 | 0.035 | 0.35 | 1.27 |
| OSBPL10 | Q9BXB5 | 0.045 | 0.27 | 1.21 |
| OTUD4 | Q01804 | 0.0073 | 0.38 | 1.3 |
| PA2G4 | Q9UQ80 | 0.0022 | 0.42 | 1.33 |
| PACRGL | Q8N7B6 | 4.00E-05 | 1.07 | 2.11 |
| PACS2 | Q86VP3 | 0.024 | 0.39 | 1.31 |
| PAK6 | Q9NQU5 | 0.0034 | 0.68 | 1.6 |
| PAN3 | Q58A45 | 1.40E-04 | 1.3 | 2.46 |
| PANK1 | Q8TE04 | 1.30E-06 | 1.49 | 2.8 |
| PAPD7 | Q5XG87 | 0.0057 | 0.78 | 1.72 |
| PAPPA | Q13219 | 0.034 | 0.34 | 1.27 |
| PATL1 | Q86TB9 | 0.0046 | 0.45 | 1.36 |
| PATZ1 | Q9HBE1 | 4.20E-05 | 1.65 | 3.13 |
| PAWR | Q96IZ0 | 6.20E-04 | 1.15 | 2.22 |
| PBX2 | P40425 | 0.022 | 0.26 | 1.2 |
| PCIF1 | Q9H4Z3 | 0.0034 | 0.3 | 1.23 |
| PCNP | Q8WW12 | 0.032 | 0.33 | 1.26 |
| PDCD4 | Q53EL6 | 2.20E-06 | 1.19 | 2.28 |
| PDLIM1 | O00151 | 0.0025 | 0.65 | 1.57 |
| PDLIM4 | P50479 | 1.60E-05 | 0.84 | 1.79 |
| PDLIM5 | Q96HC4 | 0.001 | 0.84 | 1.79 |
| PDS5A | Q29RF7 | 0.0029 | 0.66 | 1.58 |
| PEAK1 | Q9H792 | 0.0037 | 0.4 | 1.32 |
| PELO | Q9BRX2 | 3.10E-04 | 0.89 | 1.85 |
| PERQ2_HUMAN | NA | 0.0036 | 0.86 | 1.82 |
| PFKFB3 | Q16875 | 0.019 | 0.45 | 1.36 |
| PHACTR2 | O75167 | 0.0022 | 0.4 | 1.32 |
| PHF13 | Q86YI8 | 0.033 | 0.63 | 1.55 |
| PHF3 | Q92576 | 0.006 | 0.37 | 1.29 |
| PHLDB1 | Q86UU1 | 7.50E-04 | 0.73 | 1.65 |
| PI4KA | P42356 | 2.10E-04 | 0.77 | 1.7 |
| PI4KB | Q9UBF8 | 0.0069 | 0.54 | 1.45 |

| | | | | |
|---|---|---|---|---|
| PIKFYVE | Q9Y2I7 | 0.012 | 0.58 | 1.49 |
| PITPNC1 | Q9UKF7 | 1.80E-04 | 1 | 2.01 |
| PKP4 | Q99569 | 0.0092 | 0.3 | 1.23 |
| PLAA | Q9Y263 | 0.0096 | 0.29 | 1.22 |
| PLCH1 | Q4KWH8 | 0.015 | 0.34 | 1.27 |
| PLCL2 | Q9UPR0 | 7.00E-05 | 1.07 | 2.09 |
| PLEKHA1 | Q9HB21 | 0.003 | 0.66 | 1.58 |
| PLEKHG2 | Q9H7P9 | 0.039 | 0.39 | 1.31 |
| PLXDC2 | Q6UX71 | 5.80E-07 | 1.22 | 2.33 |
| POLM | Q9NP87 | 1.40E-04 | 0.6 | 1.52 |
| POLR2M | P0CAP2 | 0.0064 | 0.58 | 1.5 |
| POU2F1 | P14859 | 0.0027 | 0.35 | 1.27 |
| PPFIBP2 | Q8ND30 | 1.10E-04 | 0.48 | 1.4 |
| PPM1H | Q9ULR3 | 0.0022 | 0.52 | 1.44 |
| PPP1CB | P62140 | 7.60E-04 | 0.44 | 1.36 |
| PPP1R12A | O14974 | 0.0017 | 0.47 | 1.39 |
| PPP1R12C | Q9BZL4 | 1.20E-04 | 0.92 | 1.9 |
| PPP1R13B | Q96KQ4 | 0.0018 | 0.41 | 1.33 |
| PPP1R13L | Q8WUF5 | 0.002 | 0.94 | 1.92 |
| PPP1R1A | Q13522 | 0.017 | 0.33 | 1.26 |
| PPP1R26 | Q5T8A7 | 0.0078 | 0.29 | 1.22 |
| PPP1R35 | Q8TAP8 | 7.00E-05 | 1.45 | 2.73 |
| PPP4R3B | Q5MIZ7 | 0.011 | 0.34 | 1.27 |
| PPP6R2 | O75170 | 0.0063 | 0.27 | 1.21 |
| PRC1 | O43663 | 0.0028 | 1.06 | 2.08 |
| PRKAA1 | Q13131 | 0.0019 | 0.51 | 1.43 |
| PRKACB | P22694 | 0.0011 | 1.05 | 2.07 |
| PRKD3 | O94806 | 0.0018 | 0.35 | 1.27 |
| PRKDC | P78527 | 0.035 | 0.28 | 1.21 |
| PRKRA | O75569 | 0.021 | 0.83 | 1.78 |
| PRPF40B | Q6NWY9 | 0.0035 | 0.3 | 1.23 |
| PRR36 | Q9H6K5 | 1.30E-06 | 1.17 | 2.25 |
| PSMA3 | P25788 | 0.041 | 0.3 | 1.23 |
| PSMD11 | O00231 | 2.80E-04 | 1.4 | 2.63 |
| PTMS | P20962 | 4.10E-06 | 0.83 | 1.77 |
| PTPN14 | Q15678 | 0.0032 | 0.36 | 1.28 |
| PTPN21 | Q16825 | 0.033 | 0.34 | 1.26 |
| PUM1 | Q14671 | 5.60E-05 | 0.71 | 1.64 |
| PUM2 | Q8TB72 | 0.0013 | 0.51 | 1.43 |
| PWWP2B | Q6NUJ5 | 0.012 | 0.5 | 1.42 |
| PYGO2 | Q9BRQ0 | 1.10E-04 | 0.73 | 1.66 |

| | | | | |
|---|---|---|---|---|
| RAB11A | P62491 | 0.0034 | 0.84 | 1.79 |
| RAB11FIP1 | Q6WKZ4 | 1.00E-04 | 0.54 | 1.45 |
| RAB11FIP5 | Q9BXF6 | 1.50E-05 | 1.06 | 2.08 |
| RAB3IP | Q96QF0 | 0.0058 | 0.55 | 1.47 |
| RAB7A | P51149 | 6.90E-04 | 1.03 | 2.04 |
| RABEP1 | Q15276 | 0.0073 | 0.54 | 1.45 |
| RABGAP1L | Q5R372 | 0.036 | 0.3 | 1.23 |
| RACGAP1 | Q9H0H5 | 9.80E-05 | 0.59 | 1.51 |
| RAP1GAP | P47736 | 3.10E-04 | 0.83 | 1.77 |
| RASGRP2 | Q7LDG7 | 6.30E-04 | 1.46 | 2.76 |
| RASSF7 | Q02833 | 0.013 | 0.35 | 1.28 |
| RASSF8 | Q8NHQ8 | 0.014 | 0.44 | 1.36 |
| RBL1 | P28749 | 0.0014 | 0.65 | 1.57 |
| RBM26 | Q5T8P6 | 0.045 | 0.27 | 1.21 |
| RBM27 | Q9P2N5 | 2.10E-05 | 1.35 | 2.56 |
| RBM3 | P98179 | 1.00E-06 | 1.5 | 2.82 |
| RBM34 | P42696 | 3.70E-04 | 1.33 | 2.51 |
| RBM7 | Q9Y580 | 0.015 | 0.55 | 1.46 |
| RBPMS | Q93062 | 0.017 | 0.42 | 1.34 |
| RBSN | Q9H1K0 | 0.041 | 0.42 | 1.34 |
| RCC1 | P18754 | 0.028 | 0.48 | 1.4 |
| RCC2 | Q9P258 | 0.0042 | 0.42 | 1.34 |
| RELL1 | Q8IUW5 | 0.021 | 0.41 | 1.33 |
| RGPD5 | Q99666 | 0.017 | 0.31 | 1.24 |
| RGS12 | O14924 | 4.60E-04 | 0.71 | 1.63 |
| RHPN1 | Q8TCX5 | 0.0038 | 0.65 | 1.57 |
| RHPN2 | Q8IUC4 | 0.014 | 0.38 | 1.3 |
| RICTOR | Q6R327 | 0.017 | 0.39 | 1.31 |
| RIPK2 | O43353 | 0.01 | 0.58 | 1.49 |
| RITA1 | Q96K30 | 0.002 | 0.51 | 1.43 |
| RMDN3 | Q96TC7 | 0.01 | 0.37 | 1.29 |
| RNF123 | Q5XPI4 | 2.60E-07 | 1.93 | 3.82 |
| RNF219 | Q5W0B1 | 0.0045 | 0.32 | 1.25 |
| RNF25 | Q96BH1 | 0.0023 | 2 | 3.99 |
| RNF8 | O76064 | 0.025 | 0.62 | 1.53 |
| RPA2 | P15927 | 0.0019 | 0.49 | 1.4 |
| RPAP1 | Q9BWH6 | 0.0085 | 0.59 | 1.51 |
| RPL18 | Q07020 | 2.00E-04 | 0.89 | 1.85 |
| RPL27A | P46776 | 0.048 | 0.88 | 1.84 |
| RPL34 | P49207 | 0.04 | 1.05 | 2.06 |
| RPS28 | P62857 | 0.01 | 0.75 | 1.69 |

| | | | | |
|---|---|---|---|---|
| RPS6KB1 | P23443 | 6.20E-05 | 1.17 | 2.24 |
| RRAGC | Q9HB90 | 0.0022 | 0.46 | 1.38 |
| RRM2B | Q7LG56 | 0.016 | 0.87 | 1.83 |
| RTEL1 | Q9NZ71 | 5.00E-04 | 0.75 | 1.68 |
| RTKN | Q9BST9 | 0.0045 | 0.42 | 1.34 |
| SAAL1 | Q96ER3 | 0.02 | 0.36 | 1.28 |
| SAV1 | Q9H4B6 | 7.70E-04 | 0.68 | 1.6 |
| SBF2 | Q86WG5 | 8.80E-05 | 0.69 | 1.62 |
| SCAMP3 | O14828 | 0.0052 | 0.54 | 1.46 |
| SDCBP | O00560 | 0.032 | 0.38 | 1.3 |
| SEC24B | O95487 | 0.0064 | 0.55 | 1.46 |
| SELENOS | Q9BQE4 | 0.014 | 0.45 | 1.37 |
| SERINC1 | Q9NRX5 | 0.017 | 0.28 | 1.21 |
| SETD1B | Q9UPS6 | 0.0078 | 0.75 | 1.68 |
| SETMAR | Q53H47 | 0.004 | 0.58 | 1.49 |
| SGO1 | Q5FBB7 | 8.00E-05 | 0.9 | 1.87 |
| SGPL1 | O95470 | 0.024 | 0.38 | 1.3 |
| SGTA | O43765 | 0.0059 | 0.74 | 1.67 |
| SH2D1B | O14796 | 0.0088 | 0.47 | 1.39 |
| SH3BP2 | P78314 | 7.60E-04 | 1.53 | 2.88 |
| SHB | Q15464 | 0.033 | 0.33 | 1.26 |
| SHMT2 | P34897 | 2.40E-04 | 0.73 | 1.66 |
| SIK2 | Q9H0K1 | 0.009 | 0.47 | 1.39 |
| SKI | P12755 | 3.80E-04 | 0.65 | 1.57 |
| SKIV2L2 | P42285 | 0.0044 | 0.91 | 1.88 |
| SLAIN2 | Q9P270 | 0.0032 | 0.59 | 1.5 |
| SLC12A7 | Q9Y666 | 5.50E-04 | 0.69 | 1.62 |
| SLC2A6 | Q9UGQ3 | 0.0032 | 0.34 | 1.26 |
| SLC35F2 | Q8IXU6 | 0.0094 | 0.28 | 1.21 |
| SLC4A7 | Q9Y6M7 | 0.018 | 0.42 | 1.34 |
| SLC6A15 | Q9H2J7 | 4.00E-06 | 1.01 | 2.01 |
| SLC7A2 | P52569 | 0.037 | 1.28 | 2.43 |
| SLC9A3R2 | Q15599 | 0.0074 | 0.59 | 1.51 |
| SLCO4C1 | Q6ZQN7 | 7.70E-04 | 0.47 | 1.39 |
| SMC1A | Q14683 | 0.0034 | 0.96 | 1.94 |
| SMC5 | Q8IY18 | 8.50E-05 | 0.68 | 1.61 |
| SMCHD1 | A6NHR9 | 0.037 | 0.29 | 1.22 |
| SMCR8 | Q8TEV9 | 0.0053 | 0.36 | 1.28 |
| SMG1 | Q96Q15 | 0.0055 | 0.42 | 1.34 |
| SMG9 | Q9H0W8 | 0.015 | 0.63 | 1.55 |
| SNRK | Q9NRH2 | 0.0043 | 0.43 | 1.35 |

| | | | | |
|---|---|---|---|---|
| SNRPA | P09012 | 7.10E-05 | 0.65 | 1.57 |
| SNURF | Q9Y675 | 0.013 | 0.33 | 1.26 |
| SNX18 | Q96RF0 | 0.028 | 0.66 | 1.58 |
| SNX2 | O60749 | 0.0033 | 0.37 | 1.29 |
| SNX4 | O95219 | 0.0017 | 0.34 | 1.27 |
| SORBS1 | Q9BX66 | 0.008 | 0.28 | 1.21 |
| SORBS2 | O94875 | 2.70E-04 | 0.53 | 1.44 |
| SOWAHC | Q53LP3 | 0.028 | 0.26 | 1.2 |
| SOX10 | P56693 | 0.0012 | 1.09 | 2.14 |
| SP110 | Q9HB58 | 8.20E-04 | 0.49 | 1.4 |
| SP2 | Q02086 | 0.0019 | 0.85 | 1.8 |
| SPATS2L | Q9NUQ6 | 0.015 | 0.28 | 1.21 |
| SPICE1 | Q8N0Z3 | 0.026 | 0.41 | 1.33 |
| SPN | P16150 | 0.0062 | 1.29 | 2.44 |
| SRA1 | Q9HD15 | 3.80E-04 | 0.71 | 1.64 |
| SRC | P12931 | 0.01 | 0.48 | 1.39 |
| SRGAP1 | Q7Z6B7 | 0.0016 | 0.7 | 1.62 |
| SRSF3 | P84103 | 0.034 | 0.56 | 1.48 |
| SSH3 | Q8TE77 | 0.0032 | 0.29 | 1.22 |
| SSR3 | Q9UNL2 | 0.0066 | 0.47 | 1.38 |
| STAM | Q92783 | 0.006 | 0.43 | 1.35 |
| STAP2 | Q9UGK3 | 0.0022 | 0.59 | 1.5 |
| STAT3 | P40763 | 0.0038 | 0.55 | 1.47 |
| STBD1 | O95210 | 0.008 | 1.05 | 2.07 |
| STEAP3 | Q658P3 | 0.043 | 0.47 | 1.39 |
| STIM1 | Q13586 | 0.011 | 0.66 | 1.58 |
| STK10 | O94804 | 0.0074 | 0.65 | 1.57 |
| STK11 | Q15831 | 0.0079 | 0.31 | 1.24 |
| STRAP | Q9Y3F4 | 0.049 | 0.55 | 1.47 |
| STX17 | P56962 | 0.0034 | 0.39 | 1.31 |
| SUB1 | P53999 | 0.003 | 0.66 | 1.58 |
| SUGP1 | Q8IWZ8 | 0.0071 | 0.33 | 1.25 |
| SUGT1 | Q9Y2Z0 | 5.30E-04 | 0.69 | 1.61 |
| SUN1 | O94901 | 0.0054 | 0.63 | 1.55 |
| SUPT20H | Q8NEM7 | 0.037 | 0.36 | 1.29 |
| SUPT6H | Q7KZ85 | 0.017 | 0.27 | 1.21 |
| SVIL | O95425 | 4.10E-04 | 0.82 | 1.77 |
| SYMPK | Q92797 | 0.045 | 0.32 | 1.25 |
| SYN3 | O14994 | 0.023 | 0.38 | 1.31 |
| SYNRG | Q9UMZ2 | 0.021 | 0.29 | 1.22 |
| TAB1 | Q15750 | 0.04 | 0.38 | 1.3 |

| | | | | |
|---|---|---|---|---|
| TANC2 | Q9HCD6 | 0.017 | 0.3 | 1.23 |
| TBC1D13 | Q9NVG8 | 1.60E-05 | 1.28 | 2.43 |
| TBC1D22B | Q9NU19 | 0.034 | 0.5 | 1.41 |
| TBC1D5 | Q92609 | 3.60E-04 | 0.53 | 1.45 |
| TEAD3 | Q99594 | 4.00E-04 | 0.58 | 1.49 |
| TECPR2 | O15040 | 0.0024 | 0.8 | 1.74 |
| TERF2 | Q15554 | 0.014 | 0.4 | 1.32 |
| TGIF1 | Q15583 | 1.50E-04 | 0.8 | 1.75 |
| TICRR | Q7Z2Z1 | 5.10E-04 | 0.46 | 1.38 |
| TK1 | P04183 | 0.024 | 0.29 | 1.22 |
| TMCC1 | O94876 | 0.05 | 1.24 | 2.36 |
| TMEM237 | Q96Q45 | 0.0068 | 0.75 | 1.68 |
| TMEM51 | Q9NW97 | 0.024 | 0.3 | 1.23 |
| TMEM63B | Q5T3F8 | 0.0068 | 0.33 | 1.26 |
| TMEM9 | Q9P0T7 | 0.013 | 0.42 | 1.33 |
| TNFRSF11A | Q9Y6Q6 | 0.031 | 0.31 | 1.24 |
| TNS1 | Q9HBL0 | 0.0029 | 0.49 | 1.4 |
| TOP3A | Q13472 | 2.60E-04 | 1.42 | 2.67 |
| TP53BP2 | Q13625 | 0.014 | 0.33 | 1.26 |
| TPD52L2 | O43399 | 0.0051 | 0.42 | 1.33 |
| TPX2 | Q9ULW0 | 1.20E-04 | 1.25 | 2.37 |
| TRAF7 | Q6Q0C0 | 0.021 | 0.37 | 1.29 |
| TRIM22 | Q8IYM9 | 0.011 | 0.48 | 1.39 |
| TRIM56 | Q9BRZ2 | 0.0034 | 0.57 | 1.48 |
| TRIM71 | Q2Q1W2 | 0.02 | 0.5 | 1.41 |
| TRIO | O75962 | 3.80E-04 | 0.74 | 1.67 |
| TRIP11 | Q15643 | 1.70E-06 | 1.69 | 3.23 |
| TRIP6 | Q15654 | 0.0032 | 0.98 | 1.97 |
| TRMT44 | Q8IYL2 | 0.0036 | 0.51 | 1.42 |
| TRPM7 | Q96QT4 | 0.0053 | 0.31 | 1.24 |
| TTC28 | Q96AY4 | 0.028 | 0.33 | 1.25 |
| TTC39B | Q5VTQ0 | 0.026 | 1.01 | 2.01 |
| TTK | P33981 | 0.0021 | 0.62 | 1.54 |
| TXNRD3 | Q86VQ6 | 2.50E-05 | 1.5 | 2.82 |
| UBE2J1 | Q9Y385 | 0.0022 | 0.75 | 1.69 |
| UBE3A | Q05086 | 0.0035 | 0.4 | 1.32 |
| UBN1 | Q9NPG3 | 0.021 | 0.39 | 1.31 |
| UBQLN4 | Q9NRR5 | 0.0078 | 0.34 | 1.27 |
| UBXN1 | Q04323 | 5.50E-06 | 0.87 | 1.83 |
| UBXN2A | P68543 | 0.038 | 0.3 | 1.23 |
| UBXN2B | Q14CS0 | 0.016 | 0.42 | 1.34 |

| | | | | |
|---|---|---|---|---|
| UBXN7 | O94888 | 4.10E-04 | 1.02 | 2.03 |
| UCK2 | Q9BZX2 | 0.043 | 0.59 | 1.5 |
| UFD1 | Q92890 | 6.20E-05 | 0.64 | 1.55 |
| UGP2 | Q16851 | 0.025 | 0.28 | 1.22 |
| UHRF1 | Q96T88 | 0.0019 | 0.46 | 1.38 |
| UNC45A | Q9H3U1 | 0.0038 | 0.66 | 1.58 |
| UNG | P13051 | 0.018 | 0.43 | 1.35 |
| URI1 | O94763 | 0.044 | 0.39 | 1.31 |
| USP22 | Q9UPT9 | 0.016 | 0.38 | 1.3 |
| USP24 | Q9UPU5 | 0.029 | 0.36 | 1.29 |
| USP32 | Q8NFA0 | 0.0033 | 0.4 | 1.32 |
| USP5 | P45974 | 0.033 | 0.33 | 1.26 |
| USP6NL | Q92738 | 0.0032 | 0.74 | 1.67 |
| UTY | O14607 | 0.013 | 0.31 | 1.24 |
| VASH1 | Q7L8A9 | 0.022 | 0.41 | 1.33 |
| VIM | P08670 | 0.045 | 0.27 | 1.21 |
| VIPAS39 | Q9H9C1 | 0.018 | 0.7 | 1.63 |
| VPRBP_HUMAN | NA | 1.70E-04 | 0.59 | 1.51 |
| WDR13 | Q9H1Z4 | 4.10E-04 | 0.96 | 1.95 |
| WDR20 | Q8TBZ3 | 0.0067 | 0.96 | 1.94 |
| WDR37 | Q9Y2I8 | 0.0015 | 1.13 | 2.19 |
| WEE1 | P30291 | 0.011 | 0.38 | 1.3 |
| WIPF1 | O43516 | 0.033 | 0.81 | 1.75 |
| WIPF2 | Q8TF74 | 0.0024 | 0.6 | 1.52 |
| WNK2 | Q9Y3S1 | 0.0052 | 0.38 | 1.3 |
| WNK3 | Q9BYP7 | 0.026 | 0.26 | 1.2 |
| WRNIP1 | Q96S55 | 0.0024 | 0.45 | 1.36 |
| WTIP | A6NIX2 | 0.0098 | 0.8 | 1.74 |
| WWC1 | Q8IX03 | 0.0075 | 0.3 | 1.23 |
| WWP2 | O00308 | 4.70E-06 | 0.98 | 1.97 |
| YAP1 | P46937 | 0.0092 | 0.35 | 1.27 |
| YTHDF3 | Q7Z739 | 0.0024 | 0.59 | 1.51 |
| ZCCHC11 | Q5TAX3 | 0.0049 | 0.55 | 1.47 |
| ZCCHC17 | Q9NP64 | 0.0076 | 0.53 | 1.44 |
| ZDHHC5 | Q9C0B5 | 0.013 | 0.49 | 1.4 |
| ZDHHC8 | Q9ULC8 | 0.017 | 0.33 | 1.26 |
| ZFP36 | P26651 | 0.028 | 0.45 | 1.36 |
| ZFP36L1 | Q07352 | 0.0086 | 0.87 | 1.83 |
| ZFP36L2 | P47974 | 0.046 | 0.27 | 1.21 |
| ZHX2 | Q9Y6X8 | 0.0024 | 0.9 | 1.86 |
| ZIC3 | O60481 | 0.0016 | 0.46 | 1.38 |

| | | | | |
|---|---|---|---|---|
| ZNF185 | O15231 | 0.0088 | 0.63 | 1.55 |
| ZNF195 | O14628 | 1.00E-05 | 1.01 | 2.01 |
| ZNF3 | P17036 | 0.0018 | 0.58 | 1.49 |
| ZNF34 | Q8IZ26 | 0.041 | 0.38 | 1.3 |
| ZNF391 | Q9UJN7 | 0.023 | 0.51 | 1.43 |
| ZNF513 | Q8N8E2 | 0.0013 | 0.6 | 1.51 |
| ZNF516 | Q92618 | 1.80E-06 | 0.86 | 1.81 |
| ZNF521 | Q96K83 | 1.40E-04 | 0.6 | 1.52 |
| ZNF608 | Q9ULD9 | 5.60E-05 | 0.81 | 1.75 |
| ZNRF2 | Q8NHG8 | 0.012 | 0.35 | 1.28 |
| ZZEF1 | O43149 | 0.0038 | 0.53 | 1.44 |

| **Supplementary Table 3B:**<br>***KDM6A*KO downregulated phospho-protein expression by mass spectrometry**<br>[pVal = p-value; logFC = log of the fold change; FC = fold change in expression] | | | | |
|---|---|---|---|---|
| **Protein** | **Accession** | **pVal** | **logFC** | **FC** |
| AAAS | Q9NRG9 | 0.0021 | -0.87 | -1.83 |
| ABCF1 | Q8NE71 | 0.012 | -0.28 | -1.21 |
| ACACA | Q13085 | 0.04 | -0.29 | -1.22 |
| ACTL6A | O96019 | 9.80E-04 | -0.93 | -1.9 |
| ADD2 | P35612 | 0.0022 | -0.33 | -1.25 |
| ADGRL3 | Q9HAR2 | 9.90E-05 | -1.4 | -2.65 |
| AGBL2 | Q5U5Z8 | 0.003 | -1.03 | -2.04 |
| AGO1 | Q9UL18 | 0.039 | -0.41 | -1.33 |
| AIFM1 | O95831 | 0.0073 | -0.39 | -1.31 |
| ALDH18A1 | P54886 | 6.90E-04 | -0.79 | -1.73 |
| ALDH4A1 | P30038 | 0.018 | -0.44 | -1.35 |
| ALPL | P05186 | 1.10E-05 | -1.05 | -2.07 |
| ANAPC4 | Q9UJX5 | 0.027 | -0.27 | -1.2 |
| ANKRD13D | Q6ZTN6 | 0.034 | -0.32 | -1.25 |
| ANXA2 | P07355 | 0.02 | -0.45 | -1.36 |
| APBA2 | Q99767 | 0.0056 | -0.75 | -1.68 |
| APMAP | Q9HDC9 | 0.048 | -0.26 | -1.2 |
| APPL1 | Q9UKG1 | 0.0018 | -0.57 | -1.48 |
| ARID4A | P29374 | 0.01 | -0.35 | -1.28 |
| ARMC1 | Q9NVT9 | 0.0057 | -0.31 | -1.24 |
| ASTN1 | O14525 | 0.01 | -0.3 | -1.23 |
| ATAD3A | Q9NVI7 | 9.50E-04 | -0.37 | -1.29 |
| ATAD5 | Q96QE3 | 0.043 | -0.42 | -1.33 |
| ATHL1_HUMAN | NA | 0.013 | -0.39 | -1.31 |
| ATP13A3 | Q9H7F0 | 0.0014 | -0.35 | -1.27 |
| ATP1A1 | P05023 | 0.0044 | -0.44 | -1.35 |
| ATP2A2 | P16615 | 7.80E-06 | -1.04 | -2.05 |
| ATP2B1 | P20020 | 0.0036 | -0.32 | -1.25 |
| ATP2B4 | P23634 | 4.70E-04 | -0.57 | -1.49 |
| ATXN7L3B | Q96GX2 | 0.044 | -0.57 | -1.48 |
| AXIN2 | Q9Y2T1 | 0.0093 | -0.48 | -1.39 |
| AXL | P30530 | 0.027 | -0.33 | -1.26 |
| BARD1 | Q99728 | 0.0051 | -0.34 | -1.27 |
| BATF3 | Q9NR55 | 3.40E-05 | -0.72 | -1.65 |
| BAZ2A | Q9UIF9 | 0.021 | -0.28 | -1.22 |
| BBC3 | Q96PG8 | 0.0044 | -0.43 | -1.35 |
| BCAR1 | P56945 | 4.80E-05 | -0.69 | -1.61 |

| | | | | |
|---|---|---|---|---|
| BCCIP | Q9P287 | 0.0071 | -0.99 | -1.98 |
| BCKDHA | P12694 | 0.025 | -0.42 | -1.34 |
| BCL2L12 | Q9HB09 | 0.0032 | -0.4 | -1.32 |
| BDP1 | A6H8Y1 | 0.0095 | -0.32 | -1.25 |
| BNC1 | Q01954 | 0.023 | -0.75 | -1.68 |
| BRPF3 | Q9ULD4 | 0.031 | -0.39 | -1.31 |
| BTAF1 | O14981 | 0.0083 | -0.75 | -1.68 |
| BYSL | Q13895 | 0.0053 | -0.29 | -1.22 |
| C11orf84 | Q9BUA3 | 0.0075 | -0.31 | -1.24 |
| C17orf75 | Q9HAS0 | 0.023 | -0.3 | -1.24 |
| C19orf43 | Q9BQ61 | 0.019 | -0.29 | -1.22 |
| C19orf47 | Q8N9M1 | 0.0044 | -0.37 | -1.3 |
| C1orf52 | Q8N6N3 | 0.0016 | -0.37 | -1.29 |
| C7orf50 | Q9BRJ6 | 0.0013 | -0.33 | -1.26 |
| C9orf16 | Q9BUW7 | 0.0061 | -0.36 | -1.28 |
| CA14 | Q9ULX7 | 0.015 | -0.43 | -1.34 |
| CAAP1 | Q9H8G2 | 0.044 | -0.3 | -1.23 |
| CACNA1A | O00555 | 0.0016 | -0.68 | -1.6 |
| CADPS | Q9ULU8 | 0.027 | -0.27 | -1.21 |
| CASC3 | O15234 | 0.0035 | -0.36 | -1.28 |
| CAV1 | Q03135 | 2.00E-05 | -1.24 | -2.36 |
| CAV2 | P51636 | 1.80E-04 | -1.63 | -3.1 |
| CBARP | Q8N350 | 0.0029 | -0.31 | -1.24 |
| CBLL1 | Q75N03 | 0.0024 | -0.57 | -1.49 |
| CBR1 | P16152 | 0.0083 | -0.46 | -1.37 |
| CBX1 | P83916 | 0.0029 | -0.74 | -1.67 |
| CCDC137 | Q6PK04 | 0.0026 | -0.91 | -1.88 |
| CCDC177 | Q9NQR7 | 0.0038 | -0.92 | -1.89 |
| CCDC94 | Q9BW85 | 0.011 | -0.3 | -1.23 |
| CCNH | P51946 | 0.05 | -0.31 | -1.24 |
| CCT2 | P78371 | 0.012 | -0.5 | -1.41 |
| CCT4 | P50991 | 0.001 | -0.44 | -1.36 |
| CD44 | P16070 | 0.0014 | -0.69 | -1.61 |
| CDC25B | P30305 | 0.0011 | -0.52 | -1.44 |
| CDC26 | Q8NHZ8 | 0.0052 | -0.38 | -1.3 |
| CDCA7L | Q96GN5 | 0.0035 | -0.33 | -1.26 |
| CDK7 | P50613 | 5.00E-04 | -0.76 | -1.69 |
| CDS2 | O95674 | 0.0053 | -0.45 | -1.37 |
| CDYL | Q9Y232 | 0.0099 | -0.3 | -1.23 |
| CEBPZ | Q03701 | 3.10E-04 | -0.52 | -1.44 |
| CENPJ | Q9HC77 | 0.0081 | -0.6 | -1.51 |

| | | | | |
|---|---|---|---|---|
| CEP162 | Q5TB80 | 0.0046 | -0.45 | -1.37 |
| CEP85 | Q6P2H3 | 0.024 | -0.3 | -1.23 |
| CHFR | Q96EP1 | 0.014 | -0.33 | -1.25 |
| CHPF2 | Q9P2E5 | 8.70E-04 | -0.67 | -1.59 |
| CHTF18 | Q8WVB6 | 0.0015 | -0.92 | -1.89 |
| CJ012_HUMAN | NA | 0.0021 | -0.6 | -1.52 |
| CLCC1 | Q96S66 | 0.0025 | -0.39 | -1.31 |
| CLDN23 | Q96B33 | 0.013 | -0.27 | -1.21 |
| CLK1 | P49759 | 0.018 | -0.37 | -1.29 |
| CNBP | P62633 | 0.0073 | -0.38 | -1.3 |
| CNKSR2 | Q8WXI2 | 1.00E-04 | -0.54 | -1.46 |
| COPA | P53621 | 0.014 | -0.32 | -1.25 |
| COPS3 | Q9UNS2 | 0.0013 | -0.39 | -1.31 |
| CORO7 | P57737 | 0.0031 | -0.53 | -1.44 |
| CPNE7 | Q9UBL6 | 0.0089 | -0.32 | -1.25 |
| CPSF1 | Q10570 | 0.015 | -0.34 | -1.27 |
| CSDC2 | Q9Y534 | 0.023 | -0.47 | -1.38 |
| CSDE1 | O75534 | 6.80E-04 | -0.44 | -1.36 |
| CSRP1 | P21291 | 5.90E-04 | -0.64 | -1.56 |
| CTNNA2 | P26232 | 0.045 | -0.29 | -1.22 |
| CTPS2 | Q9NRF8 | 4.30E-04 | -1.21 | -2.32 |
| CTTN | Q14247 | 0.0022 | -0.53 | -1.44 |
| CWC22 | Q9HCG8 | 0.011 | -1.66 | -3.16 |
| CXADR | P78310 | 0.016 | -0.37 | -1.29 |
| CXXC1 | Q9P0U4 | 0.0085 | -0.57 | -1.48 |
| CYLD | Q9NQC7 | 0.0048 | -0.55 | -1.46 |
| DAAM2 | Q86T65 | 3.10E-04 | -0.55 | -1.46 |
| DAB1 | O75553 | 0.0064 | -0.49 | -1.4 |
| DAB2 | P98082 | 0.0062 | -0.32 | -1.25 |
| DAG1 | Q14118 | 0.0037 | -1.21 | -2.32 |
| DBF4 | Q9UBU7 | 0.0042 | -0.27 | -1.21 |
| DCUN1D3 | Q8IWE4 | 0.014 | -0.68 | -1.6 |
| DDX1 | Q92499 | 0.023 | -0.29 | -1.22 |
| DDX18 | Q9NVP1 | 0.027 | -0.42 | -1.33 |
| DDX21 | Q9NR30 | 0.0064 | -0.47 | -1.38 |
| DDX23 | Q9BUQ8 | 0.0065 | -0.56 | -1.47 |
| DDX24 | Q9GZR7 | 0.015 | -1.63 | -3.08 |
| DDX39B | Q13838 | 0.014 | -0.32 | -1.25 |
| DDX42 | Q86XP3 | 0.0013 | -0.39 | -1.31 |
| DDX54 | Q8TDD1 | 0.01 | -0.35 | -1.27 |
| DHX34 | Q14147 | 0.01 | -0.42 | -1.34 |

| | | | | |
|---|---|---|---|---|
| DHX9 | Q08211 | 0.0024 | -0.31 | -1.24 |
| DIDO1 | Q9BTC0 | 0.0025 | -0.3 | -1.23 |
| DKC1 | O60832 | 6.30E-04 | -0.46 | -1.38 |
| DLG1 | Q12959 | 0.0079 | -0.3 | -1.23 |
| DLG5 | Q8TDM6 | 0.028 | -0.27 | -1.21 |
| DNAJB6 | O75190 | 0.0021 | -0.79 | -1.73 |
| DNAJC17 | Q9NVM6 | 4.60E-05 | -0.83 | -1.78 |
| DNAJC6 | O75061 | 0.0023 | -0.38 | -1.3 |
| DNAJC7 | Q99615 | 0.0064 | -0.29 | -1.23 |
| DNAJC9 | Q8WXX5 | 0.035 | -0.74 | -1.67 |
| DNLZ | Q5SXM8 | 0.017 | -0.34 | -1.27 |
| DNMT3A | Q9Y6K1 | 5.90E-04 | -0.39 | -1.31 |
| DNMT3B | Q9UBC3 | 0.03 | -0.4 | -1.32 |
| DOCK2 | Q92608 | 0.026 | -0.34 | -1.27 |
| DOCK9 | Q9BZ29 | 3.10E-05 | -0.83 | -1.78 |
| DPOLZ_HUMAN | NA | 0.01 | -0.29 | -1.23 |
| DPPA2 | Q7Z7J5 | 4.50E-04 | -1.34 | -2.53 |
| DUT | P33316 | 0.0032 | -0.35 | -1.27 |
| E4F1 | Q66K89 | 0.0082 | -0.52 | -1.44 |
| EDAR | Q9UNE0 | 0.0016 | -0.72 | -1.65 |
| EDF1 | O60869 | 0.0042 | -1.4 | -2.64 |
| EEF1B2 | P24534 | 0.0056 | -0.42 | -1.34 |
| EEF1D | P29692 | 0.0016 | -0.33 | -1.26 |
| EEF2 | P13639 | 0.029 | -0.38 | -1.31 |
| EEF2K | O00418 | 0.035 | -0.27 | -1.21 |
| EFNB3 | Q15768 | 0.029 | -0.36 | -1.28 |
| EFTUD2 | Q15029 | 5.40E-05 | -0.58 | -1.49 |
| EHD1 | Q9H4M9 | 9.40E-06 | -1.05 | -2.07 |
| EHD4 | Q9H223 | 1.10E-04 | -0.59 | -1.5 |
| EIF2A | Q9BY44 | 0.0012 | -0.35 | -1.28 |
| EIF2AK1 | Q9BQI3 | 0.018 | -0.37 | -1.29 |
| EIF2AK3 | Q9NZJ5 | 0.0012 | -0.38 | -1.3 |
| EIF3B | P55884 | 0.013 | -0.34 | -1.27 |
| EIF3CL | B5ME19 | 0.005 | -0.35 | -1.28 |
| EIF3E | P60228 | 0.0062 | -0.27 | -1.21 |
| EIF3F | O00303 | 0.0062 | -0.56 | -1.47 |
| EIF3G | O75821 | 0.018 | -0.29 | -1.22 |
| EIF3H | O15372 | 0.0067 | -0.41 | -1.33 |
| EIF4E2 | O60573 | 0.018 | -0.82 | -1.77 |
| ELL | P55199 | 0.0012 | -0.39 | -1.31 |
| ELL2 | O00472 | 1.40E-04 | -0.57 | -1.48 |

| | | | | |
|---|---|---|---|---|
| ELOB | Q15370 | 0.01 | -0.47 | -1.38 |
| ENPP1 | P22413 | 0.043 | -0.64 | -1.56 |
| EPB41L1 | Q9H4G0 | 0.0025 | -0.66 | -1.58 |
| EPB41L4A | Q9HCS5 | 0.0068 | -0.56 | -1.48 |
| EPC1 | Q9H2F5 | 2.20E-05 | -0.95 | -1.93 |
| EPN2 | O95208 | 0.028 | -0.39 | -1.31 |
| EPRS | P07814 | 0.005 | -0.29 | -1.22 |
| ERBB3 | P21860 | 0.023 | -1.05 | -2.07 |
| ERICH1 | Q86X53 | 0.046 | -0.39 | -1.31 |
| ERVK-24 | P63145 | 2.20E-04 | -0.98 | -1.97 |
| ESF1 | Q9H501 | 0.0058 | -0.35 | -1.28 |
| ESPL1 | Q14674 | 0.024 | -0.33 | -1.26 |
| ESYT1 | Q9BSJ8 | 0.001 | -0.4 | -1.32 |
| ETV5 | P41161 | 0.026 | -0.42 | -1.33 |
| EXOSC5 | Q9NQT4 | 8.70E-06 | -0.65 | -1.57 |
| EXOSC7 | Q15024 | 0.011 | -0.36 | -1.29 |
| EXOSC9 | Q06265 | 8.40E-04 | -0.49 | -1.4 |
| FA63B_HUMAN | NA | 2.10E-04 | -0.61 | -1.53 |
| FAM118A | Q9NWS6 | 7.30E-05 | -2.16 | -4.46 |
| FAM124A | Q86V42 | 0.014 | -0.26 | -1.2 |
| FAM160A1 | Q05DH4 | 0.0035 | -0.62 | -1.53 |
| FAM189A2 | Q15884 | 0.0069 | -0.72 | -1.64 |
| FAM234A | Q9H0X4 | 0.046 | -0.42 | -1.34 |
| FAM83G | A6ND36 | 0.013 | -0.31 | -1.24 |
| FANCM | Q8IYD8 | 0.0019 | -0.31 | -1.24 |
| FARP1 | Q9Y4F1 | 0.038 | -0.31 | -1.24 |
| FEZ2 | Q9UHY8 | 9.50E-04 | -0.61 | -1.53 |
| FKBP5 | Q13451 | 0.021 | -0.36 | -1.29 |
| FLNB | O75369 | 4.10E-04 | -0.58 | -1.49 |
| FLVCR1 | Q9Y5Y0 | 7.60E-04 | -0.5 | -1.42 |
| FNBP4 | Q8N3X1 | 0.0054 | -0.39 | -1.31 |
| FNDC3B | Q53EP0 | 2.10E-04 | -0.74 | -1.67 |
| FOSL1 | P15407 | 0.028 | -0.61 | -1.52 |
| FOXK1 | P85037 | 4.30E-05 | -0.53 | -1.44 |
| FOXO4 | P98177 | 0.0081 | -0.59 | -1.51 |
| FSIP1 | Q8NA03 | 0.0087 | -0.37 | -1.29 |
| FTSJ1 | Q9UET6 | 0.0082 | -0.37 | -1.29 |
| FXR1 | P51114 | 3.70E-04 | -0.46 | -1.38 |
| FXYD7 | P58549 | 1.90E-05 | -1.16 | -2.23 |
| G3BP2 | Q9UN86 | 0.0015 | -0.34 | -1.27 |
| GAL | P22466 | 0.0021 | -0.55 | -1.46 |

| | | | | |
|---|---|---|---|---|
| GALNT7 | Q86SF2 | 0.015 | -0.63 | -1.55 |
| GAP43 | P17677 | 2.90E-05 | -0.94 | -1.92 |
| GAPDH | P04406 | 0.0032 | -0.45 | -1.36 |
| GEMIN8 | Q9NWZ8 | 0.031 | -0.57 | -1.48 |
| GLIS2 | Q9BZE0 | 0.005 | -0.34 | -1.26 |
| GLYR1 | Q49A26 | 0.042 | -0.44 | -1.36 |
| GMIP | Q9P107 | 0.016 | -0.32 | -1.25 |
| GMPR2 | Q9P2T1 | 9.30E-05 | -0.67 | -1.6 |
| GNL1 | P36915 | 0.0057 | -0.37 | -1.29 |
| GPATCH4 | Q5T3I0 | 0.002 | -0.89 | -1.86 |
| GPRIN1 | Q7Z2K8 | 4.80E-04 | -0.44 | -1.36 |
| GPS1 | Q13098 | 0.009 | -0.76 | -1.7 |
| GPSM1 | Q86YR5 | 6.80E-04 | -0.67 | -1.6 |
| GSG2 | Q8TF76 | 0.037 | -0.3 | -1.23 |
| GTF2A1 | P52655 | 0.017 | -0.59 | -1.5 |
| GTF3A | Q92664 | 2.50E-05 | -0.74 | -1.67 |
| GTF3C3 | Q9Y5Q9 | 7.90E-04 | -0.4 | -1.32 |
| GZF1 | Q9H116 | 0.02 | -0.3 | -1.23 |
| HADHA | P40939 | 0.034 | -0.42 | -1.33 |
| HDAC9 | Q9UKV0 | 0.0028 | -0.48 | -1.4 |
| HDLBP | Q00341 | 0.0032 | -0.36 | -1.28 |
| HEATR1 | Q9H583 | 0.011 | -0.46 | -1.38 |
| HECTD4 | Q9Y4D8 | 0.039 | -0.34 | -1.26 |
| HIC2 | Q96JB3 | 0.033 | -0.33 | -1.26 |
| HLA-A | P30443 | 5.80E-04 | -0.52 | -1.43 |
| HMGN3 | Q15651 | 0.016 | -0.43 | -1.35 |
| HMGN5 | P82970 | 0.0032 | -0.49 | -1.4 |
| HMOX1 | P09601 | 0.0019 | -0.72 | -1.65 |
| HNRNPAB | Q99729 | 0.025 | -0.62 | -1.54 |
| HNRNPC | P07910 | 0.0061 | -0.34 | -1.26 |
| HNRNPH3 | P31942 | 0.031 | -0.29 | -1.22 |
| HSP90B1 | P14625 | 8.40E-06 | -0.91 | -1.88 |
| HSPB1 | P04792 | 0.027 | -0.43 | -1.35 |
| HSPD1 | P10809 | 0.0059 | -0.34 | -1.27 |
| HTR1A | P08908 | 0.0022 | -0.79 | -1.73 |
| HYOU1 | Q9Y4L1 | 0.0036 | -1.07 | -2.1 |
| HYPK | Q9NX55 | 0.003 | -0.44 | -1.35 |
| ICE1 | Q9Y2F5 | 0.0027 | -0.3 | -1.23 |
| ICK | Q9UPZ9 | 0.029 | -0.28 | -1.21 |
| ID3 | Q02535 | 0.0014 | -1.06 | -2.09 |
| IFT43 | Q96FT9 | 0.0012 | -0.65 | -1.57 |

| | | | | |
|---|---|---|---|---|
| ILF2 | Q12905 | 0.0021 | -0.38 | -1.3 |
| IMPACT | Q9P2X3 | 0.013 | -0.26 | -1.2 |
| IMUP | Q9GZP8 | 0.013 | -0.64 | -1.56 |
| INCENP | Q9NQS7 | 0.0077 | -0.27 | -1.2 |
| ING5 | Q8WYH8 | 2.90E-04 | -0.53 | -1.44 |
| INO80E | Q8NBZ0 | 0.034 | -0.67 | -1.6 |
| IQGAP1 | P46940 | 0.027 | -0.26 | -1.2 |
| IQSEC2 | Q5JU85 | 5.70E-04 | -0.69 | -1.62 |
| ITGB5 | P18084 | 0.0036 | -0.59 | -1.5 |
| JADE3 | Q92613 | 0.0022 | -0.37 | -1.29 |
| JAM2 | P57087 | 0.03 | -0.33 | -1.26 |
| JMJD1C | Q15652 | 0.0068 | -0.26 | -1.2 |
| JPH3 | Q8WXH2 | 0.015 | -0.56 | -1.47 |
| JUND | P17535 | 0.0063 | -0.35 | -1.27 |
| KBTBD8 | Q8NFY9 | 0.015 | -0.4 | -1.31 |
| KDM2A | Q9Y2K7 | 0.0026 | -0.31 | -1.24 |
| KDM2B | Q8NHM5 | 1.80E-06 | -0.89 | -1.85 |
| KDM6A | O15550 | 2.90E-06 | -1.23 | -2.34 |
| KIAA1429 | Q69YN4 | 0.0082 | -0.29 | -1.22 |
| KIAA2026 | Q5HYC2 | 0.032 | -0.39 | -1.31 |
| KIF1C | O43896 | 0.02 | -0.26 | -1.2 |
| KIF21B | O75037 | 0.011 | -0.31 | -1.24 |
| KIF2C | Q99661 | 0.043 | -0.31 | -1.24 |
| KIRREL | Q96J84 | 9.30E-06 | -1.04 | -2.06 |
| KIZ | Q2M2Z5 | 0.0049 | -0.54 | -1.46 |
| KMT2A | Q03164 | 0.0068 | -0.34 | -1.26 |
| KMT2C | Q8NEZ4 | 0.038 | -0.35 | -1.27 |
| KMT2E | Q8IZD2 | 0.0056 | -0.33 | -1.25 |
| KNOP1 | Q1ED39 | 0.016 | -0.42 | -1.34 |
| KNSTRN | Q9Y448 | 0.05 | -0.54 | -1.46 |
| KPNA4 | O00629 | 0.048 | -0.4 | -1.32 |
| KRI1 | Q8N9T8 | 4.80E-04 | -0.43 | -1.34 |
| L1RE1 | Q9UN81 | 0.0011 | -0.49 | -1.4 |
| LAP3 | P28838 | 0.003 | -0.53 | -1.45 |
| LARP1B | Q659C4 | 0.017 | -0.26 | -1.2 |
| LARP4 | Q71RC2 | 0.0017 | -0.36 | -1.29 |
| LARS | Q9P2J5 | 0.0081 | -0.3 | -1.23 |
| LAS1L | Q9Y4W2 | 0.039 | -0.28 | -1.21 |
| LATS1 | O95835 | 0.0015 | -0.39 | -1.31 |
| LECT1_HUMAN | NA | 4.10E-06 | -1.65 | -3.13 |
| LIMA1 | Q9UHB6 | 0.0017 | -0.39 | -1.31 |

| | | | | |
|---|---|---|---|---|
| LMNA | P02545 | 0.0014 | -0.45 | -1.37 |
| LNPEP | Q9UIQ6 | 5.80E-04 | -0.62 | -1.54 |
| LPIN1 | Q14693 | 0.016 | -0.89 | -1.86 |
| LTBP1 | Q14766 | 0.0075 | -0.32 | -1.25 |
| LTV1 | Q96GA3 | 0.0018 | -2.01 | -4.02 |
| LYN | P07948 | 9.70E-04 | -0.46 | -1.38 |
| LYSMD1 | Q96S90 | 0.015 | -0.41 | -1.33 |
| LYSMD3 | Q7Z3D4 | 5.90E-05 | -1.08 | -2.12 |
| MAGEL2 | Q9UJ55 | 0.0099 | -0.34 | -1.27 |
| MAP1A | P78559 | 0.0041 | -0.41 | -1.32 |
| MARS | P56192 | 0.0064 | -0.48 | -1.4 |
| MATK | P42679 | 0.0012 | -0.5 | -1.41 |
| MATR3 | P43243 | 0.041 | -0.26 | -1.2 |
| MCF2L | O15068 | 0.014 | -0.28 | -1.21 |
| MDN1 | Q9NU22 | 0.03 | -0.62 | -1.54 |
| MECP2 | P51608 | 0.011 | -0.28 | -1.21 |
| MED22 | Q15528 | 6.40E-04 | -0.54 | -1.46 |
| MEF2D | Q14814 | 0.005 | -0.61 | -1.53 |
| METTL14 | Q9HCE5 | 0.038 | -0.45 | -1.37 |
| MFAP1 | P55081 | 0.013 | -0.26 | -1.2 |
| MFSD6 | Q6ZSS7 | 0.0094 | -0.4 | -1.32 |
| MGME1 | Q9BQP7 | 6.30E-04 | -0.59 | -1.5 |
| MIA3 | Q5JRA6 | 0.012 | -0.33 | -1.26 |
| MICAL1 | Q8TDZ2 | 0.0024 | -0.57 | -1.49 |
| MIEF1 | Q9NQG6 | 2.20E-04 | -0.46 | -1.37 |
| MIER2 | Q8N344 | 9.50E-04 | -0.36 | -1.28 |
| MMP15 | P51511 | 3.90E-05 | -0.7 | -1.62 |
| MNAT1 | P51948 | 0.028 | -0.36 | -1.28 |
| MON2 | Q7Z3U7 | 0.029 | -0.28 | -1.21 |
| MPHOSPH10 | O00566 | 2.70E-04 | -0.48 | -1.39 |
| MPHOSPH8 | Q99549 | 0.0068 | -0.29 | -1.22 |
| MREG | Q8N565 | 0.0013 | -0.48 | -1.39 |
| MRPS15 | P82914 | 0.0027 | -0.7 | -1.62 |
| MRPS18B | Q9Y676 | 0.015 | -0.43 | -1.35 |
| MSL1 | Q68DK7 | 0.018 | -0.5 | -1.41 |
| MTMR10 | Q9NXD2 | 0.0014 | -0.46 | -1.37 |
| MTRR | Q9UBK8 | 0.045 | -0.27 | -1.2 |
| MUM1 | Q2TAK8 | 2.70E-04 | -0.64 | -1.56 |
| MYC | P01106 | 0.0081 | -0.38 | -1.3 |
| MYSM1 | Q5VVJ2 | 0.0065 | -0.41 | -1.33 |
| NAA38 | Q9BRA0 | 0.005 | -1.39 | -2.61 |

| | | | | |
|---|---|---|---|---|
| NACAD | O15069 | 5.00E-05 | -0.64 | -1.56 |
| NAF1 | Q96HR8 | 1.70E-05 | -1.93 | -3.82 |
| NAGK | Q9UJ70 | 4.90E-04 | -0.43 | -1.34 |
| NAP1L2 | Q9ULW6 | 0.0063 | -0.3 | -1.23 |
| NAT10 | Q9H0A0 | 0.0014 | -0.5 | -1.41 |
| NAV3 | Q8IVL0 | 3.10E-06 | -0.9 | -1.87 |
| NCAPD3 | P42695 | 0.03 | -0.36 | -1.28 |
| NDEL1 | Q9GZM8 | 0.0027 | -0.47 | -1.38 |
| NDUFB4 | O95168 | 0.0099 | -0.35 | -1.28 |
| NECTIN1 | Q15223 | 0.049 | -0.27 | -1.21 |
| NECTIN3 | Q9NQS3 | 5.00E-04 | -0.54 | -1.45 |
| NEFL | P07196 | 1.90E-06 | -2.08 | -4.22 |
| NEFM | P07197 | 2.30E-06 | -1.44 | -2.72 |
| NELFB | Q8WX92 | 0.034 | -0.3 | -1.23 |
| NEMF | O60524 | 0.021 | -0.39 | -1.31 |
| NES | P48681 | 1.10E-05 | -0.76 | -1.69 |
| NEXN | Q0ZGT2 | 0.0027 | -0.54 | -1.45 |
| NGFR | P08138 | 0.02 | -0.5 | -1.41 |
| NIFK | Q9BYG3 | 0.0046 | -0.38 | -1.3 |
| NOC2L | Q9Y3T9 | 0.015 | -0.35 | -1.28 |
| NOL4 | O94818 | 0.018 | -0.29 | -1.22 |
| NOL8 | Q76FK4 | 7.40E-04 | -0.4 | -1.32 |
| NOM1 | Q5C9Z4 | 0.0058 | -0.37 | -1.3 |
| NOMO1 | Q15155 | 0.0019 | -0.38 | -1.3 |
| NOMO2 | Q5JPE7 | 0.0044 | -0.32 | -1.25 |
| NOP10 | Q9NPE3 | 5.20E-05 | -1.15 | -2.22 |
| NOP2 | P46087 | 0.014 | -0.32 | -1.25 |
| NOP56 | O00567 | 0.0027 | -0.65 | -1.57 |
| NOP58 | Q9Y2X3 | 0.037 | -0.56 | -1.47 |
| NRBP2 | Q9NSY0 | 2.30E-04 | -0.49 | -1.41 |
| NRK | Q7Z2Y5 | 9.70E-06 | -1.46 | -2.75 |
| NSD1 | Q96L73 | 2.70E-04 | -0.47 | -1.38 |
| NT5C | Q8TCD5 | 0.0059 | -0.59 | -1.5 |
| NUCB1 | Q02818 | 2.40E-05 | -1.25 | -2.38 |
| NUMBL | Q9Y6R0 | 0.0046 | -0.38 | -1.3 |
| NUP160 | Q12769 | 0.0015 | -0.35 | -1.28 |
| NUPL2 | O15504 | 0.016 | -0.3 | -1.23 |
| NYNRIN | Q9P2P1 | 9.20E-04 | -0.56 | -1.47 |
| OBSL1 | O75147 | 7.60E-04 | -0.35 | -1.27 |
| OCIAD1 | Q9NX40 | 0.0077 | -0.4 | -1.32 |
| OSTF1 | Q92882 | 2.90E-04 | -0.78 | -1.72 |

| | | | | |
|---|---|---|---|---|
| PAIP2B | Q9ULR5 | 0.012 | -0.28 | -1.21 |
| PAK1 | Q13153 | 0.005 | -0.33 | -1.26 |
| PAK2 | Q13177 | 0.029 | -0.29 | -1.22 |
| PALM2 | Q8IXS6 | 3.10E-04 | -1 | -2 |
| PAM | P19021 | 0.0011 | -1 | -2 |
| PAPD5 | Q8NDF8 | 0.012 | -1.08 | -2.11 |
| PARD3B | Q8TEW8 | 0.0079 | -0.32 | -1.25 |
| PARG | Q86W56 | 0.036 | -0.36 | -1.28 |
| PARP8 | Q8N3A8 | 0.016 | -0.4 | -1.32 |
| PCDH7 | O60245 | 0.049 | -0.28 | -1.21 |
| PCP4L1 | A6NKN8 | 0.014 | -0.5 | -1.41 |
| PCYT1A | P49585 | 0.026 | -0.34 | -1.27 |
| PDCD2L | Q9BRP1 | 0.0059 | -0.64 | -1.56 |
| PDE3A | Q14432 | 0.0025 | -0.44 | -1.36 |
| PEA15 | Q15121 | 0.001 | -1.1 | -2.14 |
| PFDN4 | Q9NQP4 | 0.027 | -0.55 | -1.47 |
| PFKP | Q01813 | 6.40E-04 | -0.41 | -1.33 |
| PGAM1 | P18669 | 0.01 | -0.3 | -1.23 |
| PGM1 | P36871 | 0.041 | -0.38 | -1.3 |
| PGM2L1 | Q6PCE3 | 0.0037 | -0.49 | -1.4 |
| PGRMC2 | O15173 | 2.70E-04 | -0.42 | -1.34 |
| PHAX | Q9H814 | 0.025 | -0.39 | -1.31 |
| PHF20 | Q9BVI0 | 0.0064 | -0.3 | -1.23 |
| PHF5A | Q7RTV0 | 0.043 | -0.26 | -1.2 |
| PHGDH | O43175 | 0.012 | -0.42 | -1.34 |
| PICALM | Q13492 | 0.004 | -0.78 | -1.72 |
| PITPNM1 | O00562 | 4.70E-05 | -0.55 | -1.46 |
| PKP1 | Q13835 | 8.60E-06 | -0.81 | -1.75 |
| PLA2G4A | P47712 | 0.0085 | -0.54 | -1.45 |
| PLAGL2 | Q9UPG8 | 2.50E-04 | -0.41 | -1.33 |
| PLCB4 | Q15147 | 0.018 | -0.39 | -1.31 |
| PLD2 | O14939 | 4.40E-04 | -0.51 | -1.42 |
| PMAIP1 | Q13794 | 6.30E-04 | -0.52 | -1.43 |
| PNKD | Q8N490 | 0.04 | -0.44 | -1.36 |
| PNKP | Q96T60 | 0.0066 | -0.34 | -1.27 |
| POLDIP2 | Q9Y2S7 | 0.016 | -0.26 | -1.2 |
| POLR2C | P19387 | 0.0017 | -0.36 | -1.28 |
| POLR3G | O15318 | 0.0059 | -1.17 | -2.24 |
| POP1 | Q99575 | 4.60E-04 | -0.62 | -1.53 |
| PPAN | Q9NQ55 | 0.024 | -0.56 | -1.48 |
| PPIL4 | Q8WUA2 | 0.0039 | -0.28 | -1.22 |

| | | | | |
|---|---|---|---|---|
| PPP1R12B | O60237 | 0.023 | -0.27 | -1.21 |
| PPP1R14A | Q96A00 | 0.01 | -0.36 | -1.28 |
| PPP1R18 | Q6NYC8 | 0.013 | -0.31 | -1.24 |
| PPP2R5D | Q14738 | 0.0055 | -0.47 | -1.38 |
| PPP2R5E | Q16537 | 0.0091 | -0.43 | -1.35 |
| PPP3CA | Q08209 | 0.0018 | -1.06 | -2.08 |
| PPP3CB | P16298 | 0.0029 | -0.88 | -1.85 |
| PPP4R2 | Q9NY27 | 0.0085 | -0.38 | -1.3 |
| PRDM2 | Q13029 | 0.023 | -0.3 | -1.23 |
| PRKAR1B | P31321 | 0.047 | -0.35 | -1.28 |
| PRKAR2B | P31323 | 0.0021 | -0.51 | -1.42 |
| PRKCG | P05129 | 0.003 | -0.49 | -1.41 |
| PRKCH | P24723 | 0.0069 | -0.28 | -1.22 |
| PRKG1 | Q13976 | 3.70E-05 | -1.08 | -2.12 |
| PRPF38A | Q8NAV1 | 1.10E-04 | -0.85 | -1.8 |
| PRPF4 | O43172 | 0.0091 | -0.43 | -1.34 |
| PRPF6 | O94906 | 0.0017 | -0.56 | -1.47 |
| PSMA5 | P28066 | 0.012 | -0.55 | -1.47 |
| PTK7 | Q13308 | 8.60E-04 | -0.57 | -1.48 |
| PTPRZ1 | P23471 | 0.006 | -0.36 | -1.28 |
| PTRF_HUMAN | NA | 0.002 | -0.62 | -1.54 |
| PTS | Q03393 | 0.002 | -0.58 | -1.49 |
| PUS7L | Q9H0K6 | 0.006 | -0.47 | -1.38 |
| PWWP2A | Q96N64 | 6.70E-05 | -0.63 | -1.55 |
| QSER1 | Q2KHR3 | 0.0092 | -0.36 | -1.28 |
| QSOX2 | Q6ZRP7 | 0.0027 | -0.6 | -1.52 |
| RAB13 | P51153 | 0.007 | -0.46 | -1.37 |
| RAB2B | Q8WUD1 | 6.30E-04 | -0.7 | -1.62 |
| RAD23B | P54727 | 0.0012 | -0.68 | -1.61 |
| RALY | Q9UKM9 | 0.023 | -0.26 | -1.2 |
| RANGAP1 | P46060 | 0.0027 | -0.41 | -1.33 |
| RAP1GAP2 | Q684P5 | 0.003 | -0.49 | -1.4 |
| RASA3 | Q14644 | 0.002 | -0.71 | -1.63 |
| RASL11A | Q6T310 | 0.042 | -0.33 | -1.26 |
| RBM5 | P52756 | 0.043 | -0.45 | -1.37 |
| RBM8A | Q9Y5S9 | 0.0011 | -0.61 | -1.53 |
| RBMXL1 | Q96E39 | 0.0038 | -0.31 | -1.24 |
| RCOR1 | Q9UKL0 | 0.0044 | -0.43 | -1.34 |
| RCOR2 | Q8IZ40 | 0.019 | -0.29 | -1.22 |
| REEP2 | Q9BRK0 | 0.018 | -0.32 | -1.25 |
| REEP3 | Q6NUK4 | 2.00E-04 | -0.58 | -1.49 |

| | | | | |
|---|---|---|---|---|
| RFC2 | P35250 | 0.021 | -0.26 | -1.2 |
| RIC8B | Q9NVN3 | 0.0031 | -1.57 | -2.98 |
| RIOK2 | Q9BVS4 | 0.013 | -0.43 | -1.35 |
| RLIM | Q9NVW2 | 0.0035 | -0.35 | -1.27 |
| RNF113A | O15541 | 2.60E-04 | -0.45 | -1.36 |
| RNF181 | Q9P0P0 | 0.0085 | -0.39 | -1.31 |
| RNF20 | Q5VTR2 | 0.0018 | -0.36 | -1.29 |
| ROR2 | Q01974 | 7.80E-05 | -1.17 | -2.25 |
| RPL17 | P18621 | 0.0024 | -0.43 | -1.35 |
| RPL30 | P62888 | 0.0054 | -0.3 | -1.23 |
| RPL5 | P46777 | 1.70E-04 | -0.59 | -1.51 |
| RPP25L | Q8N5L8 | 0.012 | -0.35 | -1.28 |
| RPRD1A | Q96P16 | 0.003 | -0.34 | -1.27 |
| RPS17 | P08708 | 2.30E-05 | -0.68 | -1.61 |
| RPS18 | P62269 | 0.015 | -0.37 | -1.29 |
| RPS2 | P15880 | 0.0029 | -0.45 | -1.37 |
| RPS26 | P62854 | 0.011 | -0.48 | -1.39 |
| RPS27 | P42677 | 0.0031 | -0.37 | -1.3 |
| RPS6 | P62753 | 0.0019 | -1.06 | -2.09 |
| RPS7 | P62081 | 0.016 | -0.75 | -1.68 |
| RPSA | P08865 | 2.50E-04 | -0.46 | -1.38 |
| RRP12 | Q5JTH9 | 0.012 | -0.28 | -1.21 |
| RRP1B | Q14684 | 0.0027 | -0.4 | -1.32 |
| RRP36 | Q96EU6 | 0.0048 | -0.42 | -1.34 |
| RRP7A | Q9Y3A4 | 0.015 | -0.48 | -1.39 |
| RSF1 | Q96T23 | 0.0017 | -0.33 | -1.26 |
| RSL1D1 | O76021 | 0.0017 | -0.52 | -1.44 |
| RTCB | Q9Y3I0 | 0.0076 | -0.31 | -1.24 |
| RTN1 | Q16799 | 0.0014 | -0.59 | -1.51 |
| RTN3 | O95197 | 0.019 | -0.34 | -1.27 |
| RUFY2 | Q8WXA3 | 2.50E-04 | -0.77 | -1.71 |
| RUNDC1 | Q96C34 | 0.003 | -0.48 | -1.39 |
| RUNX1T1 | Q06455 | 0.0025 | -0.39 | -1.31 |
| SART1 | O43290 | 0.009 | -0.26 | -1.2 |
| SART3 | Q15020 | 0.0063 | -0.37 | -1.29 |
| SATB1 | Q01826 | 0.048 | -0.51 | -1.43 |
| SATB2 | Q9UPW6 | 0.0026 | -0.52 | -1.43 |
| SCAF1 | Q9H7N4 | 0.009 | -0.27 | -1.21 |
| SCAP | Q12770 | 1.00E-05 | -1.41 | -2.65 |
| SCNM1 | Q9BWG6 | 0.0049 | -0.49 | -1.4 |
| SDAD1 | Q9NVU7 | 0.0017 | -0.64 | -1.56 |

| | | | | |
|---|---|---|---|---|
| SDC2 | P34741 | 0.047 | -0.61 | -1.53 |
| SEC14L1 | Q92503 | 0.0029 | -0.5 | -1.41 |
| SEC24D | O94855 | 0.033 | -0.27 | -1.2 |
| SEC61B | P60468 | 0.0044 | -0.5 | -1.41 |
| SEMA4A | Q9H3S1 | 0.012 | -0.38 | -1.3 |
| SEMA4D | Q92854 | 0.0045 | -0.92 | -1.89 |
| SENP6 | Q9GZR1 | 0.014 | -0.27 | -1.21 |
| 2-Sep | Q15019 | 0.004 | -0.52 | -1.43 |
| SERINC3 | Q13530 | 0.011 | -0.26 | -1.2 |
| SERTM1 | A2A2V5 | 0.022 | -0.37 | -1.3 |
| SF1 | Q15637 | 0.043 | -0.37 | -1.3 |
| SF3A1 | Q15459 | 0.0012 | -0.46 | -1.37 |
| SGO2 | Q562F6 | 0.013 | -0.44 | -1.36 |
| SHARPIN | Q9H0F6 | 0.0075 | -0.26 | -1.2 |
| SHCBP1 | Q8NEM2 | 0.011 | -0.39 | -1.31 |
| SIPA1L2 | Q9P2F8 | 1.10E-04 | -0.49 | -1.4 |
| SIRT6 | Q8N6T7 | 0.0097 | -0.35 | -1.27 |
| SIT1 | Q9Y3P8 | 0.0065 | -0.44 | -1.35 |
| SKP2 | Q13309 | 0.015 | -0.3 | -1.23 |
| SLAIN1 | Q8ND83 | 2.70E-05 | -0.7 | -1.62 |
| SLC16A10 | Q8TF71 | 2.60E-04 | -0.72 | -1.64 |
| SLC19A3 | Q9BZV2 | 0.032 | -0.31 | -1.24 |
| SLC1A5 | Q15758 | 0.016 | -0.34 | -1.26 |
| SLC20A2 | Q08357 | 0.0028 | -0.4 | -1.32 |
| SLC29A1 | Q99808 | 0.0059 | -0.37 | -1.29 |
| SLC29A2 | Q14542 | 0.013 | -0.36 | -1.28 |
| SLC2A1 | P11166 | 2.40E-04 | -0.62 | -1.54 |
| SLC2A3 | P11169 | 0.036 | -0.28 | -1.22 |
| SLC30A1 | Q9Y6M5 | 0.012 | -0.53 | -1.45 |
| SLC35A5 | Q9BS91 | 0.0034 | -0.49 | -1.4 |
| SLC39A14 | Q15043 | 0.0017 | -0.41 | -1.33 |
| SLC45A4 | Q5BKX6 | 0.0013 | -0.35 | -1.27 |
| SLC4A8 | Q2Y0W8 | 0.015 | -0.43 | -1.34 |
| SLIRP | Q9GZT3 | 0.003 | -0.4 | -1.32 |
| SLU7 | O95391 | 0.019 | -0.62 | -1.54 |
| SMAP | O00193 | 5.20E-04 | -0.56 | -1.47 |
| SMARCA2 | P51531 | 8.40E-04 | -0.72 | -1.64 |
| SNAI1 | O95863 | 0.012 | -0.43 | -1.35 |
| SNAP23 | O00161 | 0.034 | -0.28 | -1.22 |
| SND1 | Q7KZF4 | 3.20E-05 | -0.84 | -1.79 |
| SNRPC | P09234 | 0.0047 | -0.79 | -1.72 |

| | | | | |
|---|---|---|---|---|
| SNRPD3 | P62318 | 4.60E-06 | -0.81 | -1.76 |
| SNTA1 | Q13424 | 0.0088 | -0.5 | -1.42 |
| SNX1 | Q13596 | 0.037 | -0.28 | -1.21 |
| SNX3 | O60493 | 0.022 | -0.3 | -1.23 |
| SORT1 | Q99523 | 0.02 | -0.39 | -1.31 |
| SOX13 | Q9UN79 | 0.012 | -0.28 | -1.21 |
| SOX15 | O60248 | 1.30E-05 | -1.4 | -2.65 |
| SOX4 | Q06945 | 0.0056 | -0.38 | -1.3 |
| SPATA2 | Q9UM82 | 5.60E-04 | -0.49 | -1.41 |
| SPG20 | Q8N0X7 | 3.20E-04 | -0.58 | -1.49 |
| SPP1 | P10451 | 0.0094 | -0.66 | -1.58 |
| SPTAN1 | Q13813 | 7.90E-05 | -0.61 | -1.53 |
| SRFBP1 | Q8NEF9 | 0.015 | -1.14 | -2.21 |
| SRP72 | O76094 | 0.0039 | -0.3 | -1.23 |
| SRSF1 | Q07955 | 5.20E-05 | -0.63 | -1.55 |
| SRSF6 | Q13247 | 0.0099 | -0.28 | -1.21 |
| STAU1 | O95793 | 0.0012 | -0.42 | -1.34 |
| STIP1 | P31948 | 0.043 | -0.68 | -1.61 |
| STK25 | O00506 | 0.0085 | -0.43 | -1.35 |
| STK26 | Q9P289 | 0.014 | -0.44 | -1.36 |
| STK33 | Q9BYT3 | 0.0037 | -0.36 | -1.28 |
| STK38L | Q9Y2H1 | 0.04 | -0.34 | -1.26 |
| STRIP1 | Q5VSL9 | 0.003 | -0.66 | -1.58 |
| STRN | O43815 | 0.014 | -0.36 | -1.28 |
| STRN3 | Q13033 | 0.0095 | -0.37 | -1.29 |
| STUB1 | Q9UNE7 | 0.043 | -0.36 | -1.28 |
| STX10 | O60499 | 0.0026 | -0.35 | -1.28 |
| SUMO2 | P61956 | 0.024 | -0.3 | -1.23 |
| SUPT7L | O94864 | 0.011 | -0.37 | -1.29 |
| SUV39H2 | Q9H5I1 | 8.40E-05 | -0.85 | -1.81 |
| SUZ12 | Q15022 | 0.011 | -0.31 | -1.24 |
| SYAP1 | Q96A49 | 1.40E-04 | -0.56 | -1.48 |
| SYDE2 | Q5VT97 | 1.40E-04 | -0.58 | -1.49 |
| SYNCRIP | O60506 | 0.0013 | -0.38 | -1.3 |
| SZRD1 | Q7Z422 | 0.0069 | -0.26 | -1.2 |
| TAF1 | P21675 | 0.013 | -0.26 | -1.2 |
| TAOK2 | Q9UL54 | 0.0036 | -0.77 | -1.71 |
| TBC1D10A | Q9BXI6 | 0.013 | -0.54 | -1.46 |
| TBC1D16 | Q8TBP0 | 0.036 | -0.34 | -1.26 |
| TBCB | Q99426 | 5.40E-06 | -0.9 | -1.87 |
| TBX18 | O95935 | 0.0073 | -0.39 | -1.31 |

| | | | | |
|---|---|---|---|---|
| TCEAL2 | Q9H3H9 | 8.90E-06 | -2.22 | -4.65 |
| TET1 | Q8NFU7 | 0.0045 | -0.38 | -1.3 |
| TEX2 | Q8IWB9 | 0.0083 | -0.38 | -1.3 |
| TFAM | Q00059 | 9.10E-04 | -0.51 | -1.43 |
| TFAP2C | Q92754 | 0.0013 | -0.59 | -1.5 |
| TGFB1I1 | O43294 | 0.045 | -0.29 | -1.22 |
| THOC5 | Q13769 | 1.00E-04 | -0.59 | -1.5 |
| THUMPD1 | Q9NXG2 | 0.04 | -0.27 | -1.21 |
| TLN1 | Q9Y490 | 0.042 | -0.27 | -1.21 |
| TMCO1 | Q9UM00 | 0.019 | -0.26 | -1.2 |
| TMEM184B | Q9Y519 | 0.0081 | -0.27 | -1.21 |
| TMEM200A | Q86VY9 | 0.036 | -0.31 | -1.24 |
| TMX2 | Q9Y320 | 5.50E-06 | -0.84 | -1.79 |
| TMX4 | Q9H1E5 | 0.001 | -2.25 | -4.75 |
| TNFRSF8 | P28908 | 0.015 | -0.41 | -1.33 |
| TPBG | Q13641 | 0.026 | -0.35 | -1.28 |
| TPM1 | P09493 | 0.0068 | -0.61 | -1.52 |
| TPMT | P51580 | 0.019 | -0.27 | -1.2 |
| TPRN | Q4KMQ1 | 0.0029 | -0.37 | -1.29 |
| TRAFD1 | O14545 | 0.031 | -0.26 | -1.2 |
| TRAPPC11 | Q7Z392 | 0.028 | -0.67 | -1.59 |
| TRAPPC8 | Q9Y2L5 | 0.033 | -0.53 | -1.45 |
| TRIM2 | Q9C040 | 0.0025 | -0.4 | -1.32 |
| TRIM3 | O75382 | 0.021 | -0.26 | -1.2 |
| TRIP10 | Q15642 | 9.80E-05 | -0.7 | -1.62 |
| TRMT1 | Q9NXH9 | 0.018 | -0.26 | -1.2 |
| TSC22D1 | Q15714 | 0.0068 | -0.35 | -1.27 |
| TSC22D4 | Q9Y3Q8 | 0.007 | -0.34 | -1.26 |
| TTC1 | Q99614 | 3.90E-04 | -0.4 | -1.32 |
| TUBA1B | P68363 | 0.01 | -0.74 | -1.67 |
| TUBA1C | Q9BQE3 | 0.0093 | -1.1 | -2.15 |
| TUBA4A | P68366 | 0.0019 | -0.89 | -1.85 |
| TXLNA | P40222 | 6.30E-04 | -0.77 | -1.71 |
| TXNDC9 | O14530 | 0.0093 | -0.31 | -1.24 |
| U2AF2 | P26368 | 0.011 | -0.29 | -1.22 |
| UBR7 | Q8N806 | 0.023 | -0.33 | -1.26 |
| UCHL3 | P15374 | 0.0025 | -0.32 | -1.25 |
| UGDH | O60701 | 0.033 | -0.53 | -1.45 |
| UNC13B | O14795 | 0.037 | -0.3 | -1.23 |
| UPF3A | Q9H1J1 | 0.045 | -0.98 | -1.98 |
| USP15 | Q9Y4E8 | 0.0031 | -0.33 | -1.26 |

| | | | | |
|---|---|---|---|---|
| USP16 | Q9Y5T5 | 0.0024 | -0.34 | -1.27 |
| USP33 | Q8TEY7 | 0.0023 | -0.45 | -1.37 |
| UTF1 | Q5T230 | 0.001 | -0.65 | -1.56 |
| UTP11 | Q9Y3A2 | 0.0087 | -0.69 | -1.61 |
| UTP18 | Q9Y5J1 | 0.0096 | -0.27 | -1.21 |
| UTP20 | O75691 | 0.031 | -0.28 | -1.22 |
| UTP3 | Q9NQZ2 | 0.001 | -0.39 | -1.31 |
| UTRN | P46939 | 4.20E-04 | -0.92 | -1.89 |
| VAMP2 | P63027 | 0.0012 | -0.98 | -1.97 |
| VAPA | Q9P0L0 | 0.025 | -0.48 | -1.39 |
| VDAC2 | P45880 | 1.20E-05 | -0.71 | -1.63 |
| WASH7_HUMAN | NA | 0.011 | -0.28 | -1.21 |
| WDR12 | Q9GZL7 | 5.60E-04 | -0.38 | -1.3 |
| WDR3 | Q9UNX4 | 8.90E-05 | -0.79 | -1.73 |
| WDR43 | Q15061 | 0.012 | -0.96 | -1.94 |
| WDR44 | Q5JSH3 | 8.90E-04 | -0.38 | -1.3 |
| WDR46 | O15213 | 0.0089 | -0.35 | -1.28 |
| WDR48 | Q8TAF3 | 0.018 | -0.34 | -1.26 |
| WDR70 | Q9NW82 | 5.00E-04 | -0.46 | -1.37 |
| WFS1 | O76024 | 0.034 | -0.5 | -1.41 |
| XPC | Q01831 | 0.0029 | -0.93 | -1.9 |
| XPO6 | Q96QU8 | 0.014 | -0.37 | -1.29 |
| XPR1 | Q9UBH6 | 0.021 | -0.28 | -1.21 |
| YBX3 | P16989 | 0.017 | -0.3 | -1.23 |
| YJ005_HUMAN | Q6ZSR9 | 0.0026 | -0.36 | -1.28 |
| YTHDC1 | Q96MU7 | 0.0025 | -0.38 | -1.3 |
| ZBTB10 | Q96DT7 | 2.90E-04 | -0.5 | -1.42 |
| ZBTB7A | O95365 | 0.0022 | -0.86 | -1.81 |
| ZC3H8 | Q8N5P1 | 2.10E-04 | -1.3 | -2.45 |
| ZCCHC3 | Q9NUD5 | 0.007 | -0.93 | -1.91 |
| ZCCHC7 | Q8N3Z6 | 0.003 | -0.56 | -1.47 |
| ZCRB1 | Q8TBF4 | 8.00E-04 | -0.38 | -1.3 |
| ZNF121 | P58317 | 0.0012 | -0.41 | -1.33 |
| ZNF184 | Q99676 | 0.0024 | -0.42 | -1.34 |
| ZNF207 | O43670 | 0.018 | -0.93 | -1.91 |
| ZNF264 | O43296 | 0.0046 | -0.44 | -1.35 |
| ZNF316 | A6NFI3 | 0.0062 | -0.39 | -1.31 |
| ZNF397 | Q8NF99 | 0.033 | -0.45 | -1.36 |
| ZNF462 | Q96JM2 | 0.012 | -0.3 | -1.23 |
| ZNF592 | Q92610 | 0.014 | -0.29 | -1.22 |
| ZNF641 | Q96N77 | 0.013 | -0.45 | -1.36 |

| | | | | |
|---|---|---|---|---|
| ZNF644 | Q9H582 | 0.021 | -0.3 | -1.23 |
| ZNF660 | Q6AZW8 | 0.0034 | -0.59 | -1.51 |
| ZNF697 | Q5TEC3 | 0.0022 | -0.43 | -1.35 |
| ZNF792 | Q3KQV3 | 0.0072 | -1.03 | -2.05 |
| ZNF84 | P51523 | 0.0038 | -0.6 | -1.51 |
| ZNHIT2 | Q9UHR6 | 2.70E-04 | -0.64 | -1.56 |
| ZNHIT3 | Q15649 | 3.40E-04 | -0.95 | -1.93 |
| ZRSR2 | Q15696 | 5.90E-04 | -0.54 | -1.46 |
| ZSCAN10 | Q96SZ4 | 2.90E-04 | -0.42 | -1.34 |
| ZSCAN21 | Q9Y5A6 | 0.014 | -0.34 | -1.27 |
| ZWILCH | Q9H900 | 0.021 | -0.27 | -1.21 |

**Supplementary Table 4:** Relative protein expression of select germ layer protein markers from nano LC/MS data

| GERM LAYER MARKER | FOLD CHANGE | P-VALUE |
|---|---|---|
| **Surface Ectoderm** | | |
| KRT8 | 1.11 | 7.50E-02 |
| KRT18 | 1.12 | 6.00E-02 |
| GRHL2 | 1.15 | 4.70E-01 |
| **Neuro Ectoderm** | | |
| MAP2 | -1.19 | 1.70E-02 |
| NES | -1.66 | 2.90E-06 |
| PAX6 | 2.4 | 4.40E-05 |
| SOX11 | -1 | 9.20E-01 |
| OTX2 | 1.49 | 5.70E-03 |
| **Pluripotency** | | |
| POU5F1 (OCT4) | -1.04 | 3.30E-01 |
| SOX2 | -1.04 | 3.20E-01 |
| DNMT3B | -1.5 | 2.00E-02 |
| SALL4 | -1.06 | 8.50E-02 |
| NANOG | -1.4 | 3.80E-02 |
| ESRRB | -1.04 | 6.10E-01 |
| **Early Mesodermal Marker** | | |
| None | | |
| **Primitive Endodermal Marker** | | |
| GDF1 | -1.67 | 1.10E-03 |
| GDF3 | -1.76 | 1.10E-05 |
| SALL4 | -1.06 | 1.10E-05 |
